# Reliability and Stability Challenges in ABCD Task fMRI Data

**DOI:** 10.1101/2021.10.08.463750

**Authors:** James T. Kennedy, Michael P. Harms, Ozlem Korucuoglu, Serguei V. Astafiev, Deanna M. Barch, Wesley K. Thompson, James M. Bjork, Andrey P. Anokhin

## Abstract

Trait stability of measures is an essential requirement for individual differences research. Functional MRI has been increasingly used in studies that rely on the assumption of trait stability, such as attempts to relate task related brain activation to individual differences in behavior and psychopathology. However, recent research using adult samples has questioned the trait stability of task-fMRI measures, as assessed by test-retest correlations. To date, little is known about trait stability of task fMRI in children. Here, we examined within-session reliability and longitudinal stability of task-fMRI using data from the Adolescent Brain Cognitive Development (ABCD) Study using its tasks focused on reward processing, response inhibition, and working memory. We also evaluated the effects of factors potentially affecting reliability and stability. Reliability and stability [quantified via an intraclass correlation (ICC) that focuses on rank consistency] was poor in virtually all brain regions, with an average ICC of .078 and .054 for short (within-session) and long-term (between-session) ICCs, respectively, in regions of interest (ROIs) historically-recruited by the tasks. ICC values in ROIs did not exceed the ‘poor’ cut-off of .4, and in fact rarely exceeded .2 (only 5.9%). Motion had a pronounced effect on estimated ICCs, with the lowest motion quartile of participants having a mean reliability/stability three times higher (albeit still ‘poor’) than the highest motion quartile. Regions with stronger activation tended to show higher ICCs, with the absolute value of activity and reliability/stability correlating at .53. Across regions, the magnitude of age-related longitudinal (between-session) changes positively correlated with the longitudinal stability of individual differences, which suggests developmental change was not necessarily responsible for poor stability. Poor reliability and stability of task-fMRI, particularly in children, diminishes potential utility of fMRI data due to a drastic reduction of effect sizes and, consequently, statistical power for the detection of brain-behavior associations. This essential issue needs to be addressed through optimization of preprocessing pipelines and data denoising methods.

## Introduction

Task-based functional magnetic resonance imaging (fMRI) has become a leading methodological approach in cognitive neuroscience. While initial application of fMRI focused on group-level effects such as average differences in regional brain activation between different stimuli, more recently fMRI has been increasingly applied to individual differences research such as across-subject correlation between task-related brain activation and other variables such as genetic markers, behavioral and cognitive performance, psychological traits, and psychopathology. Much of this research critically relies on the assumption that the magnitude of task-related regional activation is a stable trait-like measure, with individual differences between subjects prevailing over within-subject fluctuations between testing occasions, which is often quantified by test-retest reliability (intraclass correlation).^1^

However, recent studies have shown generally poor test-retest reliability of task-fMRI measures (Elliott et al., 2020; Herting, et al., 2018; Noble et al., 2021). Importantly, reproducibility of group-averaged patterns of activation can still be high despite poor stability of intra-individual differences in the magnitude of activation (Chaarani et al., 2021; Herting et al., 2018), since averaging reduces error variance, as prescribed by basic statistical theory. In the most representative study to date, Elliott et al. (2020) performed a meta-analysis of 56 test-retest reliability studies using various sensory, motor, and cognitive tasks, finding an average reliability of .397. Task specific average reliability [limiting to studies that reported all reliabilities calculated, though most were region of interest (ROI), only and not whole brain] ranged from a low of -.02 for an implicit memory encoding task (Brandt et al., 2013) to a high of .87 for a pain stimulation task (Taylor et al., 2009). All studies surveyed in Elliott et al. (2020) had sample sizes under 60 subjects, most subjects were adults, and test-retest intervals were all under six months, with most under one month. Moderator analyses did not identify significant differences in reliabilities when comparing task type, event vs block design, scan duration, intertrial interval length, or clinical vs non-clinical sample, but did find lower reliabilities in subcortical relative to cortical brain regions. In a recent review, Noble et al., (2021) identified factors that tend to lead to higher test-retest reliability: shorter test-retest intervals, simple compared to complex tasks, brain regions with stronger activation, cortical regions rather than subcortical, and non-clinical populations. Recent studies in our lab examining the factors affecting test-retest reliability in brain recruitment by risk-taking and response inhibition tasks found that reliability increased with shorter interscan intervals, increasing scan duration, in ROI relative to general brain search space, and with lower subject movement, though the use of the denoising via multirun spatial ICA (Glasser et al, 2018) plus FIX (Salimi-Khorshidi et al., 2014) ameliorated the negative impact of increased subject movement (Korucuoglu et al., 2021).

A major implication of poor reliability for research relying on individual differences is diminished measured effect sizes and statistical power for detecting associations with other variables, or diminished ability to detect changes over time in longitudinal or treatment studies (Elliott et al., 2020). Detecting small effects requires large samples, which is especially problematic for MRI research, given the high cost of assessments (Dick et al., 2021).

Most previous studies of test-retest stability of task-fMRI were conducted in adult samples, and evidence for temporal stability of individual differences in task-fMRI in children is scarce (Herting et al., 2018), despite the widespread use of task-fMRI in developmental research in pediatric samples. Stability of individual differences is particularly important for longitudinal studies that aim to establish prospective associations between developmental changes in task-related brain activations and behavior. As one of the primary goals of much of developmental psychiatric imaging research is to track how neurofunctional development is associated with future onset and course of mental disorders and substance use (Bjork et al., 2018; Feldstein Ewing et al., 2018; Giedd et al., 2008; Volkow et al., 2018), knowing what neurofunctional variables show stable individual differences is critical. Systematic age-related changes due to development do not necessarily preclude test-retest stability of individual differences, provided it is operationalized as rank-order stability, such as with the “consistency” ICC(3,1) (Shrout and Fleiss, 1979). However, individual variation in the *rate* of developmental changes will result in decreases in longitudinal test-retest stability because it would alter rank-ordering between individuals.

The Adolescent Brain Cognitive Development^SM^ (ABCD) Study is an ongoing longitudinal project examining the neuropsychological development of ∼12,000 individuals nine to ten-year-olds at enrollment from 21 sites across the United States of America through adolescence (Casey et al., 2018). The ABCD Study^(C)^ protocol included three fMRI tasks focused on neurocognitive constructs deemed essential for the understanding of adolescent development: response inhibition (Stop Signal Task; SST), reward anticipation and processing (Monetary Incentive Delay; MID), and working memory (nBack; Casey et al., 2018). However, reliability of brain activations elicited by these tasks in the ABCD data has not been established. The recent 3.0 release of ABCD data contains fMRI data for two longitudinal fMRI assessments conducted two years apart (baseline and the first follow-up), and each of these sessions has two approximately five-minute runs for each task. This enables test-retest reliability assessment at two time scales.

Our goal was to examine both within-session, between-run reliability (which is analogous to split-half internal consistency reliability in psychometrics; Heale and Twycross, 2015) and between-session longitudinal stability of regional brain activations elicited in the three ABCD fMRI tasks. This information is essential to evaluate potential utility of the task fMRI data for predictive and inter-individual association analyses, as well as to evaluate potential effects of different region- and subject-level factors on reliability such as relevance of the brain region to the targeted neurocognitive construct, the magnitude of activation, amount of in-scanner movement, and sex. Additionally, an important factor that may affect within-session reliability is change in task-related BOLD responses over the course of the session (possibly reflecting task habituation, sensitization, or restructuring of activity as the task proceeds due to factors like task learning, automation, and attention). These factors can be measured by computing within-session change in activation, and can reduce reliability if the degree of change is inconsistent across individuals. For example, a previous study reported significant habituation of amygdala activation to emotional faces over a 4.5 min task; furthermore, the rate of decrease had a reliability of ∼.5 (Plichta et al., 2014), far exceeding the reliability of the activity itself (.16 for the left amygdala, -.02 for the right; Plichta et al., 2012).

Our approach is not wholly analogous to most fMRI reliability studies (surveyed in Elliott et al., 2020). While we are using the same intraclass correlation analysis approach as other researchers, the intervals between the scans being compared differ (no interval or two years vs. one day to six months). Moreover, the two-year span of ABCD is occurring across a major period of brain development across adolescence. Our between-session analyses may thus be subject to developmental effects that could make a task appear less reliable than reliability analyses using a short test-retest interval or a similarly-long interval between points in adulthood that would presumably be less impacted by developmental differences. For this reason we are hesitant to label the between-session intraclass correlation results as test-retest reliability and instead prefer the term longitudinal stability (note that the consistency intraclass correlation measure used here allows for group level differences; reliability does not decrease if everyone changes in the same direction and to the same extent).

We hypothesized that both within-session reliability and longitudinal stability would be poor on average, given previous research for the MID, SST, and nBack (Blokland et al., 2017; Caceres et al., 2009; Fleissbach et al., 2010; Holiga et al., 2018; Korucuoglu et al., 2021; Plichta et al., 2012; Schlagenhauf et al., 2008; Zanto et al., 2014), with within-session ICCs potentially negatively impacted by variable within-session change across individuals. Of note, fractionating the events of a event-related task that requires multiple runs in order to attain enough events to model, such as the MID task, may result in tepid and unreliable activation in each run *singly*, if the number of events of a given trial type (e.g. reward/loss magnitude) dips below a critical non-linear inflection point of events needed within a single run to yield a reliable activation given a certain n of subjects (Chen et al., 2021).

We also predicted that stability ICCs would be potentially negatively impacted by the long retest interval (Noble et al., 2021) and developmental change between sessions. Nonetheless, it is important to empirically establish the reliability and stability of the ABCD task fMRI data, and if these are indeed generally poor, assess how poor ICCs will affect results. It is also important to investigate some of the factors that may influence ICCs, as a way to understand potential avenues for maximizing ICCs. In that regard, we expected ROIs to have modestly higher ICCs than regions with lesser task relevance and by extension less consistent incidence of activation in the literature. We expected within-session reliability to increase with age, as movement decreases with age in developmental samples (Engelhardt et al., 2017) and movement is a considerable source of additional variance in imaging research (Bright and Murphy, 2017; Diedrichsen and Shadmehr, 2005). Consistent with our previous findings in an adult sample (Korucuoglu et al., 2020; Korucuoglu et al., 2021), we expected more active regions to be modestly more reliable. As developmental change typically occurs at different times and rates (Marceau et al., 2011) and the pubertal hormones associated with development are also related to functional activity in reward, emotional processing, and cognition processes targeted by the imaging tasks (Dai and Scherf, 2019), we expected regions that exhibit greater mean longitudinal change to also have lower longitudinal stability (i.e., a negative correlation of between-session change with between-session ICCs) as it seems likely (although not certain) that regions with greater mean longitudinal change will concurrently be more likely to have changes in rank-order between individuals over that interval given the variance in onset and speed of change, and thus lower ICCs. We also expect that activation estimated from larger regions is less susceptible to noise (though this assumes the region is filled with consistent activity), and therefore expect larger regions to be more reliable relative to smaller regions.

## Methods

### Participants

The individuals and data used for our study come from the ABCD Study’s “Curated Annual Release 3.0” (https://nda.nih.gov; NDA Study 901; DOI 10.15154/1519007). This data release includes two sessions worth of imaging data (structural, task fMRI, and resting state fMRI), with 10,432 individuals in the baseline session (having structural scans that passed ABCD’s pre- and postprocessing quality control and fMRI scanning fields that covered the participant’s brain), and approximately half of them (n=5,205) having processed data available from their first follow-up visit (on average two years later). Task fMRI data was required to pass ABCD’s quality control recommended inclusion flag^2^, leaving 7730 to 8737 individuals, depending on task, within the baseline session [mean (SD) age = 9.94 (0.63), 51% male across tasks] and 4264 to 4438 individuals at first follow-up [11.93 (0.64) years old, 53% male] (Table 1). Participants were recruited primarily through school systems with the aim of reflecting American diversity in sex, urbanicity, race and ethnicity, and socioeconomic status (Garavan et al., 2018). Informed assent was gathered for ABCD participants and consent from their parents or guardians. All procedures were approved by the central ABCD Institutional Review Board (IRB) and/or the IRB for the local scanning site.

**Table 1:** Participants with usable data by task and session and number of participants passing QC criteria. Number of usable participants for each task at each session with mean and standard deviation of their ages, the number that are male, and the number passing different quality control criteria. Sample (Male): Sample size for each task at each session; () number male; Age (SD): Mean (Standard Deviation) age for each session and task, Performance: Participants with sufficient behavioral performance, fMRI: Participants with sufficient frames of fMRI data, E-Prime: Participants with matching E-prime and imaging timing, Series: Participants with a usable series of data.

|  | Baseline |  |  | Follow-Up |  |  |
| --- | --- | --- | --- | --- | --- | --- |
|  | MID | nBack | SST | MID | nBack | SST |
| Sample (Male) | 8737<br>(4397) | 7730<br>(3956) | 8034<br>(4071) | 4426<br>(2353) | 4438<br>(2333) | 4264<br>(2228) |
| Age (SD) | 9.93<br>(0.63) | 9.96<br>(0.63) | 9.94<br>(0.63) | 11.93<br>(0.64) | 11.93<br>(0.63) | 11.93<br>(0.63) |
| Performance | 8871 | 7841 | 8259 | 4870 | 4497 | 4358 |
| fMRI | 8996 | 8768 | 8861 | 4556 | 4517 | 4492 |
| E-Prime | 9496 | 9278 | 9403 | 4785 | 4725 | 4749 |
| Series | 9533 | 9335 | 9241 | 4895 | 4846 | 4867 |

### ABCD Study: Data, Processing and Task Description

Each of the three fMRI tasks collected by ABCD consist of two approximately five-minute consecutive runs. The released task-activation data were processed through ABCD’s “Data Analysis, Informatics and Resource Center” (DAIRC) image processing pipeline (Hagler et al., 2019), which includes motion correction and frame censoring by degree of movement, correction for susceptibility-induced distortions, functional-structural coregistration, activity normalization, and activity sampling onto the cortical surface, carried out using FreeSurfer (Fischl et al., 2002), FSL (Jenkinson et al., 2012), and AFNI (Cox, 1996). Imaging data quality and task performance were evaluated by ABCD’s DAIRC as part of quality control. Based on their evaluation, at baseline, 26% of MID, 34% of nBack, and 32% of SST scans failed quality control; at follow-up those percentages were 22%, 21%, and 25%, respectively. A breakdown of the number of subjects who passed the ABCD’s quality control measures is available in Table 1. Poor behavioral performance and insufficient fMRI frames appear to be the main causes of participant exclusion for both sessions. The ABCD Release 3.0 data provides estimated activation betas for each run and modeled contrast included in the task general linear model, for cortical parcels in the gyrus-defined Desikan-Killiany parcellation (68 parcels, Desikan et al., 2006) and a more granular gyrus- and sulcus-defined Destrieux parcellation (148 parcels, Destrieux et al., 2010), as well as for thirty subcortical structures based on the FreeSurfer segmentations (Fischl et al., 2002). These approaches use the individual’s own structural data to derive the boundaries of these different regions, rather than applying a generic common space labelled atlas.

The ABCD fMRI task battery includes the Monetary Incentive Delay (MID), Stop-Signal (SST), and nBack tasks (Casey et al., 2018). The MID task is designed to elicit functional activity when people are anticipating and experiencing different magnitudes of reward and loss. The SST is designed to elicit response inhibition and error monitoring activity by asking participants to respond quickly to a “Go” cue, unless it is followed by a second “Stop” cue that prompts participants to cancel their response. The Emotional nBack is designed to elicit brain activations related to working memory, with a value-added probe of social information processing by showing participants blocks of images of places or emotional or neutral faces. The task requires participants to determine whether the current image matches a static target (0-back condition) or the image that occurred 2 images back (2-back condition). These tasks were implemented by ABCD because brain signatures of reward anticipation, response inhibition, error processing, and working memory change considerably during adolescence (Blakemore et al., 2010; Sheffield Morris et al., 2018) and have important implications for risk of substance use and psychopathology (Bjork et al., 2017; Giedd et al., 2008). For more information about these tasks, see the Supplemental Methods *Expanded Task Description* section or Casey et al. (2018).

### Data Analysis

Within-session reliability was calculate with the ICC(3,2) formula and longitudinal stability with the ICC(3,1) formula (Supplementary Methods *Reliability Formulae* section; McGraw and Wong, 1996; Shrout and Fleiss, 1979) using the ‘icc’ command in R’s IRR package (Gamer et al, 2019). Both are “consistency” measures of reliability, which focus on the relative stability of values. The ICC(3,2) model estimates reliability for the *average* of the two inputs (i.e., the estimated reliability one would achieve using the full 10 minutes of data), which is appropriate since we assume that researchers will almost always use both runs of within-session task data in their analyses. Conversely, we used ICC(3,1) for estimating longitudinal stability, since sessions are again the “basic level” at which analyses will be conducted with ABCD data. Cicchetti (1994) provides guidelines marking reliabilities below .4 as poor, .4-.59 as fair, .6-.74 as good, and .75-1.0 as excellent. Note that negative ICC estimates are possible for the sum-of-squares based estimator in the ‘irr’ package; negative ICCs were kept as is (i.e., not dropped or set to zero) to avoid an upward biasing of results. Analyses examining longitudinal stability, change between-session, and activity for each session used beta values provided in the ABCD Release 3.0 that were already averaged across both runs, weighted by the number of retained frames in each run. All other analyses used run-specific beta values from the ABCD Release 3.0.

Significance of functional activity for each contrast was calculated as a one-sample t-test against a null-hypothesis of 0. Differences in activity between sessions (activity at follow-up minus baseline) and within-session (activity in run 2 minus run 1) was tested using a paired t-test, reported as Cohen’s D effect sizes (mean of the pairwise differences divided by standard deviation of the pairwise differences), and thresholded for significance (p < .05) after a false discovery rate (FDR) correction for multiple comparisons across the number of regions (but not across contrasts or tasks). This correction was done on the combined Destrieux parcels and FreeSurfer subcortical structures. We only report data from the Destrieux parcellation and the FreeSurfer subcortical segmentation in the main text. The Desikan-Killiany reliability, stability, activity, and change values were calculated and are reported in the Supplementary Materials, but were not used for any statistical analyses, as the more granular Destrieux parcellation allows for better localized estimates of reliability, stability, activity, and change. Subcortical structures analyzed were limited to those with gray matter, excluding ventricles and white matter, leaving 19 structures [left and right hemisphere accumbens, amygdala, caudate, cerebellum cortex, hippocampus, pallidum, putamen, thalamus, and ventral diencephalon, plus the brainstem (which contains both gray and white matter)]. 95% confidence intervals for the reliability, stability, activity, and change analyses are provided in the Supplementary Materials *Data Output*. The variance components used to calculate consistency ICCs, namely MSR (the variance across each individual’s average value) and MSE (the error/residual variance) were calculated to better understand what factors were responsible for differences in within-session reliability for each session. The formulae for the variance components and ICCs can be found in the Supplemental *Reliability Formulae* section.

The initial ABCD quality controlled (QC) sample was the basis for the creation of three additional samples that were used to examine the effects of statistical approaches to data cleaning, namely outlier removal (QC+OR), motion regression (using the framewise displacement variable^3^) followed by outlier removal (QC+MV+OR), and rank normalization (QC+Rank). These samples were compared in paired-t tests. For more details, see the *Data Cleaning* section of the Supplement.

Within-session reliability, longitudinal stability, activity, and change statistics for each contrast and sample were converted into CIFTI ‘pscalar’ (parcellated scalar; Glasser et al., 2013) format for display and data dissemination purposes. Data is available on BALSA at https://balsa.wustl.edu/study/7qMqX. Maps of significant activity and change were created for only the QC and outlier removed (QC+OR) samples, as the movement regression (QC+MV+OR) and rank normalization (QC+Rank) approaches both mean center the data, rendering the computation of activity and change in those samples moot. Region and contrast specific reliability, stability, activity, and change values are also provided as supplemental tables. The R code used to generate the datasets, reliability, stability, activity, change, and ICC components are provided as supplements. All subsequent analyses comparing reliability, stability, activity, and change were performed using SPSS v27 (IBM Corp, 2020).

### Reliability, Stability, Activity, and Change

#### Regions of Interest

As our primary analysis we examined if the regions most consistently recruited by the cognitive demands of each specific task in previous research were more reliable, stable, significantly more active, or subject to greater within or between-session change. To this end, of the total 26 contrasts (10 MID, 9 nBack, 7 SST) included in the processing of the ABCD Release 3.0 data, we identified ROIs for eight targeted contrasts by taking the peak and secondary peak^4^ points from meta-analyses that report a priori important regions for each process targeted by the task/contrast, converting to Montreal Neurological Institute (MNI) coordinates if necessary using the converter included with GingerALE version 3.0.2 (Eickhoff et al., 2011), and identifying the Destrieux parcel or subcortical structure in which this coordinate fell. The same approach was used by Korucuoglu et al (2021). This is not an ideal approach as the parcels/structures are originally generated based on an individual’s specific anatomy and some variation in location within MNI space can be expected, but is reasonable given that the meta-analyses themselves report results in a common (MNI or Talairach) space. The targeted contrasts and their associated meta-analyses are as follows: MID (Supplemental Figure 1): anticipation of reward (large and small reward trials admixed) vs neutral, anticipation of loss (large and small loss trials admixed) vs neutral, and reward vs missed-reward feedback, positive vs neutral from Oldham et al. (2018); nBack (Supplemental Figure 2): 2- vs 0-back from Yaple et al. (2018), face vs place, and emotional face vs neutral face contrasts, both in Muller et al. (2018); SST (Supplemental Figure 3): correct stop vs correct go from Swick et al. (2011) and incorrect stop vs correct go from Neta et al., (2015). There were a total of 57 unique regions (35 ignoring laterality) across the 8 contrasts. Between 7 and 20 ROIs were identified for each contrast, with 20 regions appearing in at least 2 contrasts. See Supplementary Table 1 for a list of ROIs by contrast.

### ROI vs Non-ROI Comparison

The resulting ROIs can be considered to represent the regions that meta-analyses have established as among the most “task relevant” for the principal domains (i.e., contrasts) targeted by each task. To examine the impact of this “task relevance”, for each of reliability, stability, activity, and change (both within and between-session), we directly compared the ROI results with the remaining regions (“non-ROIs”, i.e., rest of the brain) for each of the 8 contrasts, using an independent sample t-test. We used an FDR correction across the number of contrasts, but the analyses for reliability, stability, activity, and change were each treated independently.

### Whole Brain

As it is debatable what regions should be considered ROIs, and since, to the best of our knowledge, meta-analyses to guide selection of ROIs were unavailable for 18 of the 26 contrasts, unbiased, whole-brain analyses were also conducted for all contrasts. Whole brain analyses also included four condition vs baseline contrasts while the ROI analyses were restricted solely to condition vs condition contrasts.

### Movement Quartile Comparison Analyses

To examine the effects of in-scanner movement on ICCs, the QC sample was first subdivided into four subgroups based on quartiles of mean framewise displacement, then framewise displacement was regressed from beta values (within quartile), and finally outliers greater than 3 standard deviations from the mean were removed (within quartile recursively, until no new additional outliers were identified). Reliabilities and stabilities were computed separately for each movement quartile. Each quartile’s regional values were compared against each other quartile for each ICC measure using paired t-tests. Comparisons surviving an FDR correction for the number of quartile comparisons (6) for a reliability/stability measure multiplied by the number of contrasts were reported in the form of the average difference between quartiles. This was done for both ROIs from the 8 targeted contrasts, and at the whole brain level across all contrasts. Further details can be found in the Supplemental *Movement Quartile Samples* section.

### Analyses of Age-Related Change in Reliability

Within-session reliabilities of the baseline and follow-up sessions were compared to examine how age affects reliability. Within-session reliability at the contrast-level at baseline and follow-up were compared (follow-up minus baseline) to each other in paired-t tests using whole brain data from the QC+OR sample (the movement quartile analyses used the QC sample as outlier distributions would likely vary across movement quartile but not intersession interval quartile/decile). Contrasts with a significant difference in reliability across regions after an FDR correction for number of contrasts are reported in the form of the average change in reliability across regions from baseline to follow-up. Similar paired t-tests were computed for the variance components that are used to calculate within-session reliability, to examine the contribution of the separate ‘MSR’ and ‘MSE’ components to the age-related difference in the within-session reliabilities.

### Post-Hoc Analyses of Correlates of Reliability and Stability

Reliability and stability were correlated with the absolute value of activity and change, regional volume, and with each other to establish what relationships exist between these measures. A more detailed overview of these methods can be found in the Supplemental Methods’ *ICC’s Association with Activity, Change, Volume, and other ICCs* section. Post-hoc analyses of the following topics were also explored: differences in ICCs between occipital and non-occipital regions; differences in ICCs between condition vs condition contrasts and the relevant condition vs baseline contrasts; sex differences in ICCs; differences in stability computed separately for run 1, run 2, and the averaged data (of both runs); comparison of within- and between-session change; the longitudinal stability of within-session change; the between-session change in behavioral variables that may be associated with change in within-session reliability; differences in ICCs by length of intersession interval; and similarities in individual activity across condition vs baseline contrasts, since high correlation of activity across different condition vs baseline contrasts has been found to be associated with poor reliability in the composite condition vs condition contrasts (Infantolino et al., 2018). The methods, rationale and interpretation of each of those analyses are described in the Supplemental Methods *Data Cleaning* and *Analyses* sections.

## Results

### Reliability, Stability, Activity, and Change in ROIs

Mean reliability and stability in ROIs, averaged across the targeted contrasts for all 3 tasks in the full “QC” sample, was .069 (SD=.075) for within-session reliability at baseline, .088 (.089) for within-session reliability at follow-up, and .054 (.052) for longitudinal stability (Figure 1A). All ROI reliabilities and stabilities were poor (i.e., < .4). In the QC sample, only 0.62% of ROIs had reliabilities or stabilities over .3 while only 5.9% had reliabilities or stabilities over .2. These poor ICCs occurred despite the fact that the ROIs were indeed generally activated at the group level by their respective tasks (Figure 1C) – 100 of 108 ROIs were statistically “active” (after FDR correction) at baseline and 96 were active at follow-up, with an average (absolute value) Cohen’s D of 0.228 (0.157) at baseline and 0.253 (0.167) at follow-up. An appreciable percentage of ROI ICC estimates were negative, with 15.7% negative within-session reliabilities at baseline, 13.9% within-session reliabilities at follow-up, and 12% negative for longitudinal stability. ROIs were also subject to statistically significant change in activation (Figure 1D), but only in 69 ROIs within-session at baseline, 35 ROIs within-session at follow-up, and 27 ROIs between-session.

**Figure 1:**
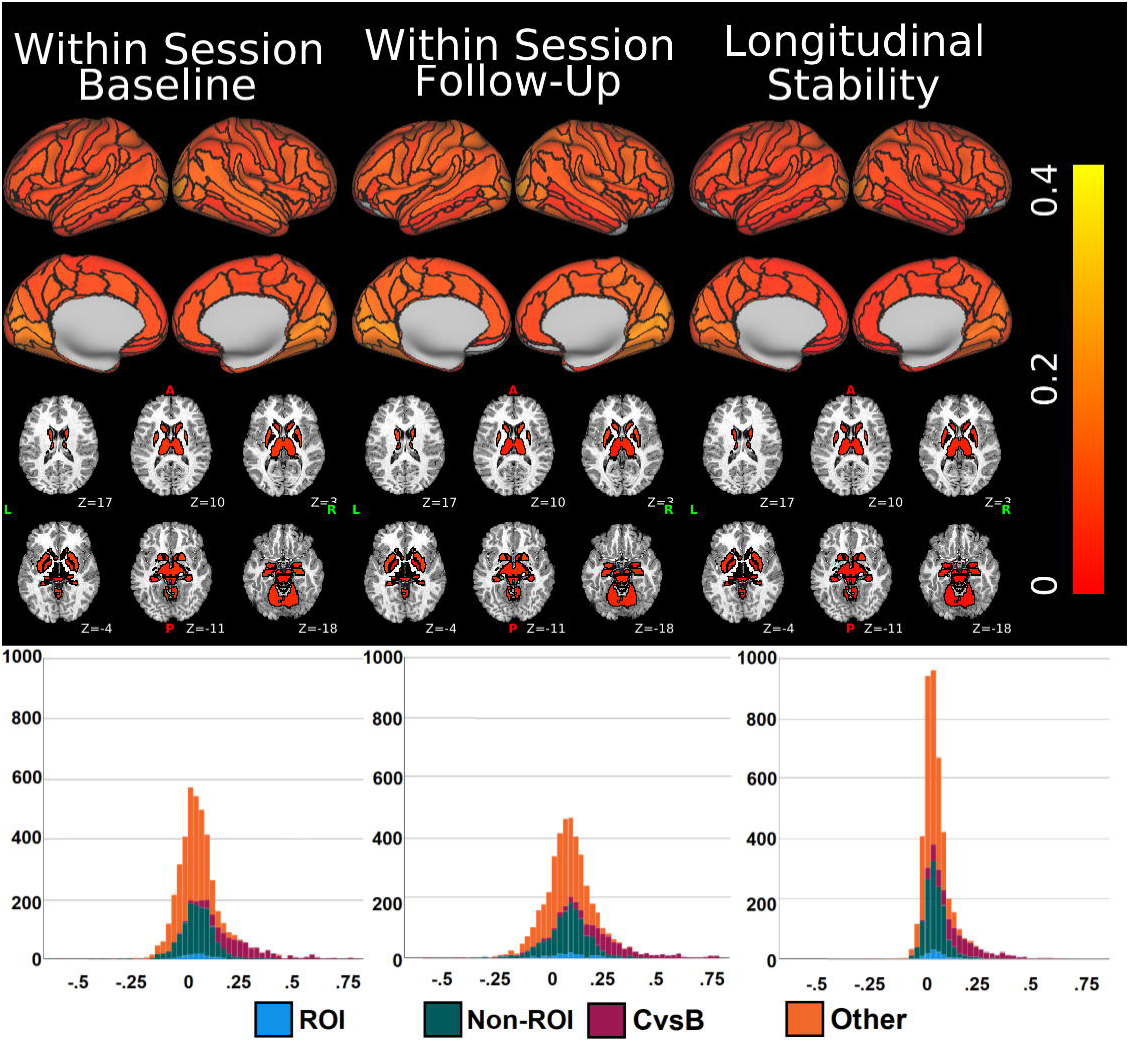
A comparison of average reliabilities, stabilities (A and B), absolute values of activities (C), and absolute values of within- and between-session changes (D) for ROIs (blue) and non-ROIs (green). Data was cleaned using different data-cleaning approaches (A) and also separated into movement quartiles (B) to assess the impact of those factors on reliability and stability. Comparisons between ROIs and non-ROIs that were significantly different in independent sample t-tests (not adjusted for multiple comparisons) are marked with an asterisk. QC: all data that passed ABCD’s quality control; QC+OR: QC sample with outliers removed; QC+MV+OR: QC sample with movement regressed and then outliers removed; QC+Rank: QC sample with rank normalization. Activity and change analyses are available only for QC and QC+OR as the movement regression and rank normalization processes demean the data, making meaningful between region comparisons impossible. ROI: Regions of Interest, N-ROI: Non-ROI.

Data cleaning slightly increased mean ICCs (though average values remained poor) from .070 (.074) for the QC sample (mean (SD) across ICCs) to .092 (.076) for QC+OR, to .091 (.076) for QC+MV+OR, and to .093 (.075) for QC+Rank. While these increases in mean ICCs were small, they occurred consistently, such that the increase was highly significant (all p values from paired t-tests comparing data cleaning types to QC sample < .001, Cohen’s D for paired comparisons of QC vs QC+OR sample: -0.370; vs QC+MV+OR: -0.349; vs QC+Rank = -0.437; Figure 1A).

A much bigger impact was observed by subsetting participants into different movement quartiles, where mean ICCs in ROIs was more than three times higher in the lowest movement group [1st quartile; average (SD) across ICCs of .156 (.120)] compared to the highest movement group (4th quartile; .044 (.070); paired comparison significance p < .001; 1st-4th Cohen’s D = 0.958; Figure 1B). Nonetheless, mean ICCs even of the lowest motion quartile remained well within the ‘poor’ range. Mean and standard deviations for ROIs by contrast and sample can be found in Supplementary Table 2. Analyses comparing reliabilities and stabilities to each other, to the absolute value of activity, and absolute value of within- and between session change (using values derived from the QC+OR data) found significant positive correlations for all analyses (Table 2). Notably, within-session reliability and longitudinal stability were correlated at greater than .80 (across regions, see Supplemental Figure 4B for the 2 vs 0-back correlations of reliability and stability), indicating that more reliable regions tended to be more reliable, regardless of when assessed. More modest correlations (.410-.583) were observed between reliabilities and stabilities with the absolute value of activity and between-session change, while the correlations between ICCs with the absolute value of within-session change were weaker, though still significant (.231-.435). These results show that more reliable and stable regions tend to also be more active, as hypothesized, but also more subject to change, in contrast to what we hypothesized.

**Table 2:** Correlation between reliability, stability, |activity|, and |change| in ROIs. Pearson correlations between ICCs, the absolute value of activity, and the absolute value of change, within and between-session using results from the QC+OR analyses. All correlations significant after an FDR correction for number of comparisons.

| Statistic |  | ICC |  |  | \|Activity\| |  | \|Change\| |  |  |
| --- | --- | --- | --- | --- | --- | --- | --- | --- | --- |
|  |  | Within Baseline | Within Follow-Up | Stability | Baseline | Follow-Up | Within Baseline | Within Follow-Up | Between |
| ICC | Within Baseline | - | 0.879 | 0.817 | 0.590 | - | 0.290 | - | 0.417 |
|  | Within Follow-Up | 0.879 | - | 0.802 | - | 0.583 | - | 0.231 | 0.571 |
|  | Between | 0.817 | 0.802 | - | 0.443 | 0.503 | 0.435 | 0.302 | 0.410 |

The preceding analysis used mean ICCs across *a priori* defined ROIs as a way to broadly summarize our findings. However, the different tasks and contrasts are targeting different aspects of functional processing and it is natural to wonder if ICCs may be higher in particular contrasts. Thus, analyses were repeated at the contrast level. Mean ICCs (across the ROIs for each contrast) was highest in the 2 vs 0-back contrast [mean (SD) ICC for within-session reliability at baseline: .113 (.077); for reliability at follow-up: .160 (.070); longitudinal stability: .103 (.053)] and was lowest in the emotion vs neutral face contrast [ICC for within-session reliability at baseline: .053 (.074); for reliability at follow-up: .009 (.041); and for longitudinal stability: .014 (.033)]. Contrast specific comparisons of the 1st and 4th movement quartiles generally confirmed our finding of higher ICCs in the lowest movement quartile for the individual contrasts, with a significant difference in 19 of 24 comparisons (Table 3).

**Table 3:** ICCs in the 1st and 4th movement quartile for ROIs. Mean for the 1st and 4th movement quartiles for ROIs and their difference, separated by contrast. 1st: mean ICC for the lowest movement quartile; 4th: mean ICC for the highest movement quartile. All bolded Difference results were significant after an FDR correction for the number of contrasts done separately on each ICC type.

| Task | Contrast | Within Baseline ICCs |  |  | Within Follow-Up ICCs |  |  | Stability ICCs |  |  |
| --- | --- | --- | --- | --- | --- | --- | --- | --- | --- | --- |
|  |  | 1st | 4th | Difference | 1st | 4th | Difference | 1st | 4th | Difference |
| MID | Anticipation loss vs neutral | 0.096 | 0.037 | **0.059** | 0.110 | 0.081 | 0.029 | 0.068 | 0.060 | 0.008 |
| MID | Anticipation reward vs neutral | 0.176 | 0.040 | **0.136** | 0.168 | 0.085 | **0.083** | 0.119 | 0.036 | **0.082** |
| MID | Reward positive vs negative feedback | 0.171 | 0.099 | **0.072** | 0.156 | 0.090 | **0.066** | 0.076 | -0.016 | **0.092** |
| nBack | 2 back vs 0 back | 0.343 | 0.086 | **0.257** | 0.396 | 0.164 | **0.232** | 0.240 | 0.098 | **0.142** |
| nBack | Emotion vs neutral face | -0.020 | 0.024 | -0.044 | 0.027 | 0.004 | 0.023 | 0.005 | -0.010 | 0.015 |
| nBack | Face vs place | 0.180 | 0.076 | **0.104** | 0.210 | 0.045 | **0.165** | 0.169 | 0.014 | **0.155** |
| SST | Correct stop vs correct go | 0.224 | -0.036 | **0.260** | 0.163 | 0.039 | **0.124** | 0.142 | 0.015 | **0.127** |
| SST | Incorrect stop vs correct go | 0.265 | 0.051 | **0.214** | 0.228 | 0.106 | **0.122** | 0.186 | 0.059 | **0.127** |

### Comparison of ROIs and Non-ROIs

Reliability and stability values in ROIs were not significantly higher than reliability and stability in non-ROIs, regardless of data cleaning method (Figure 1A), even though *post hoc* comparisons found ROIs were significantly more active (independent sample t-tests comparing one sample Cohen’s D values, mu=0, of ROIs vs non-ROIs using the QC+OR sample; baseline: Cohen’s D 0.576, p = .001; follow-up: Cohen’s D 0.628, p < .001, Figure 1C). ROIs were subject to greater within-session change at baseline before and after data cleaning and between-session, though the between-session differences were only observed after removing outliers (Figure 1D).

Contrast specific analyses found significantly greater ICCs in ROIs relative to non-ROIs only for the 2 vs 0-back contrast of the nBack (Supplemental Figure 5 shows contrast specific bar graphs). Greater absolute value of activity in ROIs relative to non-ROIs was observed in most contrasts at both sessions (10 of 16 analyses, Supplemental Figure 6). Greater absolute value of change in ROIs relative to non-ROIs was found only between-sessions in the 2 vs 0-back and incorrect stop vs correct go contrasts (Supplemental Figure 7). Table 4 details the mean differences between ROIs and non-ROIs when significant differences were observed.

**Table 4:** Comparison of reliability, stability, activity, and change between ROIs and non-ROIs. Mean differences between reliability, stability, the absolute value of activity, and the absolute value of within and between session change between ROIs and non-ROIs. All results significant after an FDR correction for multiple comparisons for number of contrasts with identified regions of interest. W/ - Within session, BL – Baseline session, FU – Follow-Up session, ICC - Intraclass Correlation Coefficient, % - Percent of ROIs with significant activity or change, |Activity| - One sample (mu=0) analyses, |Change| - Paired analysis.

| Task | Contrast | ICC |  |  | \|Activity\| |  | \|Change\| |  |  | % Active |  | % Change |  |  | N ROI |
| --- | --- | --- | --- | --- | --- | --- | --- | --- | --- | --- | --- | --- | --- | --- | --- |
|  |  | W/<br>BL | W/<br>FU | Stability | BL | FU | W/<br>BL | W/<br>FU | Between | BL | FU | W/<br>BL | W/<br>FU | Between |  |
| MID | Anticipation loss vs neutral | - | - | - | - | - | - | - | - | 92% | 92% | 67% | 83% | 0% | 12 |
| MID | Anticipation reward vs neutral | - | - | - | 0.06 | - | - | - | - | 100% | 83% | 78% | 50% | 11% | 18 |
| MID | Reward positive vs negative feedback | - | - | - | 0.19 | 0.14 | - | - | - | 100% | 100% | 50% | 63% | 75% | 8 |
| nBack | 2 back vs 0 back | 0.07 | 0.10 | 0.06 | - | 0.23 | - | - | 0.09 | 100% | 100% | 57% | 57% | 86% | 7 |
| nBack | Emotion vs neutral face | - | - | - | 0.04 | 0.04 | - | - | - | 85% | 85% | 69% | 46% | 0% | 13 |
| nBack | Face vs place | - | - | - | - | - | - | - | - | 85% | 85% | 85% | 55% | 0% | 20 |
| SST | Correct stop vs correct go | - | - | - | 0.21 | 0.22 | - | - | - | 100% | 100% | 65% | 30% | 45% | 20 |
| SST | Incorrect stop vs correct go | - | - | - | 0.20 | 0.19 | - | - | 0.03 | 90% | 100% | 90% | 60% | 70% | 10 |

### Whole Brain Analyses

Since meta-analyses to guide ROI selection were not available for most (18 of 26) of the provided ABCD task contrasts, and since what qualifies as a “region of interest” is partly subjective, whole brain analyses were also performed for all contrasts. Using the QC sample, across all regions and contrasts mean (SD) reliability and stability was .087 (.120) for within-session reliability at baseline, .091 (.130) for within-session reliability at follow-up, and .061 (.080) for longitudinal stability. Contrast and sample specific mean (SD) ICCs can be found in Supplemental Table 3. Figure 2 (top) shows the mean reliability and stability for each region (across all available 26 contrasts). Occipital ICCs tend to be higher than other brain regions, while frontal and temporal polar regions lower to the point that average values were often negative. Figure 2 (bottom) shows histograms of ICCs in ROIs, non-ROIs, and ICCs in condition vs baseline contrasts. The histograms show that ROIs have similar distributions to non-ROIs and that the high end of the distribution is primarily regions from contrast vs baseline contrasts. Using all participants that passed QC, the percentage of ICC values that were negative were 18.4% for within-session reliability at baseline, 19.4% for within-session reliability at follow-up, and 12.3% for longitudinal stability. This far exceeds the percentage of ICCs that were in the fair or higher (> .4) range (2.4% for within-session reliability at baseline, 2.5% for within-session reliability at follow-up, and 0.9% for longitudinal stability). Complete data (and figures) for reliability and stability per parcel by data cleaning and ICC type (within-session at baseline and follow-up and longitudinal stability) can be found on BALSA at https://balsa.wustl.edu/study/7qMqX.

**Figure 2:**
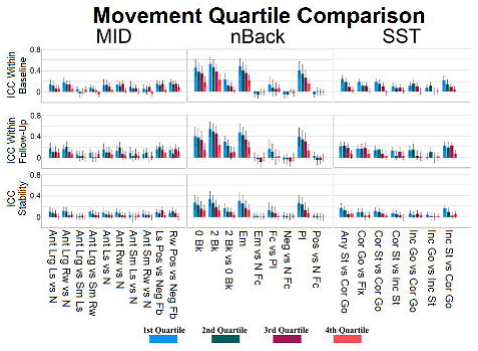
Top: Task fMRI reliability and stability by region, averaged across all 26 contrasts released by the ABCD. Mean ICC values at or below 0 in gray. Bottom: Histograms of reliabilities across all contrasts and regions. Blue: ICCs from ROIs for the 8 condition vs condition contrasts for which meta-analyses to guide ROI identification were available; green: ICCs from non-ROIs; red: ICCs from condition vs baseline contrasts; orange: ICCs from the remaining (18) condition vs condition contrasts (for which meta-analyses to guide ROI identification were not available). ROI: Regions of Interest. N-ROI: Non-ROI. CvsB: Condition vs baseline. All data based on the QC sample.

The data cleaning comparison applied to the whole brain analysis found that removing outliers again slightly increased mean ICCs – from a mean (SD) of .080 (.113) for the QC sample to .102 (.117) for QC+OR (p < .001); see Supplemental section *Effect of Data Cleaning* and Supplemental Table 4 for a comparison of ICCs by data cleaning approaches. Scatterplots comparing region specific ICCs in the 2 vs 0-back contrast before and after outlier removal showed that ICCs were greater in the QC+OR sample relative to the QC sample in most regions (130, 146, and 161 of 167 total regions for within-session reliability at baseline, within-session reliability at follow-up, and longitudinal stability, respectively; Supplemental Figure 4).

*Post hoc* analyses identified higher reliabilities and stabilities in certain regions and contrasts. The following comparisons are based on the QC+OR sample since outlier removal is a common step in data-analysis. ICCs in occipital regions were higher (ICC=.155 (.166)) than in the rest of the brain (ICC=.091 (.100)) and ICCs in condition vs baseline contrasts (ICC=.293 (.149)) were higher than condition vs condition contrasts (ICC=.068 (.066)). For more information, see Supplemental Table 5 for contrast specific occipital results, Supplemental Table 6 for a comparison of condition vs baseline contrasts with the condition vs condition contrasts they are associated with, and sections *Occipital vs Non-Occipital Comparisons* and *Reliability and Stability in ‘Condition vs Baseline’ vs ‘Condition vs Condition’ Contrasts*.

### Movement Quartile Comparison Analyses

A comparison of reliabilities and stabilities computed separately in each of the movement quartiles showed significantly higher ICCs in the quartiles with less movement. Figure 1B shows the average ICCs for ROIs by quartile. For the whole brain, the average reliability increased from .048 for the 4th (highest) movement quartile to .172 for the 1st (lowest) movement quartile within the baseline session (□ = .124), from .068 to .172 within the follow-up session (□ = .104), and longitudinal stability increased from .041 to .122 (□ = .081). Increasing ICCs with less movement was observed in 67 of 78 analyses when comparing 1st to 4th quartiles in paired t-tests, though a minority (6) were significantly less reliable or stable with less movement. Figure 3 shows similar results for the mean ICC across the whole brain, but separated into each of the 26 contrasts provided by ABCD. Those results show that the nBack task had the contrasts with the highest ICCs. Maps of the ICCs for the 2 vs 0-back contrast are shown for all movement quartiles in Figure 4. This figure demonstrates decreasing ICCs with increasing movement and greater reliability within-session at follow-up (when participants were older) relative to the baseline session. Table 3 provides the mean values and their differences for the 1st and 4th quartile by contrast for ROIs. A whole brain comparison of ICCs and their components across quartiles for each of the 26 contrasts is provided in Supplemental Tables 7 and 8. Complete data for regional reliability and stability by movement quartile are available on BALSA at https://balsa.wustl.edu/study/7qMqX.

**Figure 3:**
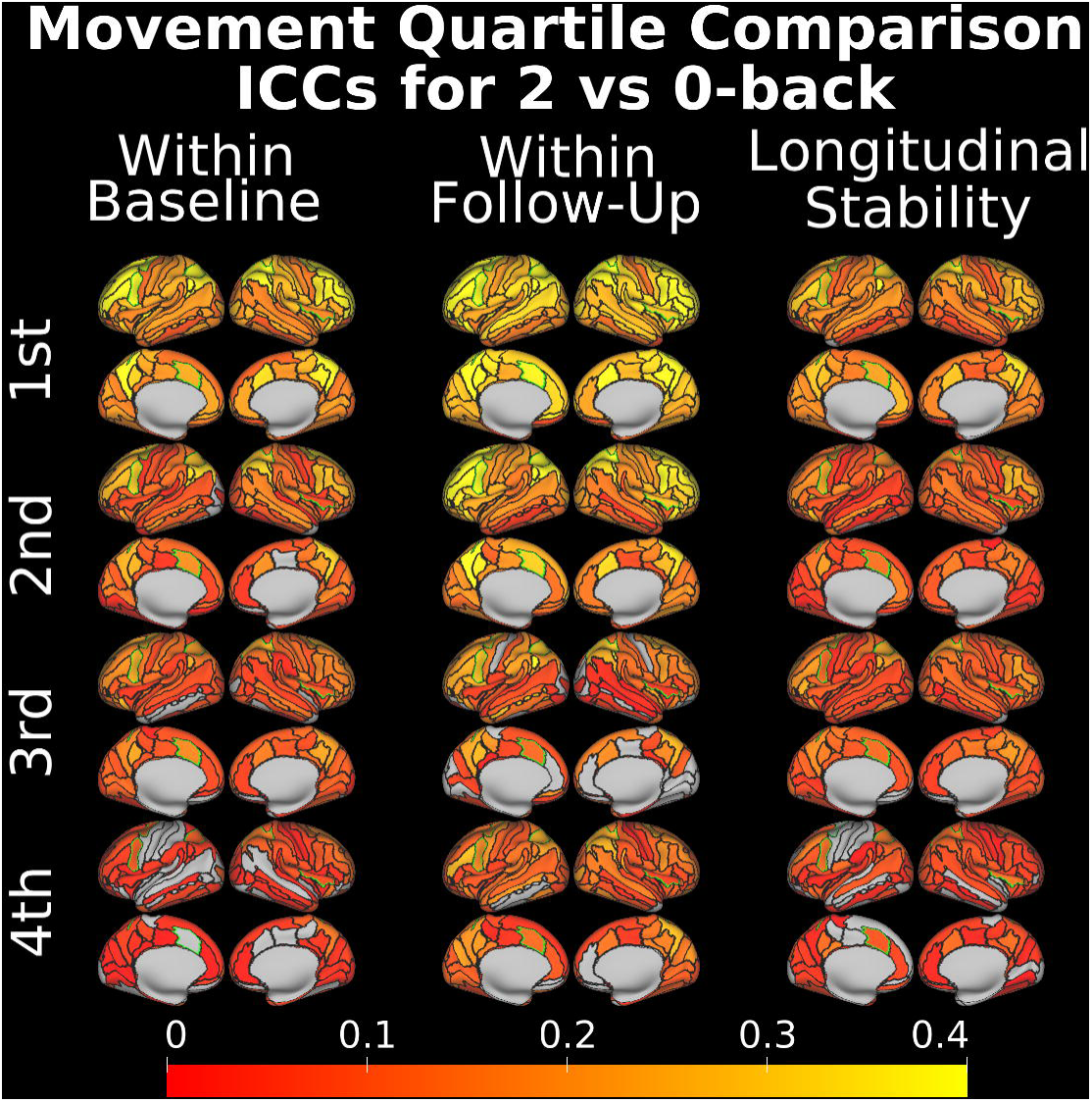
Task, contrast, and reliability/stability measure specific reliability/stability for each movement quartile, averaged across all regions from the whole brain analysis. MID: Monetary incentive delay task, nBack: Emotional nBack task, SST: Stop signal task. W: Within-session. Bars: 1 standard deviation. Ant: Anticipation, Bk: Back, Cor: Correct, Em: Emotion, Fb: Feedback, Fc: Face, Fix: Fixation, Inc: Incorrect, Lrg: Large, Ls: Loss, N: Neutral, Neg: Negative, Pl: Place, Pos: Positive, Rw: Reward, Sm: Small, St: Stop.

**Figure 4:**
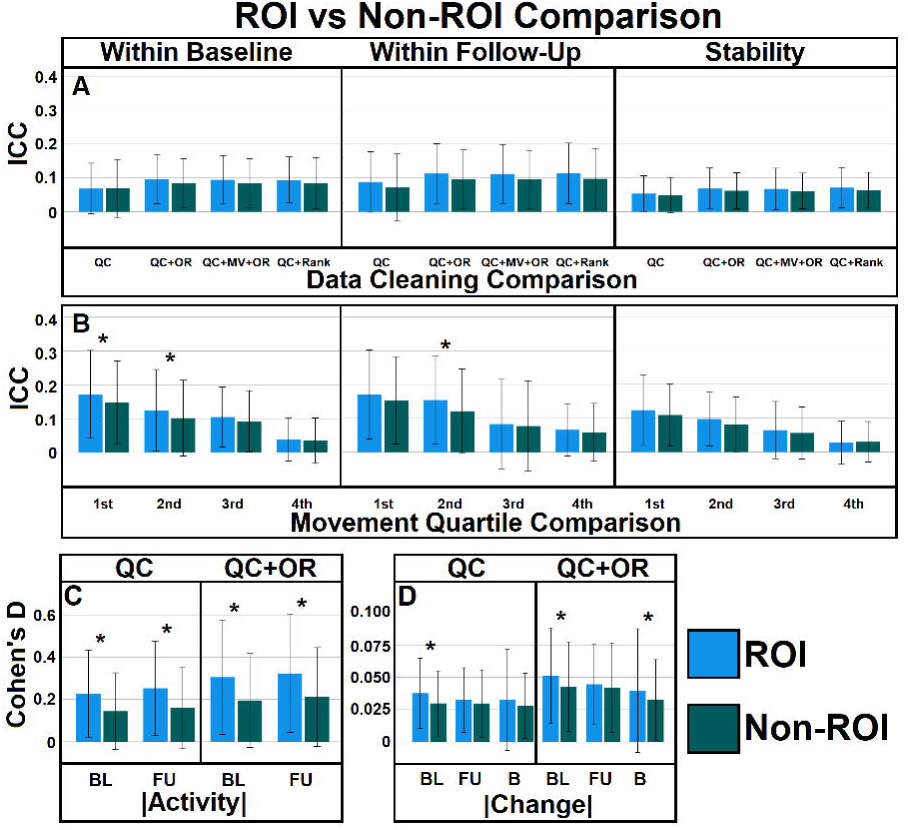
Destrieux parcellation cortical ICCs for each movement quartile (1st = lowest, 4th = highest movement) for the nBack 2 vs 0-back contrast. ICC values at or below 0 in gray. ROIs outlined in green.

### Analyses of Age-Related Change in Within-Session Reliability

Paired comparisons of the within-session reliability measures and their variance components, based on whole brain values from the QC+OR sample, demonstrated significant differences between baseline and 2-year follow-up sessions, possibly due to the age difference (2 years) between them. Mean within-session reliability across all regions significantly increased from baseline to follow-up in most contrasts (15 of 26; average difference across those 15 contrasts of 0.017, with Cohen’s D effect size across them of 0.483). This effect remained in *post hoc* analyses regressing for movement (reliabilities within-session at follow-up > within-session at baseline, paired t-statistic = 19.23, p < .001, Cohen’s D = 0.292). The reliability variance components related to inter-individual variance (MSR) and error (MSE) both significantly decreased between sessions in most contrasts (MSR: 21 of 26 contrasts; MSE: 23 of 26), with the degree of this decrease being greater in MSE than MSR in the majority of contrasts (18 of 26 contrasts). See Supplementary Table 9 for contrast specific comparison of within-session reliabilities by session and Supplementary Table 10 for a comparison of between session change in the variance components used to calculate reliability. A scatterplot comparing within-session reliability at baseline and follow-up for the 2 vs 0-back contrast based on the QC+OR sample can be found in Supplemental Figure 4D.

### High Reliability and Stability Regions and Contrasts

The results observed so far, examining ROI and whole brain ICCs, suggest that the ABCD task fMRI data is largely unreliable and unstable within-subject. However, certain contrasts and regions of course had higher ICCs than the averages reported above. Using the 1st movement quartile sample from the follow-up (within) session reliability (the sample/session with the highest average ICC at .172), 7.0% of the total (26 contrasts by 167 regions) had a reliability between .4 and .59 (fair), 2.3% had reliabilities between .6 and .75 (good), and 0.9% had reliabilities between .75 and 1 (excellent; Cicchetti, 1994), with reliability highest in the left transverse posterior collateral sulcus in the 2-back vs baseline contrast at .899. It is worth noting that 83.4% of these regions with fair to excellent reliability occur in condition vs baseline contrasts which generally do not isolate specific aspects of functional processing. Further, 30.5% of these contrasts/regions (with ICC > 0.4) occurred within occipital cortex^5^, despite being only 17.9% of regions. Only 1.8% (8 of 446) contrasts/regions with reliability > .4 are ROIs (2 vs 0-back’s bilateral intraparietal and posterior transverse sulcus, left inferior precentral sulcus, and left superior precentral sulcus; and face vs place’s bilateral lateral fusiform gyrus, right inferior occipital gyrus and sulcus, and left middle occipital gyrus, all values in the fair range). Within-session reliabilities and stabilities for primary samples/quartiles can be found (and conveniently sorted/filtered) in Supplementary - ICC Output, while full results can be found in Supplementary - Full Output.

### Other Factors Affecting Reliability and Stability

A number of additional analyses were conducted to better understand possible associations between ICCs and other factors. These analyses used whole brain data from the QC+OR samples unless otherwise noted in the Supplementary Methods. There was no significant relationship between reliability or stability and volume of the ROI or non-ROI parcel. See the Supplemental Results’ *Association Between ICCs and Task-Related Activation, Age-Related Change, Region Volume, and other Types of ICCs* section for more detailed results examining the association of ICCs with other statistics. Supplemental Table 11 contains contrast specific correlations of ICCs with other statistics; Supplemental Table 9 contains contrast specific results detailing the relationship between different reliability and stability measures; and Supplemental Table 10 contains contrast specific comparisons of the differences in variance components used to calculate within-session reliability, compared between baseline and follow-up sessions. Sex differences in ICCs were observed, but whether males or females were more reliabile or stable was not consistent across contrasts (See Supplemental Table 12 for a whole brain, contrast specific comparison of sex differences and the *Development and Sex* section of the Supplemental Results).

Variation in habituation may have also degraded reliability in that longitudinal stability was higher in the first run relative to the second of tasks. Moreover, in some MID anticipation contrasts and emotional nBack contrasts, stability based on the first run of data was higher than stability based on the combined runs (Supplemental Table 13 and *Comparison of Longitudinal Stabilities by Runs* section of the Supplemental). Within-session change was generally greater than between-session change (Supplemental Table 14, Supplemental Figure 8 shows significant within- and between-session change). The amount of within- and between-session change was itself not stable, with average longitudinal stabilities across all parcels and contrasts of .013 for within-session change (Supplemental Table 15) and .039 for between-session change (Supplemental Table 16, see *Longitudinal Stability of and Association of Change Within and Between Session* section of the Supplemental for more detailed analysis of the stability of within and between session change). Behaviorally, movement increased within-session and decreased between. Performance on all tasks increased between sessions (See *Change in Movement and Task Performance* section of the Supplemental). Age interval analyses found that reliability and stability were significantly different depending on intersession interval grouping, but not always in the same direction (Supplemental Tables 17 and 18 and *Intersession Interval* section of the Supplemental). Overall, analyses suggested that greater interval was associated with higher longitudinal stabilities, and oddly, higher within-session reliability, even in the baseline session when such differences should be irrelevant. Comparisons of activity between related condition vs baseline contrasts from the nBack task found modest correlations between activity that were related to ICCs in the face vs place contrast, but not the 2 vs 0-back. For more information, see the *Condition vs Baseline Contrast Correlations* section of the Supplement.

## Discussion

### 1. Poor Overall Within-Session Reliability and Longitudinal Stability

Our main finding was that within-session reliability and longitudinal stability of individual differences in task-related brain activation was consistently poor for all three ABCD tasks. Data cleaning approaches like outlier removal, movement regression, and rank normalization significantly increased reliability and stability, but by a small, seemingly inconsequential amount (average change of less than .025). While the finding of poor within-session reliability and longitudinal stability in the ABCD task fMRI data did not come as a surprise, given the mounting evidence for generally lackluster reliability of task-fMRI in mostly adult samples (Elliott et al., 2020; Herting et al., 2018; Noble et al, 2021), the present estimates are far below the .397 average reliability of task-fMRI activation estimated in the meta-analysis by Elliott et al. (2020). The question then arises, what factors could contribute to this particularly disappointing outcome? Previous reliability studies largely involved adult participants who will likely move less in the scanner and used shorter retest intervals relative to the between session analyses (within-session analyses would presumably be subject to habituation/automation/task-reorganization effects that would diminish over a few weeks; Spohrs et al., 2018). Although average ICCs in the ABCD task fMRI data is poor overall and thus subject to a “floor effect”, with limited variability of ICC values across tasks, contrasts, and brain regions, we have examined these and other factors as potential determinants of reliability and stability. Some sample/contrast combinations had values in the fair to excellent range, however these typically occurred in the condition vs baseline contrasts where activity is not specific to task relevant processing and in the low movement quartile samples. Moreover, the highest reliabilities and stabilities were also in occipital lobe, raising the possibility that non-specific responses to actionable visual stimuli are what is most reliable and stable. While the large ABCD sample size means there is still sufficient power to identify effects with only a quarter of the sample, movement itself is frequently associated with measures of interest (e.g., fluid intelligence, externalizing behavior, adiposity; Hodgson et al., 2017; Lukoff et al., 2020; Siegel et al., 2017) and limiting analyses to only low movement samples may therefore lead to a biased sample with results that are not representative of the general population.

### 2. Factors affecting reliability and longitudinal stability

#### 2.1 In-scanner movement

To examine the effect of movement on reliability and stability, participants were separated into quartiles based on movement and ICCs were calculated separately for each quartile. The comparison of ICCs between movement quartiles showed that the lowest quartile (the least moving participants) had an average reliability/stability of .155 while the highest movement quartile average was nearly a third of that value at .053. Although both ICC values are in the “poor” range, this significant difference indicates that efforts to mitigate the impact of movement (including frame censoring and motion parameter regression at the preprocessing stage, as well as excluding subjects with high movement) did not fully control for the effect of movement on ICCs. Decreased ICCs due to movement may be the result of either subthreshold movement artifacts distorting activity or a loss of data due to dropping noisy frames. Frame removal diminishes the amount of data available for analysis, which can reduce the precision of activation estimates and negatively affect ICCs, resulting in a trade-off between data quality and quantity. Though this cannot be addressed using the released computed data, reprocessing ABCD data with different movement thresholds or removing an equal number of frames from low moving subjects (to match frame removal rates from high moving subjects) may better establish how movement affects ICCs. As amount of movement (mean framewise displacement) and number of dropped frames are almost fully confounded (r = .97 in the MID task), we cannot say whether the loss of data or subthreshold movement effects in the retained frames are responsible for the poorer ICCs in high movement quartiles. More generally, our finding of a strong effect of our movement quartiles on ICCs calls for approaches to reduce the impact of movement. Data driven noise removal (ICA-FIX, Salimi-Khorshidi et al., 2014) has been found to increase reliability in high moving adult participants, though by only .06-.08 (Korucuoglu et al., 2021). Given substantially stronger impact of movement on reliability in children, this approach may potentially lead to larger ICC gains in children, including ABCD data.

#### 2.2. The length of retest interval and age-related changes

Poor longitudinal stability in the ABCD task fMRI data, consisting of children aged 9-10 at baseline, could conceivably result from a large intertest interval (two years), during which significant age-related changes can be expected. However, the present findings do not support this hypothesis. First, within-session reliability was also poor for all three tasks. Also, within-session reliability significantly predicted longitudinal stability in most contrasts. While within-session reliability could conceivably be poor and not affect longitudinal stability (if each individual’s within-session change was consistent for both sessions, a measure could be stable between-session and unreliable within), this was not the case, as within-session change was found to be unstable (average ICC = 0.013). Further analysis and discussion of the impact of within-session change can be found in the Supplemental *Within-Session Change* section.

Contrary to our expectation, we found a positive, rather than negative, relationship between longitudinal stability and the extent of age-related changes in task-related activation. This finding can be explained by two factors. First, the ICC(3,1) indicates “consistency” reliability, i.e., stability of individual differences in terms of relative rank-order of individuals, rather than unchangeability of absolute values (Shrout and Fleiss, 1979). Consistency reliability is not affected by systematic changes of values between measurements due to factors like developmental changes, unless the extent of these changes varies across individuals, in which case reliability would be diminished. Since previous studies have reported sex differences in the rate of development (Gur and Gur, 2016), we examined stability separately by sex, but did not find consistent effects of sex on stability. Discussion of sex differences can be found in the Supplemental *Development and Sex* section. Second, statistical power to detect significant changes over time in repeated measures designs, such as age-related changes, increases as a function of test-retest reliability (Rochon, 1991; Vonesh, 1986). Measurements with low test-retest reliability are noisy, and random variance can mask systematic between-session changes, if any. Consequently, in our analysis age-related changes could be best detected for measures that show at least some longitudinal stability.

Finally, comparing groups based on low or high intersession interval did not find that shorter intervals were associated with higher stabilities (rather stability was generally higher with longer intervals). Overall then, our analyses do not find support for developmental change as the cause of poor stability in the ABCD sample.

#### 2.3. Task design and specific contrasts

Overall, reliability and stability were substantially higher for the working memory contrasts, although they were still in the poor range. These task differences may be related to the use of an adaptive procedure to equalize performance across subjects in the MID and SST tasks, which could also attenuate individual differences in task-related brain activation.

Within tasks, there were differences in reliability and stability between specific contrasts, which was most evident for nBack task (because reliability for MID and SST was close to zero, there was too little variability to examine differences across contrasts within those tasks). Contrasts between active condition and baseline consistently showed higher reliabilities and stabilities than contrasts between two active conditions (e.g., greater reliability and stability of 2-back vs baseline compared with 2-back vs 0-back). This is consistent with psychometric and neurofunctional evidence (Caruso, 2004; Infantolino et al., 2018) that contrast (difference) scores typically show lower reliability than their constituent measures because error variances of both constituents contribute to the error variance of the difference score and activity is highly correlated for condition vs baseline contrasts. For fMRI measures, this results in a trade-off between reliability or stability and validity of activation metrics. For example, an activation elicited by emotional faces relative to baseline shows higher ICCs than activation of emotional faces relative to neutral faces (which is totally unreliable in the ABCD data). Similarly, Baranger et al. (2021) recently demonstrated using a number-guessing reward task that reward activation contrasted with baseline had greater reliability than reward contrasted directly with loss. However, contrasts with baseline lack specificity because they may include activation elicited by emotional content as well as nonspecific activation that can be attributed to any outcome regardless of semantic content, and even activation that is common to visual stimuli in general, resulting in poor discriminant validity.

#### 2.4. Regions of Interest

We expected ROIs to be more reliable and stable than other brain regions because task fMRI is specifically intended to activate certain regions engaged in particular cognitive processes, and it seems reasonable that individual differences in the degree of activation under task performance would be more consistent over time than regions that are not task-engaged and whose activity may fluctuate more (e.g., due to subject state). Indeed, this premise has been fundamental to the whole task fMRI endeavor. However, contrary to this expectation, our analyses show that, except for the 2 vs 0-back contrast of the nBack, a priori ROIs are not more reliable or stable than the rest of the brain. Furthermore, across tasks, higher ICCs were observed largely in occipital regions that are generally of limited interest in the context of neurocognitive constructs targeted by the tasks used in the ABCD Study. Reliability, and stability were also significantly correlated with the absolute value of activity in most contrasts. This was most prominent in the face vs place contrast of the nBack (correlations between .78-.81), with most of these relationships having a correlation in the range of .4-.5. This is inconsistent with our finding of greater activity in ROIs relative to non-ROIs but not accompanying greater ICCs in ROIs for most contrasts. A possible explanation is that the effect of activity on ICCs was not strong enough to manifest as greater ICCs in ROIs relative to non-ROIs. A different approach to identifying ROIs (e.g., data driven relative to based on published meta-analyses) may have given different results.

### 3. Implications of Low Reliability in the ABCD Task fMRI Data

The main (and certainly unwelcome) conclusion from the present analysis is that poor reliability and stability of child task fMRI activity in the MID, nBack, and SST calls into question their suitability for most analyses focused on individual differences, as well as any analyses that rely on the assumption (explicit or implicit) that brain activation measures represent reliable and stable trait-like variables. Such studies include correlations between brain activations and individual differences in behavior or psychopathology (particularly, prospective longitudinal brain-behavior associations), within-subject analyses of longitudinal changes, genetic associations, effects of individual differences in environmental exposures, and many other research designs. Reliability imposes the upper limit on the measurable correlation between variables (Nunnally et al., 1970; Vul et al., 2009), and traits with low reliability or stability cannot produce high correlations with other traits, even other highly reliable or stable ones. The very large sample size of the ABCD study affords enough statistical power to detect significant correlations even with low-reliability/stability traits. However, these correlations will predictably be very low, though that does not mean they are not predictive insights into biological mechanisms (Dick et al., 2021).

Since ABCD is an ongoing longitudinal study, a question arises whether there is a possibility that poor reliability and stability found in the present analysis is related to the participants’ young age, and thus whether, in subsequent longitudinal waves, ICCs will improve. Some evidence supports this expectation. We found that within-session reliability increased from baseline to follow-up for most contrasts, albeit by a small amount. Part of this increase is likely due to movement-related effects, since movement decreased between sessions. However, the overall increase over two years was small, with the largest increase in average whole-brain contrast wide reliability being .061. Nevertheless, one can reasonably expect at least some improvement of ICCs with age, at least until the propensity to move in the scanner stabilizes (around the mid-teenage years; Satterthwaite et al., 2012). Our recent study of test-retest reliability of the ABCD SST task in a sample of young adults showed fair and even good reliability for some contrasts/ROIs, though using a different preprocessing pipeline and parcellation (Korucuoglu et al., 2021).

Another parsimonious account of the lackluster ICCs found here (as well as an account for the slight improvement with age from baseline to follow-up) is simply the relatively lackluster task engagement of children compared to adults. This has been evidenced not only by decreases in trial-to-trial reaction time variability from childhood to adulthood in signal detection tasks (Tamnes et al., 2012), but also evidenced in developmental pupillometry studies, wherein for example, task-demand-elicited noradrenergic activation (indexed by pupil dilation) waned during memory encoding in children, while remaining active in adults (Johnson et al., 2014), resulting in poorer recall in children. It stands to reason that as ABCD participants mature into more consistent and unflagging task engagement, this will entail deeper and more consistent encoding of task information, that would lend itself to greater reliability and stability.

Researchers using ABCD task-fMRI data are strongly urged to select variables that show at least some trait stability and evaluate the upper boundary of expected correlations or effect sizes for other analyses. For example, attenuation of observed correlation between two variables can be easily estimated if reliabilities or stabilities of both variables are known with the formula r_ObservedA,ObservedB_=r_A,B_*sqrt(ICC_A_*ICC_B_) where r_A,B_ is the “true” correlation between two constructs (Nunnally, 1970); in the ABCD sample, longitudinal stability of variables can be readily computed using data from subsequent assessment waves. However, reporting “reliability adjusted” correlations is generally inadvisable as the measurement errors responsible for low reliability or stability can be correlated between variables and applying the above formula can bias results, erroneously increasing or decreasing “true” correlations (Saccenti et al., 2020). For cognitive neuroscience research outside of ABCD, we suggest that establishing and reporting test-retest reliability and stability of task-fMRI phenotypes is imperative for planning studies and publishing results. In particular, computations of statistical power should account for imperfect reliability of task-fMRI data, because poor reliability leads to the reduction of the measured effect size and, consequently, increases the sample size needed (Baugh, 2002). *Post hoc* power analyses (calculated with the pwr.r.test R function; Champley, 2018) examining the sample sizes needed to find a significant correlation between a variable with a reliability of .8 and a true correlation of .3 with variables with reliabilities of .099, .123, and .072 (the average reliabilities within-session at baseline, at follow-up, and the average longitudinal stability for ROIs from the QC+OR sample), found that samples would need to be 1099, 884, and 1511 participants, respectively. While far below the sample available for the ABCD Study, this greatly exceeds the average sample size for fMRI studies (Poldrack et al., 2017) and is consistent with research finding that large samples are needed to find a consistent correlation between imaging data and other variables (Marek et al., in press).

The preponderance of small effects in imaging research that would necessarily result from poor ICCs is one of the reasons large, consortium-scale studies like the ABCD are needed (Dick et al., 2021). As the statistical approaches to increasing ICCs addressed here had small effects that did not increase reliabilities or stabilities out of the ‘poor’ range, developmental task fMRI researchers may need to plan studies around the limitation of poor task reliability. As restricting analyses to low movement groups tripled ICCs relative to high movement, a greater emphasis on accounting for movement, either through participant training or processing, may be warranted. Neglecting the reliability and stability challenges in task fMRI research may result in further proliferation of small sample, underpowered studies and dissemination of spurious, false positive and non-replicable findings that undermine the credibility of cognitive neuroscience research relying on task-fMRI data.

Our results do not necessarily mean that task fMRI activity is inherently unreliable or unstable. It remains unknown how much the reliability of task fMRI could be increased by acquiring more data per individual, although the resting-state literature suggests that the gains could be substantial (Birn et al., 2013; Gordon et al., 2017) if the challenges regarding learning and adaptation effects in task performance can be managed. Also, the task data released by the ABCD Study reflects only one approach to task fMRI processing and processing approaches can vary substantially, as do subsequent results, even when using the same data (Botvinik-Nezer et al., 2020). Identifying processing approaches that promote reliable and stable individual level data is vital to identifying reproducible individual differences in functional activity. Cohort (e.g., age) effects may also be a profound factor in the current results. For example, we have reported that SST task activity in young adults has fair to good reliability while using the same scanner, task design, and scan acquisition parameters as the ABCD Study, but processed using a different pipeline (though also with a shorter intersession interval of ∼6 months; Korucuoglu et al., 2021). While an increase in reliability with age is expected (due to less motion), we cannot definitively say that this was the source of the higher reliability in that study, since there were processing differences as well, including the use of the Human Connectome Project pipelines (Glasser et al., 2013) and parcellation using a more functionally relevant multi-modal parcellation (Glasser et al., 2016). Notably, ICA-FIX (Glasser et al, 2018; Salimi-Khorshidi et al., 2014) was able to increase reliability in subjects with high movement (Korucuoglu et al., 2021), albeit to a small extent (average increase in ICC of .06). Ultimately, alternate approaches to processing ABCD data and accounting for noise and movement may result in more reliable data.

### Limitations

These analyses are not without limitations. Reliability values will be influenced by differences in within-session change while stability will be influenced by between-session change/development that cannot be fully accounted for given the way the data was processed for public release (i.e., whole run beta values rather than a more granular analysis of possible temporal effects, such as minute by minute or even block by block estimates). Also, the number of sessions available (2) is currently a limitation, as more advanced statistical approaches to modelling and accounting for individual differences in change are not available with only two sessions of data (e.g., mixed effects modelling of nonlinear trajectories, Herting et al., 2018). It may be the case that reliability/stability using these tasks in this age group would improve with a brief enough intersession interval to avoid developmental differences but long enough to avoid task reorganization/habituation effects, but we cannot establish that with the present data. Meta-analyses were only available for 8 contrasts, so ROIs were not identified for the remaining 18 of 26 released contrasts. We investigated reliability in a univariate framework, and it is possible that more multivariate-oriented analyses will have higher reliability (Kragel et al., 2020), although this remains to be established. Last, data was only available for structurally-based parcellations. However, functionally derived parcellations are likely to be more relevant and may be accompanied by increased reliability and stability.

## Conclusions

Overall, reliability and stability of task-fMRI data in the ABCD sample was poor. Movement decreases ICCs, but even selecting only the lowest movement quartile for analysis didn’t raise average reliability or stability out of the poor range. ICCs were only very minimally improved by the investigated data cleaning approaches. Reliability and stability were generally not better in ROIs relative to the rest of the brain. ICCs tended to be best in working memory related and condition vs baseline contrasts. Decreases in movement with age may somewhat increase ICCs in later ABCD assessment waves. Future ABCD scanning and processing may benefit from a more aggressive approach to controlling movement. For the amount of task fMRI data collected in the current study (∼ 10 min per participant), the MID and SST tasks, and to a lesser extent the nBack task as well, may not be practical for other studies examining childhood development unless they can obtain sample sizes in the 1500+ range.

## Supporting information

Full Output

ICC Output

Code for generating data and results

## Acknowledgements

Data used in the preparation of this article were obtained from the Adolescent Brain Cognitive DevelopmentSM (ABCD) Study (https://abcdstudy.org), held in the NIMH Data Archive (NDA). This is a multisite, longitudinal study designed to recruit more than 10,000 children age 9-10 and follow them over 10 years into early adulthood. The ABCD Study® is supported by the National Institutes of Health and additional federal partners under award numbers U01DA041048, U01DA050989, U01DA051016, U01DA041022, U01DA051018, U01DA051037, U01DA050987, U01DA041174, U01DA041106, U01DA041117, U01DA041028, U01DA041134, U01DA050988, U01DA051039, U01DA041156, U01DA041025, U01DA041120, U01DA051038, U01DA041148, U01DA041093, U01DA041089, U24DA041123, U24DA041147. A full list of supporters is available at https://abcdstudy.org/federal-partners.html. A listing of participating sites and a complete listing of the study investigators can be found at https://abcdstudy.org/consortium_members/. ABCD consortium investigators designed and implemented the study and/or provided data but did not necessarily participate in the analysis or writing of this report. This manuscript reflects the views of the authors and may not reflect the opinions or views of the NIH or ABCD consortium investigators. The ABCD data repository grows and changes over time. The ABCD data used in this report came from NIMH Data Archive Study 901. DOIs can be found at DOI 10.15154/1519007.

## Footnotes

1 It is important to distinguish between measures intended to capture a stable trait-like attribute versus measures that may be heavily influenced by state effects (such as attention, caffeine level, hydration, previous night sleep quality, current anxiety level, etc). A measure could in principle have a high test-retest reliability if measured in a consistent and well-controlled subject state, yet empirically appear to have a low reliability because possible state influences are either not controlled, or the relevant state influences affecting the measurement are simply unknown. While it is highly valuable from a scientific perspective to study the effect of state on both within- and between-subject variance (and thus reliability), a measure that is only reliable under limited, state-specific conditions is by-definition not a stable “trait-like” measure.

2 Variables `imgincl_{mid,nback,sst}_include` of the `abcd_ingincl01` instrument. See “ABCD Release 3.0 release notes”, available at https://nda.nih.gov/edit_collection.html?id=2573

3 tfmri_{mid,nback,sst}_{all,run1,run2}_beta_mean.motion using the harmonized “DEAP” variable name, as specified in the “21. abcd_3.0_mapping.csv” file in the ABCD Release 3.0 release notes, which also provides the mapping to the NDA instrument and corresponding NDA variable name in which the mean framewise displacement values can be located.

4 A part of a large cluster with activity higher than its surrounding voxels that is not the highest point in the entire cluster.

5 Occipital regions are defined as Destrieux parcels with ‘_oc_’ or ‘_occipital_’ in their names as well as the calcarine and cuneus.

