## Supplement for "Reliability and Stability Challenges in ABCD Task fMRI Data"

**Supplemental Methods**

*Expanded Task Description*

The task functional MRI component of the ABCD project consists of three tasks; the MID, SST, and nBack; designed to elicit activity in developmentally relevant domains. The aim of the MID imaging task is to examine functional activity when people are anticipating and responding to reward and loss. In the MID, participants are told they will win or not lose a small ($0.20) or large ($5) amount of money or not win or lose anything if they press a button during the period when a target is briefly shown. Participants are given a brief time to anticipate reward or loss and receive feedback of reward or loss. The amount of time the target is shown is automatically adjusted to ensure a 60% accuracy rate. The SST is designed to elicit response inhibition and error monitoring activity. In the SST, participants are asked to press a button to indicate the direction an arrow is pointing unless another ‘stop’ arrow is presented a short time after the first. The amount of time before the stop signal is presented is also automatically adjusted to ensure a 50% accuracy. The Emotional nBack is designed to elicit working memory, face processing, and emotion processing activity. The Emotional nBack shows participants blocks of images of faces (showing positive, negative, or neutral expressions) or places; the working memory component requires participants to indicate if the image they are currently viewing is the same or different from the image shown two previously (2-Back) or from the image shown at the beginning of the block (0-Back), requiring differing levels of working memory. These tasks were chosen as working memory, error monitoring, response inhibition, reward and punishment processing change during adolescence (Blakemore et al., 2010; Sheffield Morris et al., 2018) and can contribute to substance use and psychopathology (Bjork et al., 2018; Giedd et al., 2008).

*Data Cleaning*

An “outlier removed sample” (referred to as ‘QC+OR’ in figures) was generated from the ABCD’s quality controlled (QC, Release 3.0) task fMRI data . This was accomplished by standardizing the data and having parcels with activity more than three standard deviations away from the mean recursively removed. Recursive removal ensured that the change in sample distribution from only one round of outlier removal did not leave new outliers when restandardized. This was done separately for the baseline and follow-up sessions. The missing data pattern from the standardized, outlier removed data, was then applied to the unstandardized beta values. This procedure removed, on average across all regions and contrasts, 4.25% of the data (standard deviation of 1.84%). Occipital brain regions were the least affected by this outlier removal process, with the medial lingual gyrus in the nBack emotion contrast at follow-up the least impacted (losing 0.74% of subjects in the left and right hemisphere’s parcels). Frontopolar regions were most affected, with the left transverse frontopolar region in run 2 of the baseline session of the nBack’s negative vs neutral face contrast losing the most data, with 12.54% of subjects’ data removed. The QC+OR sample was used for analyses of activity and to estimate change within- and between-sessions.

Another sample was created to address the impact of movement on ICCs (referred to as ‘QC+MV+OR’ in figures). Run, task, and session specific movement was regressed from the data using R’s lm command, using the framewise displacement variable, which combines head rotation and translation measures (tfmri_mid/nback/sst_all/run1/run2_beta_meanmotion using the redcap naming), supplied with the task beta values as an independent variable. The same outlier removal approach (described above) was applied to the movement regressed data as well. On average, 4.26% of the sample (standard deviation 1.84%) was removed from the movement regressed data. For the ‘QC+MV+OR’ sample, the regions most and least impacted by this were the same as in the ‘QC+OR’ sample, with occipital regions losing the least subjects and frontopolar losing the most. The right medial lingual gyrus as the follow-up session in the emotion contrast again retained the most subjects, losing only 0.74% of the sample, while the transverse frontopolar region of the 0-Back contrast at baseline in run 2 lost 12.71%.

The QC sample was subject to rank-based inverse normalization to form one additional sample (labelled ‘QC+Rank’ in figures). This was accomplished using the RNomni (McCaw, 2020) package’s RankNorm command. This approach used the entire QC sample and was not subject to neither outlier removal nor movement regression steps, before or after normalization.

*Movement Quartile Samples*

To further investigate the impact of movement on ICCs, the QC sample was split into quartiles based on movement. Once separated, each quartile was subject to the same movement regression and outlier removal process as the movement regressed sample using all QC subjects. The outlier removal process removed (mean percentile removed(standard deviation of percentile removed)) 2.56 (1.4)% (mean (SD)) of the sample in the first quartile, 2.67( 1.41)% of the sample in the second quartile, 2.82 (1.47)% of the sample in the third quartile, and 4.35 (2.11)% of the sample in the fourth quartile. Quartile assignment was task, run, and session specific, so it is possible an individual could be in the first quartile in the initial run and the fourth quartile in the second run of a given task.

*Analyses*

*Reliability Formulae - Intraclass correlation coefficients (ICCs)*

Both ICC(3,1) and ICC(3,2) depend on a comparison of variance components related to individuals (Mean Square for Rows, MSR), raters/sessions (Mean Square for Columns, MSC), and variance unaccounted for by MSR or MSC (Mean Square of Error, MSE). MSR is equal to the variance of individuals (MSR=variance(mean(rows))*nColumns), MSC is the variance of the rater/session (MSC=variance(mean(columns))*(nRows)), and MSE is equal to the variance across all individuals and raters/sessions (Sum of Squares total, SStotal, or variance(data)*((nRows*nColumns)-1)) minus MSR and MSC ((SStotal-MSR*(nRows-1)-MSC*(nColumns-1))/(nRows-1)*(nColumns-1)). The ICC(3,1) formula is (MSR - MSE)/(MSR+(nColumns-1)*MSE) and the ICC(3,2) formula is (MSR - MSE)/(MSR). While some stability formulas cannot produce negative values (e.g. variance of a random effect/total variance), it is possible to produce negative ICCs if MSE exceeds MSR.

*Effects of Data Cleaning Comparison*

Reliability and stability differences between data cleaning approaches were calculated at the contrast level separately for within-session reliability and longitudinal stability measures. Each approach, (a) all usable subjects (QC), (b) outlier removed (QC+OR), (c) movement regressed and outlier removed (QC+MV+OR), and (d) rank normalized (QC+Rank), were subjected to a pairwise comparison with t-tests to determine which one was most reliable. All comparisons significant after an FDR correction for the total number of pairwise comparisons for each ICC measure multiplied by the total number of contrasts are reported in the form of the average difference in ICC. All analyses were performed in SPSS.

*ICC’s Association with Activity, Change, Volume, and other ICCs*

The relationship between ICCs and other variables was calculated using Pearson correlations in SPSS. Analyses were calculated both by combining all contrasts, regardless of whether or not they were used in ROI analyses and separately for each contrast (referred as 'contrast-level analysis'). Results were reported if they were significant after an FDR correction for the number of contrasts. ICCs were compared against each other to examine if regions more reliable within-session were also more reliable between. While it would seem that longitudinal stability would be contingent on within-session reliability, it is not necessarily so as degree of within-session change could vary between individuals but in a consistent manner across sessions, resulting in the averaged or run specific values used for the longitudinal stability analyses being reliable even if the individual runs composing them within-session are not. To examine how estimated activity and change may be affected by ICC, the absolute value of activity at each session and the absolute value of change, within and between-session, were correlated with ICCs^[1]^. To determine the impact of parcel size on ICC, volume was correlated with reliability and stability. Volume was computed by taking the average across individuals of the volume of each parcel in the Destrieux parcellation and subcortical structure from the FreeSurfer segmentation from the structural data of the baseline session from the ABCD 3.0 release.

*Occipital Analyses*

Post-hoc analyses, carried out after noticing relatively high ICCs in occipital regions and greater activity relative to the rest of the brain, examined how occipital regions differed in ICCs, activity, and change. Reliability, stability, the absolute value of activity at each session, and the absolute value of change, within- and between-session, were compared between occipital and non-occipital brain regions (30 occipital regions, including parcels that bridge lobes, e.g. the lateral occipito-temporal gyrus, out of 167 total cortical and subcortical regions) in independent sample t-tests conducted in SPSS. Values that survived an FDR correction for number of contrasts were reported. Each statistical analysis was subjected to an FDR correction separately.

*Reliability and Stability in Condition vs Baseline vs Condition vs Condition Contrasts*

Researchers would likely want to use the regions and contrasts with the highest ICCs. We observed that condition vs baseline contrasts were more likely to have higher ICCs than condition vs condition contrasts. We tested this relationship by comparing ICCs (calculated using the QC+OR sample) for condition vs baseline contrasts against the condition vs condition contrasts they composed (e.g. 0-back vs baseline vs 2 vs 0-back). Differences significant after an FDR correction for multiple comparisons were reported.

*Sex Differences in Reliability and Stability*

In order to examine a potential component of low longitudinal stability, ICCs were computed separately for males and females and compared in paired t-tests. The consistency IICC approach employed here depends on subjects maintaining the same position relative to other subjects between assessments; that this is a developmental sample and the speed and onset of change can vary between groups (Marceau et al., 2011) may limit stability over long periods as an individual’s relative position will be inconsistent with multiple groups changing at different rates and times. One well known and simple to analyze factor associated with the timing of developmental change is sex. By splitting the sample by sex and calculating stability independently, it is possible to mitigate one source of developmental differences that could be reducing stability by increasing between-session variance. If sex specific differences in developmental onset and trajectory are reducing longitudinal stability, then sex specific stability should increase over stability of the combined sample. If not, the average of male and female stabilities should be similar to the stability of the combined sample. This was examined at the contrast level using paired t-tests, analyses significant after multiple comparison correction for the number of contrasts are reported. Analyses were performed using the outlier removed (QC+OR) sample.

*Effect of Number of Runs on Longitudinal Stability*

Post-hoc analyses examining the effect of the amount of data on stability were performed. Differences in stability based on the amount of data being analyzed first and second half of the task may be indirect evidence of habituation or other factors that could change functional activity, as a decrease in stability with more data may indicate that the effect of interest has changed and may indicate if adding more runs would increase stability. Longitudinal was calculated for run 1 and run 2 separately using the outlier removed sample and with the use of the ICC(3,1) formula. Run specific stabilities were compared against each other and against the stability using all runs in paired t-tests. Results significant after an FDR correction for the number of contrasts are reported.

*Comparison of Change Within- and Between Session and Stability of Change*

Post-hoc analyses examining one possible reason for low within-session reliability examined the degree of within-session change relative to between-session change. This was done in SPSS using paired t-test analyses at the contrast level to compare the absolute value of between-session change with the absolute value of within-session change to examine if within-session change could exceed developmental change over a two-year period. Results significant after a multiple comparison correction for the number of contrasts were reported. Analyses were carried out using the outlier removed (QC+OR) sample.

Stability of change was calculated to examine if individual change metrics were stable over time. This was accomplished by calculating change within-session (run 2 – run 1) at baseline and follow-up and calculating stability using these two change statistics. To examine if the degree of change between-session was reliable, baseline and follow-up change at the individual level was calculated for each run (run 1 at follow-up – run 1 at baseline, run 2 at follow-up – run 2 at baseline) and used to calculate stability. These stability of change scores were correlated with each other, with the regular ICCs, with the absolute value of activity at baseline and follow-up (using Cohen’s D effect size values), and with the absolute value of change, within- and between-session (using effect size values). Results significant after a multiple comparison correction for the number of contrasts were reported. Analyses were carried out using the outlier removed (QC+OR) sample.

*Behavioral Differences Between Session and Possible Effects on Reliability and Stability*

Post-hoc analyses comparing between-session changes in behavior were calculated for variables that may contribute to differences in reliability and stability. What task specific behavioral tests were analyzed was guided by the outcomes of the ICC differences analyses. These included movement, the rate of incorrect stop and incorrect go trials in the SST, the rate of positive and negative reward and loss outcomes in the MID, and accuracy on the 0-Back and 2-Back for the nBack. Within-session change was also analyzed for movement and the SST performance variables. Movement is a source of noise in fMRI research (Bright and Murphy, 2017; Diedrichsen and Shadmehr, 2005) and fewer trial outcomes may result in not enough data being present to accurately model functional responses. Movement analyses were performed on run and session specific frame-wise displacement values from subjects who passed the task’s imaging quality control measure. Similarly, only subjects who passed the task’s behavioral quality control were analyzed. Differences in behavior were calculated as paired t-tests between run or session (run/session 2 > run/session 1), values significant after an FDR correction for multiple comparisons were reported as Cohen’s D values. D values for movement differences were averaged across task.

*Intersession Interval*

To measure the effect of intersession interval on stability, subjects were separated into quartiles based on the time between scans, with the average difference between quartiles of four months. Analyses were repeated using deciles to increase the intersession range to six months. The reliabilities and stabilities for 1st and 4th quartile and 1st and 10th decile were compared in contrast specific paired t-tests at the whole brain level. Results significant after an FDR correction for the number of contrasts were reported as Cohen’s D values and mean differences. Intersession interval separation used the QC+OR sample.

*Condition vs Baseline Contrast Correlations*

Previous research has found that condition vs baseline activity can be highly reliable, but similarity between individual’s activity across different condition vs baseline contrasts can result in poor reliability for the related condition vs condition contrast (Infantolino et al., 2018). To examine how an individual’s regional condition vs baseline activity was similar across condition vs baseline contrasts, activity from each region was correlated (Pearson) with activity from the same region in related condition vs baseline contrasts (face vs baseline and place vs baseline, 0-back vs baseline and 2-back vs baseline) for each session. These correlations were then correlated with the relevant within-session condition vs condition reliability to examine how similarity across condition vs baseline contrasts related to reliability. This was done for ROIs only and at the whole brain level. Correlations in ROIs were also compared against non-ROIs in independent sample t-tests. All analyses used the QC+OR sample.

**Supplemental Results**

*Association Between ICCs and Task-Related Activation, Age-Related Change, Region Volume, and other Types of ICCs*

*Do regions with greater task-related activation have higher ICCs?*

Reliability and stability were significantly correlated with activation level and the amount of change within- and between-session in most contrasts, with this relationship least likely to be found when examining change within-session. Significant correlations were all positive except for the correlation between activity and within-session reliability at baseline in the positive vs neutral face contrast. Across contrasts and ICCs, the average correlation between activity level and ICC was 0.42 (excluding 6 non-significant correlations, out of a total of 104 analyses). As occipital regions tended to be relatively highly reliable and active and of little interest, post-hoc analyses were performed after removing these regions to see if they were driving these correlations. Correlations remained largely the same. See Supplementary Table 11 for more information.

*Do regions with greater age-related changes show lower ICCs?*

Across contrasts that showed significant correlations, the average correlation of ICCs with the between-session change was 0.32 and with the within-session change was 0.36.

*Do larger regions show higher ICCs?*

Region volume did not predict reliability or stability with the exception of a few weak but significant relationships for the nBack and SST task (range of -0.07 to -0.09).

*Does within-session reliability predict longitudinal stability?*

Reliability within-session at baseline and follow-up and longitudinal stability were significantly correlated with each other in most contrasts (except the MID’s anticipation of large vs small loss and most of the emotion contrasts of the nBack), with a high of 0.98 for the correlation between the within-session reliabilities for the 0-back vs baseline contrast. The average correlation (significant correlations only) between reliabilities and stabilities was 0.78. See Supplemental Table 11 for a contrast specific breakdown of reliability and stability’s association with activity and change and Supplemental Table 9 for contrast specific correlations of ICCs with ICCs.

*Effects of Data Cleaning*

[Insert Supplementary Figure 4]

Small, though significant, differences were found between the ICCs of the different data cleaning approaches (quality control, quality control with outlier removal, quality control with movement regression and outlier removal, and quality control with rank normalization) types. Across contrasts, the greatest difference following data cleaning was observed in the within follow-up session reliability of the 2-back vs baseline contrast, where removing outliers increased the average reliability by 0.133, from 0.289 to 0.422). The average gain in ICC (across within-session reliabilities and longitudinal stabilities, contrasts with significant differences only) from removing outliers was 0.027, from rank normalizing data was also 0.027, and from regressing out movement and removing outliers was 0.029. Out of the 26 contrasts provided by the ABCD 3.0 release, significant gains in stability were observed after removing outliers in 19 contrasts, while gains in reliability were observed 23 contrasts within-session at baseline, and 22 within at follow-up. Regressing out movement and removing outliers increased stability in 17 contrasts, and increased reliability in 22 contrasts within-session at baseline and 21 within-session at follow-up, and significantly decreased within-session reliability at baseline and follow-up in the MID’s anticipation of large vs small loss. Rank normalizing data increased stability in 17 contrasts between-session and increased reliability in 21 contrasts within-session at baseline and 23 within at follow-up. For a comparison of reliabilities and stability from the nBack’s 2 vs 0-back contrast after removing outliers, see Figure 5, Figure 4A, and for a contrast specific comparison of the different data cleaning approaches, see Supplemental Table 4.

*Occipital vs Non-Occipital Comparisons*

Average reliability, stability, activity, and change (the regional absolute value of change within- and between-session) were significantly higher in occipital (regions with ‘oc’ or ‘occipital’ in their name as well as the calcarine and cuneus) relative to non-occipital brain regions in the majority of contrasts, with change between-session the measure with the fewest significant differences. A contrast specific comparison of occipital and non-occipital reliability, stability, activity, and change statistics (See Supplemental Table 5 for a contrast specific comparison) shows that among contrast with significant differences, occipital regions have ICCs an average of 0.08 higher and the Cohen’s D effect sizes of activation is on average 0.23 larger. These effects are most prominent in the condition vs baseline contrasts of the nBack, but are significant in most other contrasts as well. Within- and between-session change differences were also observed, though the average differences in effect sizes of change was very small, only 0.03.

*Reliability and Stability in ‘Condition vs Baseline’ vs ‘Condition vs Condition’ Contrasts*

ICCs were significantly higher in condition vs baseline contrasts relative to condition vs condition contrasts, with an average Cohen’s D effect size of 1.913, corresponding to a very large effect size. Reliability and stability patterns were significantly correlated in most comparisons except for the 0-back vs 2 vs 0-back within-session at baseline and follow-up.

*Sex Differences*

As sex impacts developmental slopes in adolescence (Gur and Gur, 2016), sex specific analyses were performed to examine Significant sex differences in reliability and stability were observed in most contrasts, though the difference was generally low, with the average difference (significant contrasts, absolute value of the mean difference) between-session was 0.014, within at baseline was 0.021, and within at follow-up was 0.024. The greatest contrast-wide difference between the sexes was observed in the within-session reliability at baseline in the anticipation of large vs small loss, where males had an average increase in reliability over females of 0.0614. Significant differences in stability between sexes contrast-wise were observed in 16 contrasts, favoring females in 11 contrasts, in 21 contrasts within-session at baseline, favoring females in 12, and 15 contrasts within-session at follow-up, favoring females in 6. There was little commonality across reliability/stability sex comparisons in what contrasts were significantly different in males and females and in which sex had the higher ICC. The greatest parcel-specific difference in stability between sexes was in the left superior parietal gyrus in the incorrect stop vs correct go contrast, where females had a 0.147 increase in stability over males. Within-session, the greatest parcelwise sex difference in reliability was observed at follow-up in the left temporal pole of the anticipation of loss vs neutral contrast, where reliability in males was 0.262 higher. Dividing the sample by sex, calculating ICC per sex, then averaging across contrasts did not significantly increase stability between-session in most contrasts, which would occur if differences in developmental trajectories between males and females were shifting individual’s relative positions, decreasing consistency-based stability. Significant differences between ICCs when calculated using all participants and when calculated by sex then averaged were found in 19 contrasts, with contrasts involving reward processing in the MID and emotion/face processing in the nBack least likely to be significant. Greater ICCs when averaging sex specific values were observed in only anticipation of large vs small reward and in incorrect go vs incorrect stop. The difference between the all-subject ICC and sex specific averaged ICC was very small, with the greatest difference being a 0.0041 average increase in the all-subjects ICC relative to the sex averaged ICC in the place vs baseline contrast of the emotional nBack. Similar patterns were observed in the within-session reliability comparisons. For more information, see Supplementary Table 12 and the BALSA archive.

*Comparison of Longitudinal Stabilities by Runs*

Longitudinal stability changed depending on the number of runs being analyzed. Stability based on the first run was higher than stability based on both runs in any MID contrast involving loss anticipation and any nBack contrast comparing emotional and neutral faces. There were no significant differences between using the full data and only the first run in the comparison of faces vs places in the nBack and the anticipation of large reward vs neutral and anticipation of reward vs neutral in the MID. Comparison of between-session stability using only run 1 and only run 2 found that stabilities from run 1 were higher or not significantly different than stabilities from run 2 in most contrasts except for the SST’s correct stop vs incorrect stop, incorrect go vs correct go, and incorrect go vs incorrect stop. Stability when using both runs was significantly higher or not significantly different than stability based solely on run 2 for all contrasts. See Supplemental Table 13 for more information.

*Longitudinal Stability of and Association of Change Within- and Between Session*

[Insert Supplementary Figure 8]

Post-hoc analyses investigating one possible reason for the low reliability and stability examined the degree of change within- and between-session (change due to habituation, task reorganization, automation, or other factors and longitudinal change, respectively) and the stability of change. Within- and between-session differences in activity were observed in the majority of contrasts and mostly took the form of decreases in activity from baseline to follow-up. Within-session changes were slightly larger and showed both increases and decreases in activity. Meta-analysis guided task-relevant regions analyses of 8 contrasts demonstrated that task-relevant regions exhibited significantly greater change than task-irrelevant regions between-session in only the 2 vs 0-back and incorrect stop vs correct go contrasts, with no differences in change within-session. While the average change in activity within time point was small (average absolute value of change across all contrasts within-session at time 1 and time 2 = 0.042 Cohen’s D), this was still larger than the average change between-session (average between-session change = 0.033), with paired comparison analyses finding a small but significant effect when comparing the amount of change within- and between-session (|change between| > |change within at baseline or follow-up| Cohen’s D at time 1 = -0.197, at time 2 = -0.2). When examined at the contrast level, similar patterns emerged, though greater or no significant differences for change between vs within-session was observed in the contrast of anticipation of large and small reward or loss values in the MID and in the working memory and condition vs baseline contrasts of the nBack. See Supplementary Table 14 and Supplementary Figure 5 for a contrast specific examination of change differences and an example of an nBack contrast with significant change observed within- and between-session and another nBack contrast with change only within-session. Changes in activity, shown for surfaces and subcortical slices for all change measures, can be found on BALSA at [URL].

The stability of change, within or between-session, was poor for all regions. The highest stability for within-session change was observed in the negative vs neutral face contrast of the emotional nBack in the left posterior collateral transverse sulcus (ICC = 0.118) and the highest stability for between-session change was seen in the same region in the 2-Back vs baseline contrast (0.391). The average stability across all contrasts was 0.013 for within-session change and 0.034 for between-session change, with a high average stability of within-session change of 0.039 for the negative vs neutral face contrast and a low of -0.013 in the loss feedback contrast, the contrast with the highest average between-session stability was the 2-back vs baseline (0.132) and the lowest was positive vs neutral faces (-0.009). Contrast-wise correlations between change stability and other ICCs, |activity|, and |change| statistics found that the stability of change within-session was significantly and positively correlated with the stability of change between-session (low of 0.243 for the 0-Back to a high of 0.612 for face vs place). Significant, and largely positive, correlations between the stability of within-session change and longitudinal stability was observed in 15 contrasts, with non-significant relationships observed for most MID contrasts, the 2 vs 0-back, and any contrast involving correct stop. Stability of change within-session was not significantly correlated with within-session reliability, the absolute value of activity, or the absolute value of between or within-session change for most contrasts, and for those it was, most were involved in the face and emotion processing contrasts of the nBack. The stability of between-session change was more strongly correlated with other statistical values relative to the stability of within-session change, with significant correlations for stability observed in 19 of 26 contrasts, ranging from a high of 0.821 for 0-Back, to a low of -0.348 for the anticipation of large vs small loss. The stability of between-session change was significantly and positively associated with within-session reliability in all contrasts except correct stop vs incorrect stop at baseline, with a high of 0.91 for the 0-Back at follow-up to a low of 0.215 for correct stop vs incorrect stop at follow-up. This measure was significantly and positively correlated with the absolute value of activity in 12 contrasts at baseline and 17 at follow-up, with significant contrasts coming from all tasks and no clear link between non-significant contrasts. Between-session change stability was not significantly associated with the absolute value of within or between-session change for most contrasts, and for the contrasts where there was a significant association, no clear pattern emerges across tasks or contrasts. For more information, see Supplementary Tables 15 and 16 and the BALSA archive.

*Change in Movement and Task Performance*

Significant differences in behavior that may impact reliability and stability were found for movement, MID, and SST performance. Movement significantly increased within-session (D = 0.537 at baseline, 0.460 at follow-up) and decreased between (D = -0.288). False alarm rate decreased between-sessions (D = -0.141) while incorrect stop rate increased (D = 0.158). Within-session, false alarms rate increased at baseline and follow-up (D = 0.203 and 0.181, respectively) as did incorrect stop (D = 0.405 and 0.366). The number of positive reward and loss outcomes increased (D = 0.142 and 0.195, respectively) while the number of negative reward and loss outcomes decreased (D = -0.137 and -0.189, respectively). 2-back and 0-back accuracy significantly increased between sessions (D = 0.9 and 0.57, respectively). All values are significant at a p < 0.001.

*Intersession Interval*

ICCs generally increased with longer intersession intervals. The differences between groups were small, with the average difference in longitudinal stability (only in contrasts where a significant effect was observed) was -0.019 for quartiles and -0.017 for deciles. While these differences seem negligible, the underlying stabilities are very weak and contrast specific differences can have Cohen’s D effect sizes that exceed 1, corresponding to large differences. Effect of intersession interval within session was larger for deciles (average for contrasts with significant differences = -0.028 for both sessions) relative to quartiles (-0.005).

*Condition vs Baseline Contrast Correlations*

Correlations between activity from different condition vs baseline contrasts were moderate. Mean (SD) pearson correlations across all regions for face vs baseline and place vs baseline = .527 (.060) for the baseline session and .525 (.072) at follow-up in ROIs. Average correlations were .534 (.083) for the baseline session and .534 (.088) at follow-up for whole brain analyses. For the 0-back vs baseline and 2-back vs baseline comparisons, mean (SD) correlations for ROIs were .635 (.038) for the baseline session and .616 (.047) at follow-up. Whole brain mean (SD) correlations were .646 (.084) for the baseline session and .629 (.088) at follow-up. Correlations were not significantly higher in ROIs relative to non-ROIs. Face vs place correlations were not significantly correlated with within-session reliability at the ROI level, presumably due to the few ROIs (20 for face vs place and 7 for 2 vs 0-back) but were at the whole brain level for face vs place (r = .462 at baseline, .359 at follow-up). This result is conceptually not consistent with Infantolino et al. (2018), who reported that the high correlation between the amygdala response to faces and shapes (each to implicit baseline) was accompanied by poor consistency of the differential response between faces and shapes. These results indicate that across the brain as a whole, similarity of condition vs baseline activity did not have a consistent relationship to the reliability of a corresponding condition vs condition contrast.

**Supplemental Discussion**

*Effect of Data Cleaning Comparison*

ICCs were calculated after different data cleaning procedures were applied to measure the effect of cleaning. Different approaches to cleaning the data (outlier removal, movement regression, rank ordering) did significantly increase ICCs, however this increase was small, increasing ICCs an average of 0.028 across the different approaches. Comparisons of the different data cleaning approaches do not show one approach is consistently better than any other. As the rank ordering and movement regression approaches tested here mean center results, making cross session comparisons useless, the outlier removal approach is recommended when analyzing ABCD data. Note that these are only a few data cleaning approaches, other approaches (e.g. performance filtering, multivariate outlier removal) may produce better results.

*Occipital vs Non-Occipital Comparisons*

Higher reliability, stability, activity, and change statistics were observed in occipital regions in most contrasts, prompting post-hoc analyses to quantify these differences. They found that these measures were indeed significantly greater in occipital regions relative to the rest of the brain in most contrasts. This is unfortunate as the regions that are most stable are generally not considered relevant to the developmental adolescent substance use and psychopathology research and are likely of little use.

*ICCs in Condition vs Baseline vs Condition vs Condition Contrasts*

Condition vs baseline ICCs were significantly higher than condition vs condition ICCs, though the vs baseline contrasts may not necessarily be desirable. The vs baseline contrasts are not isolating a particular effect so we cannot be certain what aspect of activity is reliable, making any inferences formed from analyzing them questionable.

*Development and Sex*

We examined the effect that separating the sample by sex would have on stability as sex contributes to differences in development (Gur and Gur, 2016), a factor that would change relative rankings and lower ICCs. These analyses were done in an attempt to partially mitigate differences in development that would lower ICCs as more advanced statistical measures capable of accounting for nonlinear slopes (e.g. latent growth curve analysis; Herting et al., 2018) require more sessions of data than are currently available. Paired t-tests found significant differences in activity in most contrasts between sessions, showing that activity is subject to developmental change. While development itself is not the focus of these analyses, it can influence stability. Developmental change is expected in this late childhood-early adolescence sample, yet change by itself doesn’t necessarily mean poor stability as long as individuals remain in relatively the same position. Heterogeneity in change may be expected depending on sample and time interval between-session, and one possible source, developmental sex differences, was addressed. We found stabilities differed by sex, though did not consistently favoring males or females. Averaging the per sex stabilities, which would eliminate the role of sex differences in developmental onset and trajectory, and comparing them to the stabilities calculated using all subjects did not show an increase in stability and was in fact slightly weaker, with an average ICC (in contrasts with significant differences) 0.001 lower. Out of the 65,000+ variables reported in the ABCD project, some could conceivably affect developmental change in a way that would reduce stability, however these analyses suggest that sex by itself is not one of them. Analyses looking at the interaction of sex, pubertal hormone levels, pubertal stage, etc., or variables completely unrelated to sex may identify what could cause heterogeneous change that would reduce stability. While developmental differences may account for poor between-session stability, they would not account for the poor within-session reliability, though habituation, sensitization, and functional reorganization might.

*Within-Session Change*

The sequential nature of the scans in the within-session reliability analyses may mean they are subject to change, be it through habituation, sensitization, or shifting functional patterns as the task loses novelty and becomes automated. The inclusion of a mock scanner session, where participants were tested for movement and familiarized with the tasks prior to actual scanning, means some of these factors may already be present in the first run. Comparison of the absolute value of change within- and between-session found that for most contrasts, there was greater change due to these factors than development. Significantly greater between-session change relative to within-session change was consistently observed only in the comparison of large and small reward anticipation, the anticipation of small reward vs neutral, and the comparison of the 2 vs 0-back. Analyses of the stability of this change found very little consistency in individual change patterns, even though group level effects found significant differences. Furthermore, analyses of run specific stabilities found that in some contrasts stability was not significantly different with only one run of data, and even significantly increased with less data. This effect was most prominent in the nBack’s face and emotion processing contrasts, which have been previously demonstrated to be subject to habituation (Plichta et al., 2014), and in the anticipation contrasts of the MID, suggesting that researchers interested in this task may want to analyze runs separately or examine the change between runs as an individual difference measure, though the poor stability in change may make this untenable. Average stability calculated using only the first run of data was higher than stability calculated using only the second in most contrasts except for the SST contrasts detailed under the movement section, though this could be due to the increase in movement in the second relative to first fun of each session. These results are indicative of within-session changes effects in the ABCD task data that could reduce reliability. They also indicate that increasing run number or length may do little to increase stability for the MID anticipation and nBack emotion tasks. The increase in incorrect responses in the second run of the SST and the accompanying increase in stability indicates that increasing available data may increase stability, but we do not see evidence for that outside of these incorrect trials for the SST. Examining within-session change in task activity with only two sessions of data is unlikely to capture the full extent of within-session change, however, with reprocessing of the raw data provided by ABCD it may be possible to examine the change in activity on a trial by trial or block by block basis.

*Intersession Interval*

Comparison of stabilities by low vs high intersession interval quartiles and deciles found that stability increased with longer intervals. This is surprising as, if developmental differences are responsible for low longitudinal stability, stabilities would increase with shorter intervals. It may be that the difference between quartiles or deciles of four or six months doesn’t matter relative to the overall two year gap between scanning sessions, however, group differences were observed within session at baseline, when no effect of varying intersession interval should be present. It may be the case that with such low overall reliability, small, intergroup deviations appear significant, even if they are just the result of small differences in group selection.

*Condition vs Baseline Contrast Correlations*

Regional activity in condition vs baseline contrasts was moderately related to activity in the same region in another condition vs baseline contrast and positively correlated with within-session reliability, but only for the face vs place contrast and related condition vs baseline contrasts. These analyses partially replicate findings from Infantolino et al. (2018), who found that condition vs baseline reliability in the amygdala (average of left and right activity) was highly reliable for face (.97) and shape (.95) condition vs baseline activity, but not for the contrast of faces and shapes (-.06), due to a high (.97) correlation between condition vs baseline activities. For contrast, the average correlation (across hemispheres and sessions) between activity in the amygdala for the face vs baseline and place vs baseline contrasts using ABCD data was .428 and the average amygdala within-session reliability (using the QC+OR sample) was .194 for 0-back vs baseline, .246 for 2-back vs baseline, and .082 for 2 vs 0-back. Differences in reliability between ABCD and Infantolino et al. (2018) may be the result of different designs (face vs place as part of an nBack task in ABCD and face vs shape as part of an emotion matching task for Infantolino et al. (2018)) or due to how reliability was calculated (comparison of separate runs for ABCD and a comparison of alternating blocks in Infantolino et al. (2018), the first approach would be subject to habituation effects (Plichta et al., 2014) while the second would not be to the same extent). Infantolino et al.’s (2018) assertion that the high correlation between condition vs baseline contrasts as being responsible for the poor reliability of condition vs condition activity is somewhat undercut by our findings of a positive relationship between reliability and condition vs baseline contrasts, however the relative scale of the conditions vs baseline correlations in our analyses (~.5-.6) versus Infantolino et al’s (.97; 2018) mean that individual differences in activity would be greater, allowing for more variable individual rank orderings.

**Supplemental Figure 1**

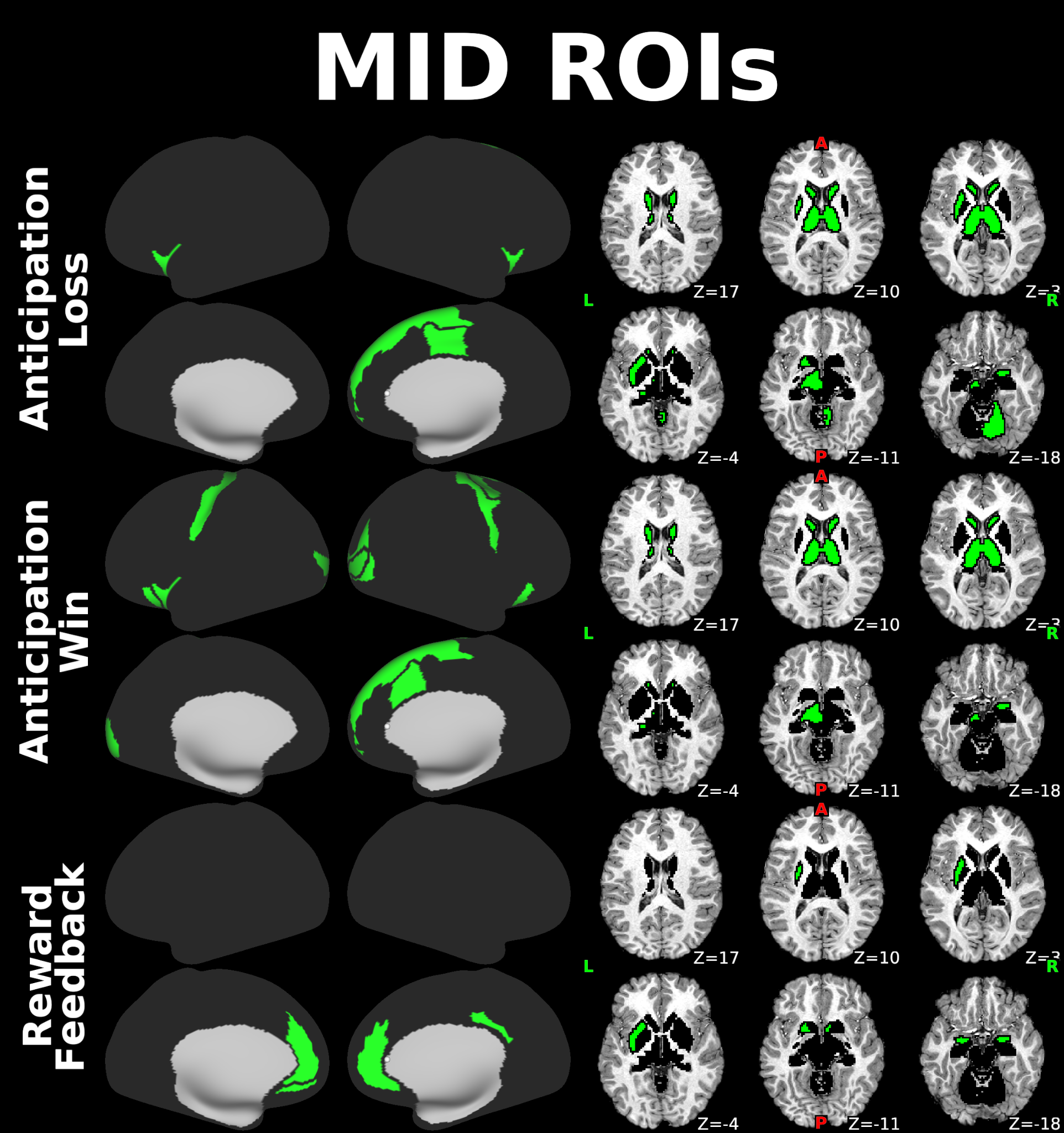

Regions of interest for the Monetary Incentive Delay task, as derived from Oldham et al. (2018), shown in green.

**Supplemental Figure 2**

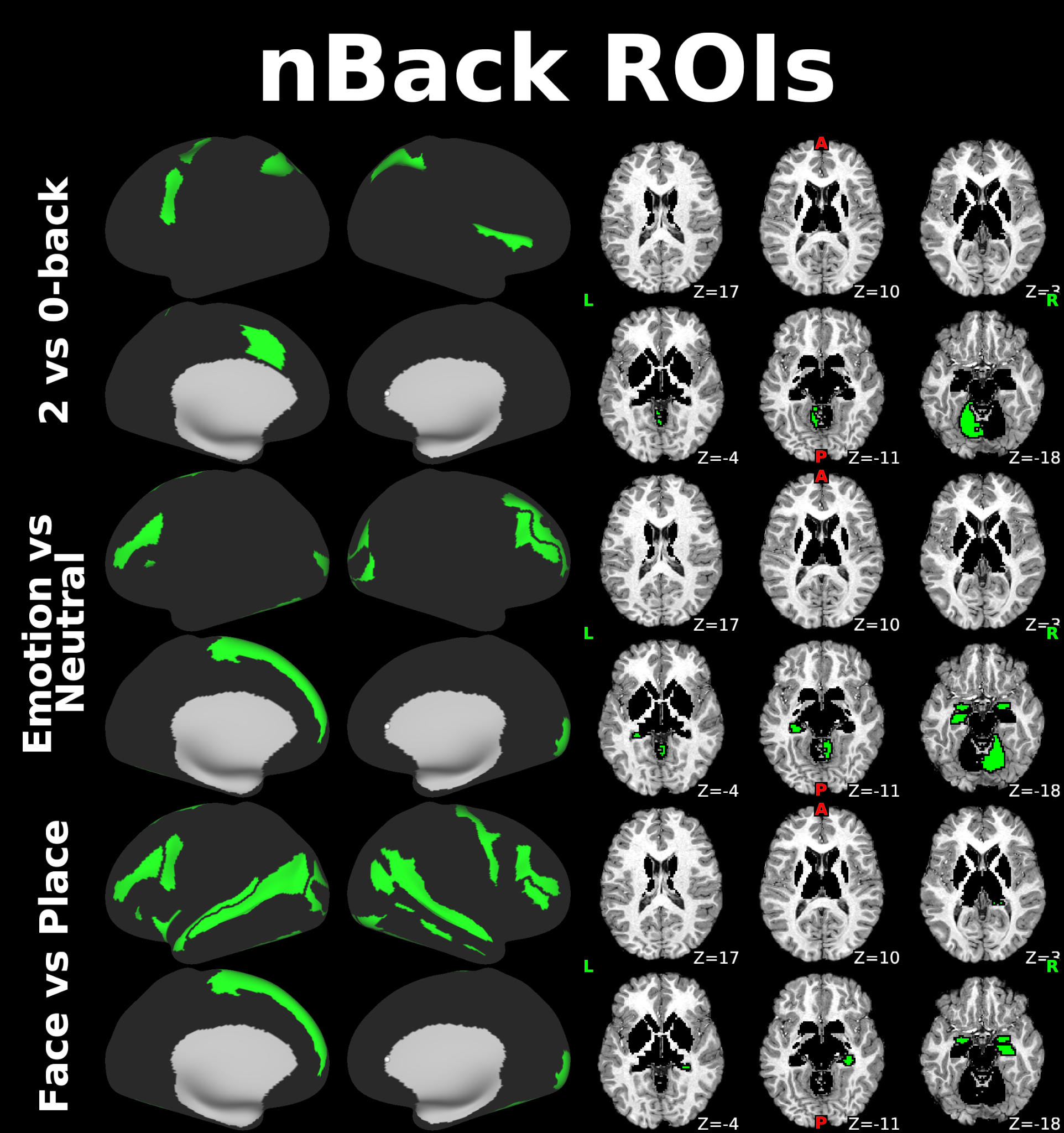

Regions of interest for the emotional nBack task shown in green, based on meta-analyses by Yaple et al. (2018) for 2 vs 0-back contrast and Muller et al. (2018) for the emotion vs neutral and face vs place contrasts.

**Supplemental Figure 3**

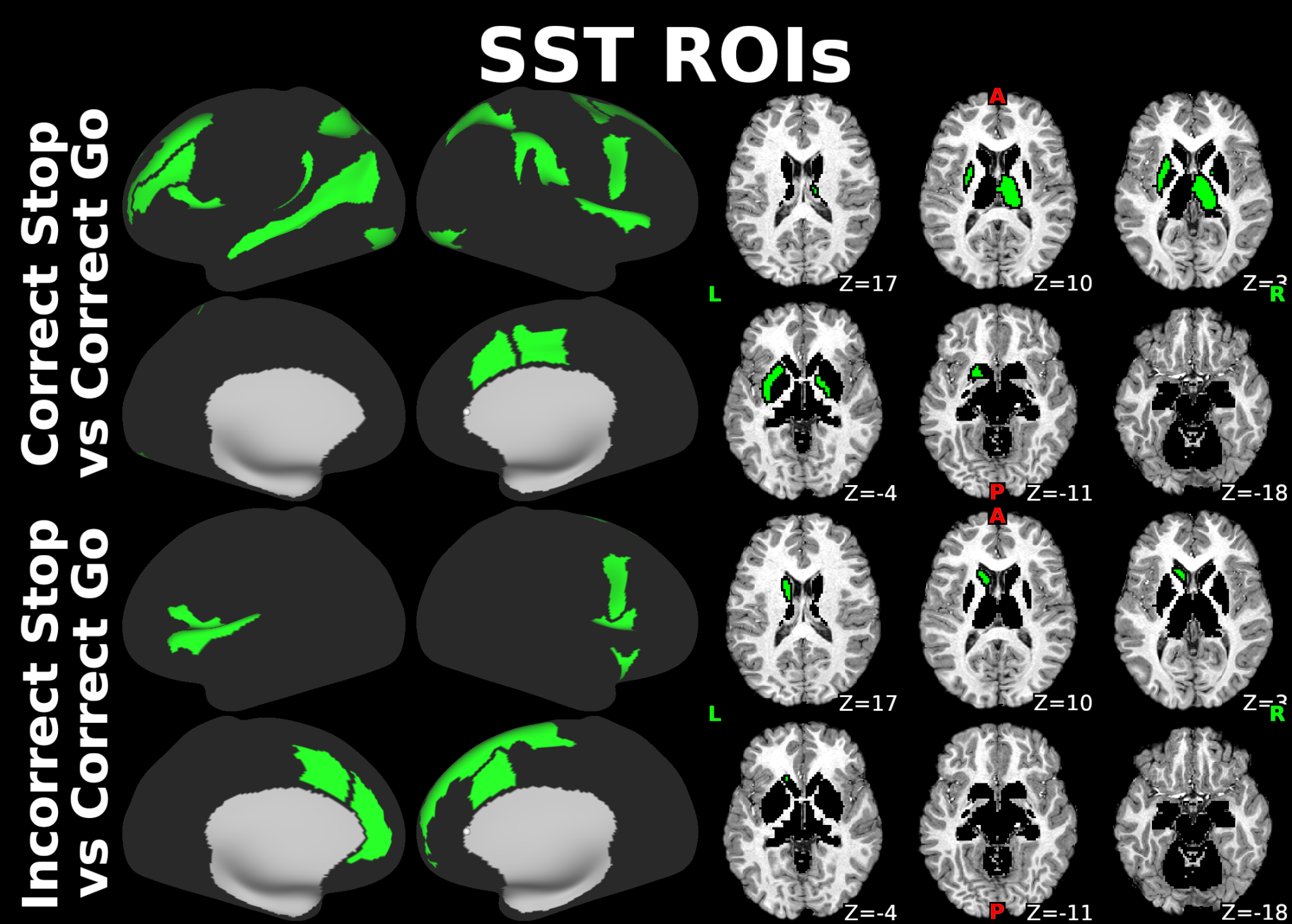

Regions of interest for the Stop Signal Task shown in green, based on meta-analyses by Swick et al. (2011) for correct stop vs correct go contrast and Neta et al. (2015) for the incorrect stop vs correct go contrast.

**Supplemental Figure 4**

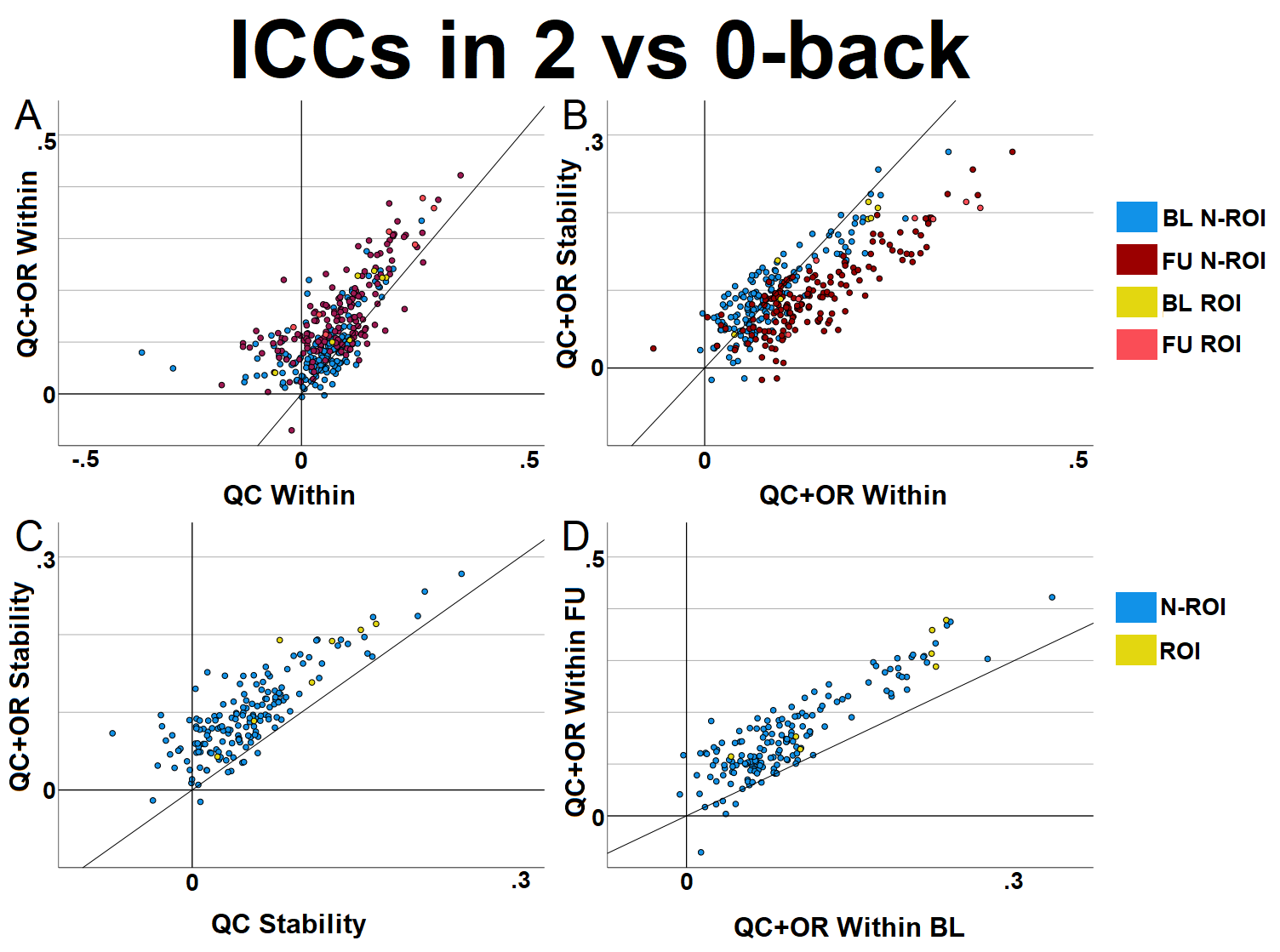

Scatter plots from the 2 vs 0 Back comparing (A) the within session reliability when using QCed data (QC) and outlier removed data (QC+OR), (B) Stability vs reliability from baseline and follow-up (QC+OR analyses), and (C) Stability when using QCed data (QC) and outlier removed data (QC+OR), and (D) Within-session reliability at baseline compared to follow-up (QC+OR analyses). BL: Baseline, FU: Follow-Up, N-ROI: Non-ROI regions.

**Supplemental Figure 5**

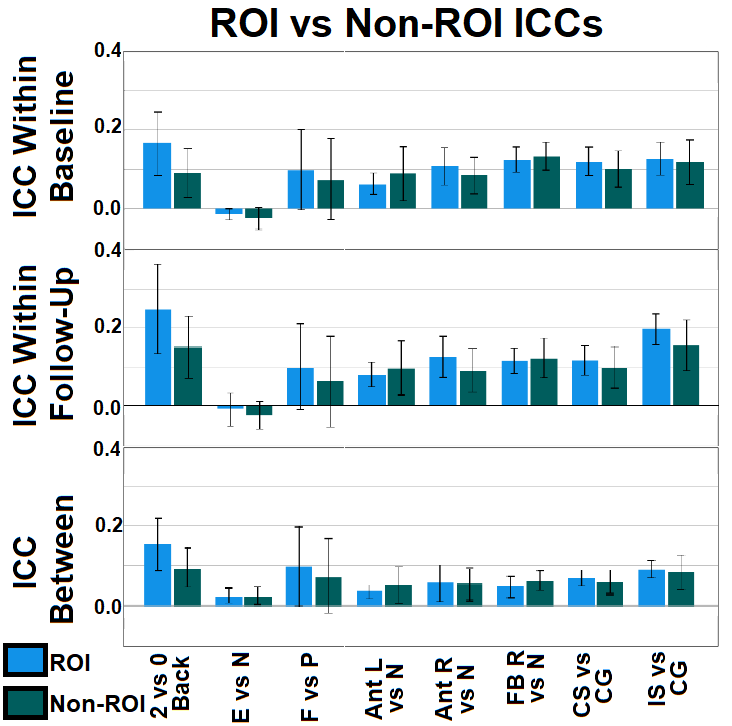

Average ICCs of beta values in task-relevant (Green) and task-irrelevant regions (Blue). E – Emotion, N – Neutral, F – Face, P – place, ANT – Anticipation, FB – Feedback, R – Reward, L – Loss, CS – Correct Stop, CG – Correct Go, IS – Incorrect Stop, W – Within-session. Bars – 1 standard deviation.

**Supplemental Figure 6**

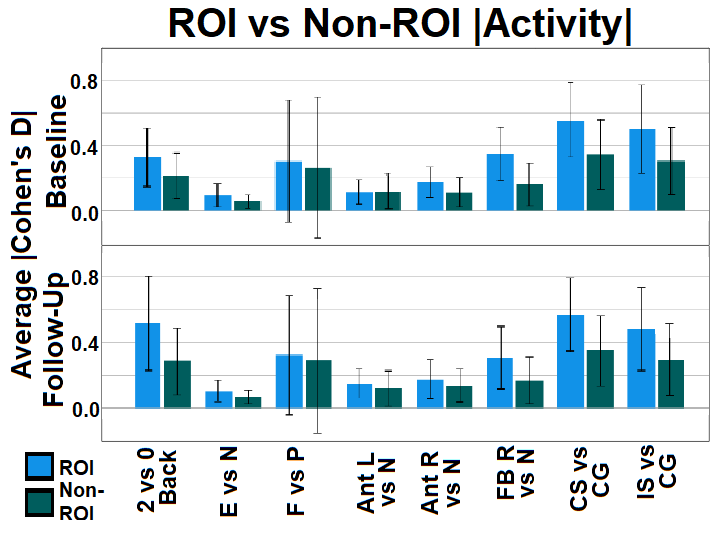

Average absolute values of effect sizes (Cohen’s D from one-sample comparison against 0) for activation data in task-relevant regions (Green) and task-irrelevant regions (Blue). E – Emotion, N – Neutral, F – Face, P – place, ANT – Anticipation, FB – Feedback, R – Reward, L – Loss, CS – Correct Stop, CG – Correct Go, IS – Incorrect Stop. Bars – 1 standard deviation.

**Supplemental Figure 7**

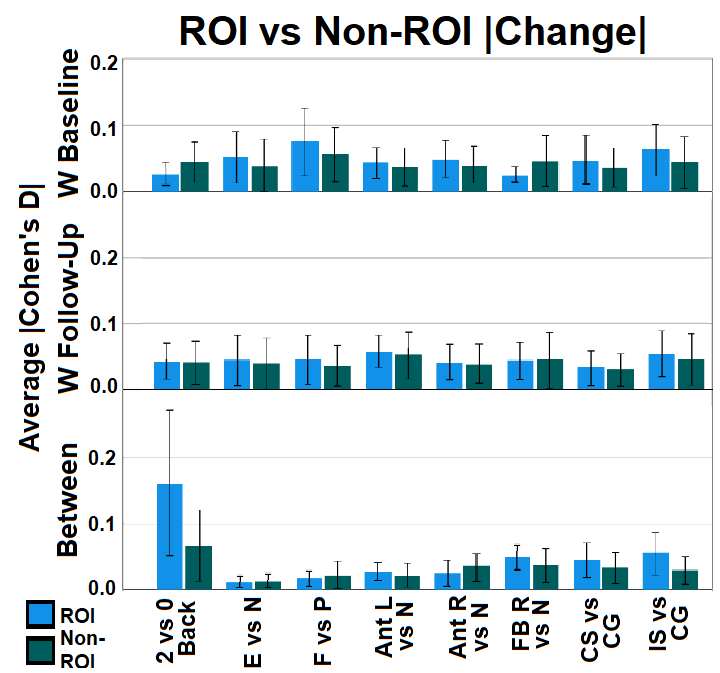

Average absolute values of within and between-session change effect sizes (Cohen’s D of paired comparison) in task-relevant regions (Green) and task-irrelevant regions (Blue). E – Emotion, N – Neutral, F – Face, P – place, Ant – Anticipation, FB – Feedback, R – Reward, L – Loss, CS – Correct Stop, CG – Correct Go, IS – Incorrect Stop, W – Within-session. Bars – 1 standard deviation.

**Supplemental Figure 8**

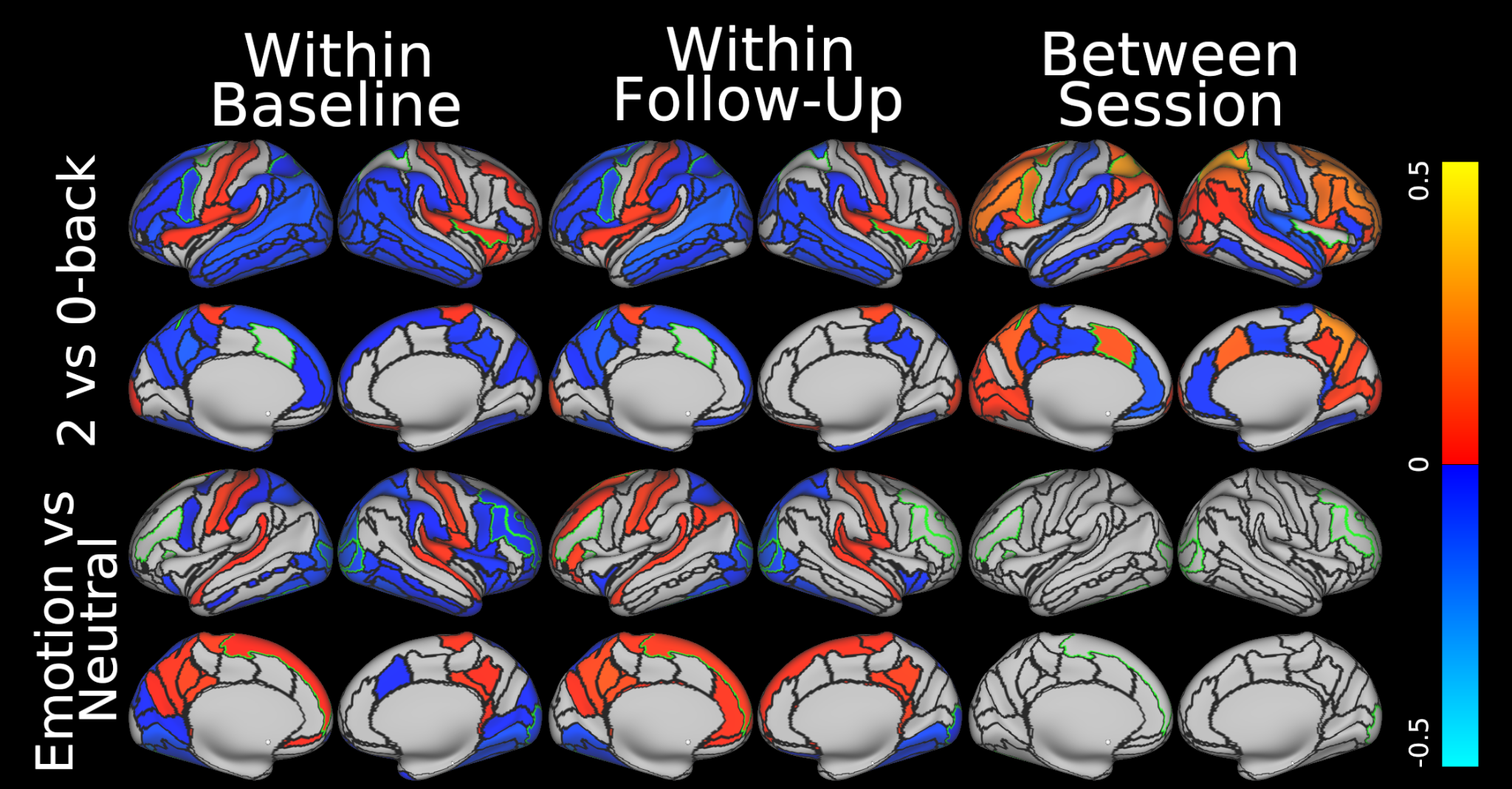

Change within (run 2 - run 1) and between-session for the 2 vs 0-back and Emotion vs Neutral contrast from the nBack task. Effect sizes of change shown, gray indicates not significant after an FDR multiple comparison correction. Red-Yellow indicates activity increases, Blue indicates activity decreases. Significant change only observed within-session in the Emotion vs Neutral task.

1. Within-session reliability for each session was correlated with the activity and within-session change of the same session.
