## Supplementary material for "Reliability and Stability Challenges in ABCD Task fMRI Data": Code for generating data and results: Script 1 - DataPrep.R

```
#ICC data prep

setwd("C:/Users/James/Desktop/Release 3 Nov 2020")#Set to ABCD text release folder

#Copy and rename task beta files to a new folder
dir.create('./ICC/')
taskbeta<-c('abcd_midabwdp01','abcd_midabwdp202','abcd_midr1bwdp101','abcd_midr1bwdp202','abcd_midr1bwp102','abcd_midr1bwp202','abcd_tfabwdp101',
            'abcd_tfabwdp201','abcd_tfnbr2bwdp101','abcd_tfnbr2bwdp201','abcd_tfncr1bwdp101','abcd_tfncr1bwdp201',
            'abcd_tfsstabwdp101','abcd_tfsstabwdp201','abcd_tfsstr1bwdp101','abcd_tfsstr1bwdp201','abcd_tfsstr2bwdp101','abcd_tfsstr2bwdp201','abcd_tr2bwdp01',
            'abcd_tr2bwdp202','midaparc03','midaparcp203','midr2bwp102','midr2bwp202','mrisst02','mrisstr1bw01','mrisstr2bw01','nback_bwroi02','nbackr101',
            'nbackr201','abcd_imgincl01')
for(i in taskbeta){
  file.copy(from = paste0('./',i,'.txt'), to = './ICC/')
}

funcrename<-read.csv('funcrename.csv')#This file renames variables based on the '21.abcd_3.0_mapping.csv' deap names, can't create it due to unrecognized symbols stopping r from reading the whole file
setwd('./ICC/')

file.rename(from = './abcd_midabwdp01.txt', to = './MID_Avg_Beta_Dx_1.txt')
file.rename(from = './abcd_midabwdp202.txt', to = './MID_Avg_Beta_Dx_2.txt')
file.rename(from = './abcd_midr1bwdp101.txt', to = './MID_R1_Beta_Dx_1.txt')
file.rename(from = './abcd_midr1bwdp202.txt', to = './MID_R1_Beta_Dx_2.txt')
file.rename(from = './abcd_midr1bwp102.txt', to = './MID_R1_Beta_ASDS_1.txt')
file.rename(from = './abcd_midr1bwp202.txt', to = './MID_R1_Beta_ASDS_2.txt')
file.rename(from = './abcd_tfabwdp101.txt', to = './nBack_Avg_Beta_Dx_1.txt')
file.rename(from = './abcd_tfabwdp201.txt', to = './nBack_Avg_Beta_Dx_2.txt')
file.rename(from = './abcd_tfnbr2bwdp101.txt', to = './nBack_R2_Beta_Dx_1.txt')
file.rename(from = './abcd_tfnbr2bwdp201.txt', to = './nBack_R2_Beta_Dx_2.txt')
file.rename(from = './abcd_tfncr1bwdp101.txt', to = './nBack_R1_Beta_Dx_1.txt')
file.rename(from = './abcd_tfncr1bwdp201.txt', to = './nBack_R1_Beta_Dx_2.txt')
file.rename(from = './abcd_tfsstabwdp101.txt', to = './SST_Avg_Beta_Dx_1.txt')
file.rename(from = './abcd_tfsstabwdp201.txt', to = './SST_Avg_Beta_Dx_2.txt')
file.rename(from = './abcd_tfsstr1bwdp101.txt', to = './SST_R1_Beta_Dx_1.txt')
file.rename(from = './abcd_tfsstr1bwdp201.txt', to = './SST_R1_Beta_Dx_2.txt')
file.rename(from = './abcd_tfsstr2bwdp101.txt', to = './SST_R2_Beta_Dx_1.txt')
file.rename(from = './abcd_tfsstr2bwdp201.txt', to = './SST_R2_Beta_Dx_2.txt')
file.rename(from = './abcd_tr2bwdp01.txt', to = './MID_R2_Beta_Dx_1.txt')
file.rename(from = './abcd_tr2bwdp202.txt', to = './MID_R2_Beta_Dx_2.txt')
file.rename(from = './midaparc03.txt', to = './MID_Avg_Beta_ASDS_1.txt')
file.rename(from = './midaparcp203.txt', to = './MID_Avg_Beta_ASDS_2.txt')
file.rename(from = './midr2bwp102.txt', to = './MID_R2_Beta_ASDS_1.txt')
file.rename(from = './midr2bwp202.txt', to = './MID_R2_Beta_ASDS_2.txt')
file.rename(from = './mrisst02.txt', to = './SST_Avg_Beta_ASDS.txt')
file.rename(from = './mrisstr1bw01.txt', to = './SST_R1_Beta_ASDS.txt')
file.rename(from = './mrisstr2bw01.txt', to = './SST_R2_Beta_ASDS.txt')
file.rename(from = './nback_bwroi02.txt', to = './nBack_Avg_Beta_ASDS.txt')
file.rename(from = './nbackr101.txt', to = './nBack_R1_Beta_ASDS.txt')
file.rename(from = './nbackr201.txt', to = './nBack_R2_Beta_ASDS.txt')

#Make movement file
mvmtfiles<-list.files(pattern=".*.Beta_ASDS.*.")
mvmtfiles<-mvmtfiles[!grepl('.*._2.txt',mvmtfiles)]
mvmt<-as.data.frame(matrix(ncol=2,nrow=0))
names(mvmt)<-c('subjectkey','eventname')

for(i in mvmtfiles){
  abcddata<-read.table(i, sep='\t', header=T, stringsAsFactors = F)[-1,]
  abcddata<-abcddata[,which(names(abcddata) %in% c('subjectkey','eventname',names(abcddata)[grepl('meanmotion',names(abcddata))],
                                                   names(abcddata)[grepl('_mm',names(abcddata))]))]
  mvmt<-merge(mvmt,abcddata,by=c('subjectkey','eventname'),all=T)
}
rm(abcddata)
mvmtT1<-subset(mvmt,eventname=='baseline_year_1_arm_1')
mvmtT2<-subset(mvmt,eventname=='2_year_follow_up_y_arm_1')
mvmtT1<-mvmtT1[,-2]
mvmtT2<-mvmtT2[,-2]
names(mvmtT1)<-c('ID','MA_0m','M1_0m','M2_0m','NA_0m','N1_0m','N2_0m','SA_0m','S1_0m','S2_0m')
names(mvmtT2)<-c('ID','MA_24m','M1_24m','M2_24m','NA_24m','N1_24m','N2_24m','SA_24m','S1_24m','S2_24m')
mvmt<-merge(mvmtT1,mvmtT2,by='ID',all=T)
write.csv(mvmt,'mvmt.csv',row.names=F,na='')

#Select usable subjects, rename variables, split files by time point, merge output
imginclude<-read.table('abcd_imgincl01.txt',sep='\t',header=T)[-1,]
imginclude<-imginclude[,c(4,9,15,16,17)]
file.remove('abcd_imgincl01.txt')
usable<-list.files(pattern='.*.txt')
dir.create('OLD')
for(i in usable){
  abcddata<-read.table(i,sep='\t',header=T,stringsAsFactors = F,na='')[-1,]
  namesplit<-unlist(strsplit(i,"_"))
  if(namesplit[1]=="MID"){abcddata<-merge(imginclude[,1:3],abcddata,by=c('subjectkey','eventname'),all=T)}
  if(namesplit[1]=="nBack"){abcddata<-merge(imginclude[,c(1:2,4)],abcddata,by=c('subjectkey','eventname'),all=T)}
  if(namesplit[1]=="SST"){abcddata<-merge(imginclude[,c(1:2,5)],abcddata,by=c('subjectkey','eventname'),all=T)}
  names(abcddata)[3]<-'Usable'
  abcddata<-subset(abcddata,Usable=='1')
  abcddata<-abcddata[,-3]
  datanames<-names(abcddata)
  for(z in datanames){
    pairedval<-which(funcrename==z)
    if(length(pairedval)>0){
      datanames<-gsub(paste0("\\b",z,"\\b"),funcrename[pairedval,2],datanames)
    }
  }
  names(abcddata)<-datanames
  abcddata<-cbind(abcddata[,1:2],abcddata[,c(names(abcddata)[grepl('aseg',names(abcddata))],names(abcddata)[grepl('desikan',names(abcddata))],
                                         names(abcddata)[grepl('destrieux',names(abcddata))])])
  abcdT1<-subset(abcddata,eventname=='baseline_year_1_arm_1')
  abcdT2<-subset(abcddata,eventname=='2_year_follow_up_y_arm_1')
  names(abcdT1)<-paste0(names(abcdT1),"_0m")
  names(abcdT2)<-paste0(names(abcdT2),"_24m")
  abcdT1<-abcdT1[,-2]
  abcdT1<-abcdT1[complete.cases(abcdT1),]
  abcdT2<-abcdT2[,-2]
  abcdT2<-abcdT2[complete.cases(abcdT2),]
  names(abcdT1)[1]<-'ID'
  names(abcdT2)[1]<-'ID'
  abcddata<-merge(abcdT1,abcdT2,by='ID',all=T)
  abcddata <- Filter(function(x)!all(is.na(x)), abcddata)
  write.csv(abcddata,paste0('usable_renamed_',gsub('.txt','.csv',i)),na="",row.names=F)
  rm(abcdT1,abcdT2)
  file.rename(from=i,to=paste0('./OLD/',i))
}
rm(abcddata,imginclude)

tocombine1<-list.files(pattern='.*._1.csv')
for(i in tocombine1){
  part1<-read.csv(i)
  part2<-read.csv(gsub('_1.csv','_2.csv',i))
  combined<-merge(part1,part2,by='ID',all=T)
  write.csv(combined,gsub('_1.csv','.csv',i),na='',row.names=F)
  file.rename(from=i,to=paste0('./OLD/',i))
  file.rename(from=gsub('_1.csv','_2.csv',i),to=paste0('./OLD/',gsub('_1.csv','_2.csv',i)))
}
rm(part1,part2,combined,tocombine1)
tocombine2<-list.files(pattern='.*._R1_.*.csv')
for(i in tocombine2){
  run1<-read.csv(i)
  run2<-read.csv(gsub('_R1_','_R2_',i))
  R1R2<-merge(run1,run2,by='ID',all=T)
  write.csv(R1R2,gsub('_R1_','_R1R2_',i),row.names=F,na='')
  file.rename(from=i,to=paste0('./OLD/',i))
  file.rename(from=gsub('_R1_','_R2_',i),to=paste0('./OLD/',gsub('_R1_','_R2_',i)))
}
rm(list=ls())

#Create outlier removed data - somtimes randomly dies, rerunning the for loop doesn't necessarily create the same errors with the same files, editing pattern to be more selective can help move past error
toout<-list.files(pattern='^usable_renamed.*.csv$')
OutVal<-3
for(i in toout){
  func<-read.csv(i)  
  measurename<-gsub('usable','Outlier',i)
  Vars<-scale(func[,2:ncol(func)])
  while(length(which(Vars > abs(OutVal)))!=0){
    Vars[Vars > OutVal]<-NA
    Vars[Vars < -OutVal]<-NA
    Vars<-scale(Vars)
  }
  funcOut<-func[,2:ncol(func)]
  funcOut[is.na(Vars)]<-NA
  output<-as.data.frame(cbind(func[,1],funcOut))
  names(output)[1]<-'ID'
  write.csv(output,measurename,row.names = F,na='')
}
rm(list=ls())

#Create rank based data
if('RNOmni' %in% rownames(installed.packages())==FALSE){
  install.packages('RNOmni')}
library('RNOmni')

torank<-list.files(pattern='^usable_renamed.*.csv$')
for(i in torank){
  func<-read.csv(i)  
  measurename<-gsub('usable','Rank',i)
  VarsNorm<-as.data.frame(func[,1])
  IDVars<-as.data.frame(func[,1])
  Variables<-names(func)[-1]
  for(z in Variables){
    tmp<-(z)
    tmp<-subset(func, select = tmp)  #Rank norm command doesn't handle missing data well, this makes sure only subjects with usable data at this session are used
    tmpID<-cbind(IDVars,tmp)
    tmpID<-tmpID[complete.cases(tmpID[,2]),]
    tmpNorm<-RankNorm(unlist(tmpID[,2]))
    tmpID[,2]<-tmpNorm
    VarsNorm<-merge(VarsNorm,tmpID, by.x=colnames(VarsNorm)[1],by.y=colnames(tmpID)[1],all=T)
  }
  Vars<-merge(IDVars,VarsNorm,by.x = names(IDVars)[1], by.y = names(VarsNorm)[1],all = T)
  colnames(Vars)<-c(colnames(IDVars),Variables)
  names(Vars)[1]<-'ID'
  write.csv(Vars,measurename,row.names=F,na='')
}
rm(list=ls())

#Create movement regressed variables - separates previously created 'usable' files into each run by each time point and regresses out movement then removes outliers
mvmt<-read.csv('mvmt.csv')
abcd<-list.files(pattern='^usable_renamed_.*._Avg_Beta_.*.csv')

for(i in abcd){
  abcddata<-read.csv(i)
  abcdR1R2name<-gsub('Avg','R1R2',i)
  abcdR1R2<-read.csv(abcdR1R2name)
  namecomp<-unlist(strsplit(i,"_"))
  if(namecomp[3]=='MID'){task<-"MID"}
  if(namecomp[3]=='nBack'){task<-"nBack"}
  if(namecomp[3]=='SST'){task<-"SST"}
  T1Data<-cbind(abcddata[,1],abcddata[grepl('_0m',names(abcddata))])
  names(T1Data)[1]<-'ID'
  T2Data<-cbind(abcddata[,1],abcddata[grepl('_24m',names(abcddata))])
  names(T2Data)[1]<-'ID'
  T1R1R2<-cbind(abcdR1R2[,1],abcdR1R2[grepl('_0m',names(abcdR1R2))])
  names(T1R1R2)[1]<-'ID'
  T2R1R2<-cbind(abcdR1R2[,1],abcdR1R2[grepl('_24m',names(abcdR1R2))])
  names(T2R1R2)[1]<-'ID'
  T1R1Data<-cbind(T1R1R2[,1],T1R1R2[grepl('_run1_',names(T1R1R2))])
  names(T1R1Data)[1]<-'ID'
  T1R2Data<-cbind(T1R1R2[,1],T1R1R2[grepl('_run2_',names(T1R1R2))])
  names(T1R2Data)[1]<-'ID'
  
  T2R1Data<-cbind(T2R1R2[,1],T2R1R2[grepl('_run1_',names(T2R1R2))])
  names(T2R1Data)[1]<-'ID'
  T2R2Data<-cbind(T2R1R2[,1],T2R1R2[grepl('_run2_',names(T2R1R2))])
  names(T2R2Data)[1]<-'ID'
  
  if(task=="MID"){T1Data<-merge(mvmt[,1:2],T1Data,by='ID',all=T)
  T2Data<-merge(mvmt[,c(1,11)],T2Data,by='ID',all=T)
  T1R1Data<-merge(mvmt[,c(1,3)],T1R1Data,by='ID',all=T)            
  T2R1Data<-merge(mvmt[,c(1,12)],T2R1Data,by='ID',all=T)            
  T1R2Data<-merge(mvmt[,c(1,4)],T1R2Data,by='ID',all=T)
  T2R2Data<-merge(mvmt[,c(1,13)],T2R2Data,by='ID',all=T)
  }
  if(task=="SST"){T1Data<-merge(mvmt[,c(1,8)],T1Data,by='ID',all=T)
  T2Data<-merge(mvmt[,c(1,17)],T2Data,by='ID',all=T)
  T1R1Data<-merge(mvmt[,c(1,9)],T1R1Data,by='ID',all=T)            
  T2R1Data<-merge(mvmt[,c(1,18)],T2R1Data,by='ID',all=T)            
  T1R2Data<-merge(mvmt[,c(1,10)],T1R2Data,by='ID',all=T)
  T2R2Data<-merge(mvmt[,c(1,19)],T2R2Data,by='ID',all=T)
  }
  if(task=="nBack"){T1Data<-merge(mvmt[,c(1,5)],T1Data,by='ID',all=T)
  T2Data<-merge(mvmt[,c(1,14)],T2Data,by='ID',all=T)
  T1R1Data<-merge(mvmt[,c(1,6)],T1R1Data,by='ID',all=T)            
  T2R1Data<-merge(mvmt[,c(1,15)],T2R1Data,by='ID',all=T)            
  T1R2Data<-merge(mvmt[,c(1,7)],T1R2Data,by='ID',all=T)
  T2R2Data<-merge(mvmt[,c(1,16)],T2R2Data,by='ID',all=T)
  }
  toregressT1<-names(T1Data)[3:ncol(T1Data)]
  toregressT2<-names(T2Data)[3:ncol(T2Data)]
  toregressT1R1<-names(T1R1Data)[3:ncol(T1R1Data)]
  toregressT2R1<-names(T2R1Data)[3:ncol(T2R1Data)]
  toregressT1R2<-names(T1R2Data)[3:ncol(T1R2Data)]
  toregressT2R2<-names(T2R2Data)[3:ncol(T2R2Data)]
  
  timepoints<-c('T1Data','T2Data','T1R1Data','T1R2Data','T2R1Data','T2R2Data')
  
  for(z in timepoints){
    datatime<-get(z)
    datatime<-datatime[complete.cases(datatime[,2:3]),]
    IDVars<-as.data.frame(datatime$ID)
    regressed<-data.frame(matrix(ncol=0,nrow=nrow(datatime)))
    toregress<-names(datatime)[3:ncol(datatime)]
    for(x in toregress){ #This code was originally created to work around residualizing not handling missing data well, not really applicable to this dataset but was already written so...
      tmp<-(x)  
      tmp<-subset(datatime, select = tmp)
      if(all(is.na(tmp))==FALSE){
        tmp<-resid(lm(as.matrix(tmp) ~ datatime[,2], datatime, na.action = na.exclude))
        regressed<-cbind(regressed,tmp)}
      }
    regressed<-as.data.frame(regressed)
    names(regressed)<-toregress
    regressed<-cbind(IDVars,regressed)
    names(regressed)[1]<-'ID'
    assign(paste0(z,"_regressed"),regressed)
  }
  
  Avg<-merge(T1Data_regressed,T2Data_regressed,by='ID',all=T)
  
  R1R2<-merge(T1R1Data_regressed,T1R2Data_regressed,by='ID',all=T)
  R1R2<-merge(R1R2,T2R1Data_regressed,by='ID',all=T)
  R1R2<-merge(R1R2,T2R2Data_regressed,by='ID',all=T)
  
  tout<-Avg
  toout<-tout[,2:ncol(tout)]
  toout<-scale(toout)
  while(length(which(toout > abs(3)))!=0){
     toout[toout > 3]<-NA
     toout[toout < -3]<-NA
     toout<-scale(toout)
  }
  toout<-cbind(tout[,1],toout)
  tout[is.na(toout)]<-NA
  names(tout)[1]<-'ID'
  t1t2outname<-gsub('usable','Regressed',i)
  write.csv(tout,t1t2outname,row.names=F,na='')
  
  tout<-R1R2
  toout<-tout[,2:ncol(tout)]
  toout<-scale(toout)
  while(length(which(toout > abs(3)))!=0){
    toout[toout > 3]<-NA
    toout[toout < -3]<-NA
    toout<-scale(toout)
  }
  toout<-cbind(tout[,1],toout)
  tout[is.na(toout)]<-NA
  names(tout)[1]<-'ID'
  t1t2outname<-gsub('usable','Regressed',abcdR1R2name)
  write.csv(tout,t1t2outname,row.names=F,na='')
}
rm(list=ls())


#Create movement quartiles, regress out movement, and remove outliers
mvmt<-read.csv('mvmt.csv')
abcd<-list.files(pattern='^usable_renamed_.*._Avg_Beta_.*.csv')

for(i in abcd){
  abcddata<-read.csv(i)
  abcdR1R2name<-gsub('Avg','R1R2',i)
  abcdR1R2<-read.csv(abcdR1R2name)
  namecomp<-unlist(strsplit(i,"_"))
  if(namecomp[3]=='MID'){task<-"MID"}
  if(namecomp[3]=='nBack'){task<-"nBack"}
  if(namecomp[3]=='SST'){task<-"SST"}
  T1Data<-cbind(abcddata[,1],abcddata[grepl('_0m',names(abcddata))])
  names(T1Data)[1]<-'ID'
  T2Data<-cbind(abcddata[,1],abcddata[grepl('_24m',names(abcddata))])
  names(T2Data)[1]<-'ID'
  T1R1R2<-cbind(abcdR1R2[,1],abcdR1R2[grepl('_0m',names(abcdR1R2))])
  names(T1R1R2)[1]<-'ID'
  T2R1R2<-cbind(abcdR1R2[,1],abcdR1R2[grepl('_24m',names(abcdR1R2))])
  names(T2R1R2)[1]<-'ID'
  T1R1Data<-cbind(T1R1R2[,1],T1R1R2[grepl('_run1_',names(T1R1R2))])
  names(T1R1Data)[1]<-'ID'
  T1R2Data<-cbind(T1R1R2[,1],T1R1R2[grepl('_run2_',names(T1R1R2))])
  names(T1R2Data)[1]<-'ID'
  
  T2R1Data<-cbind(T2R1R2[,1],T2R1R2[grepl('_run1_',names(T2R1R2))])
  names(T2R1Data)[1]<-'ID'
  T2R2Data<-cbind(T2R1R2[,1],T2R1R2[grepl('_run2_',names(T2R1R2))])
  names(T2R2Data)[1]<-'ID'
  
  if(task=="MID"){T1Data<-merge(mvmt[,1:2],T1Data,by='ID',all=T)
  T2Data<-merge(mvmt[,c(1,11)],T2Data,by='ID',all=T)
  T1R1Data<-merge(mvmt[,c(1,3)],T1R1Data,by='ID',all=T)            
  T2R1Data<-merge(mvmt[,c(1,12)],T2R1Data,by='ID',all=T)            
  T1R2Data<-merge(mvmt[,c(1,4)],T1R2Data,by='ID',all=T)
  T2R2Data<-merge(mvmt[,c(1,13)],T2R2Data,by='ID',all=T)
  }
  if(task=="SST"){T1Data<-merge(mvmt[,c(1,8)],T1Data,by='ID',all=T)
  T2Data<-merge(mvmt[,c(1,17)],T2Data,by='ID',all=T)
  T1R1Data<-merge(mvmt[,c(1,9)],T1R1Data,by='ID',all=T)            
  T2R1Data<-merge(mvmt[,c(1,18)],T2R1Data,by='ID',all=T)            
  T1R2Data<-merge(mvmt[,c(1,10)],T1R2Data,by='ID',all=T)
  T2R2Data<-merge(mvmt[,c(1,19)],T2R2Data,by='ID',all=T)
  }
  if(task=="nBack"){T1Data<-merge(mvmt[,c(1,5)],T1Data,by='ID',all=T)
  T2Data<-merge(mvmt[,c(1,14)],T2Data,by='ID',all=T)
  T1R1Data<-merge(mvmt[,c(1,6)],T1R1Data,by='ID',all=T)            
  T2R1Data<-merge(mvmt[,c(1,15)],T2R1Data,by='ID',all=T)            
  T1R2Data<-merge(mvmt[,c(1,7)],T1R2Data,by='ID',all=T)
  T2R2Data<-merge(mvmt[,c(1,16)],T2R2Data,by='ID',all=T)
  }
  toregressT1<-names(T1Data)[3:ncol(T1Data)]
  toregressT2<-names(T2Data)[3:ncol(T2Data)]
  toregressT1R1<-names(T1R1Data)[3:ncol(T1R1Data)]
  toregressT2R1<-names(T2R1Data)[3:ncol(T2R1Data)]
  toregressT1R2<-names(T1R2Data)[3:ncol(T1R2Data)]
  toregressT2R2<-names(T2R2Data)[3:ncol(T2R2Data)]
  
  timepoints<-c('T1Data','T2Data','T1R1Data','T1R2Data','T2R1Data','T2R2Data')
  quartiles<-c('firstquartile','secondquartile','thirdquartile','fourthquartile')
  
  for(z in timepoints){
    
    datatime<-get(z)
    datatime<-datatime[complete.cases(datatime[,2:3]),]
    datatime<-datatime[order(datatime[,2]),]
    quartilelength<-round(.25*nrow(datatime))
    firstquartile<-datatime[1:quartilelength,]
    secondquartile<-datatime[(quartilelength+1):(2*quartilelength),]
    thirdquartile<-datatime[(2*quartilelength+1):(3*quartilelength),]
    fourthquartile<-datatime[(3*quartilelength+1):nrow(datatime),]
    for(y in quartiles){
      datatimequartile<-get(y)
      IDVars<-as.data.frame(datatimequartile$ID)
      regressed<-data.frame(matrix(ncol=0,nrow=nrow(datatimequartile)))
      toregress<-names(datatimequartile)[3:ncol(datatimequartile)]
      for(x in toregress){
        tmp<-(x)  
        tmp<-subset(datatimequartile, select = tmp)
        if(all(is.na(tmp))==FALSE){
          tmp<-resid(lm(as.matrix(tmp) ~ datatimequartile[,2], datatimequartile, na.action = na.exclude))
          regressed<-cbind(regressed,tmp)}
        assign(x, tmp)
      }
      regressed<-as.data.frame(regressed)
      names(regressed)<-toregress
      regressed<-cbind(IDVars,regressed)
      names(regressed)[1]<-'ID'
      assign(paste0(z,"_",y),regressed)
    }
    
  }
  
  
  firstquartileAvg<-merge(T1Data_firstquartile,T2Data_firstquartile,by='ID',all=T)
  secondquartileAvg<-merge(T1Data_secondquartile,T2Data_secondquartile,by='ID',all=T)
  thirdquartileAvg<-merge(T1Data_thirdquartile,T2Data_thirdquartile,by='ID',all=T)
  fourthquartileAvg<-merge(T1Data_fourthquartile,T2Data_fourthquartile,by='ID',all=T)
  
  firstquartileR1R2<-merge(T1R1Data_firstquartile,T1R2Data_firstquartile,by='ID',all=T)
  firstquartileR1R2<-merge(firstquartileR1R2,T2R1Data_firstquartile,by='ID',all=T)
  firstquartileR1R2<-merge(firstquartileR1R2,T2R2Data_firstquartile,by='ID',all=T)
  
  secondquartileR1R2<-merge(T1R1Data_secondquartile,T1R2Data_secondquartile,by='ID',all=T)
  secondquartileR1R2<-merge(secondquartileR1R2,T2R1Data_secondquartile,by='ID',all=T)
  secondquartileR1R2<-merge(secondquartileR1R2,T2R2Data_secondquartile,by='ID',all=T)  
  
  thirdquartileR1R2<-merge(T1R1Data_thirdquartile,T1R2Data_thirdquartile,by='ID',all=T)
  thirdquartileR1R2<-merge(thirdquartileR1R2,T2R1Data_thirdquartile,by='ID',all=T)
  thirdquartileR1R2<-merge(thirdquartileR1R2,T2R2Data_thirdquartile,by='ID',all=T)
  
  fourthquartileR1R2<-merge(T1R1Data_fourthquartile,T1R2Data_fourthquartile,by='ID',all=T)
  fourthquartileR1R2<-merge(fourthquartileR1R2,T2R1Data_fourthquartile,by='ID',all=T)
  fourthquartileR1R2<-merge(fourthquartileR1R2,T2R2Data_fourthquartile,by='ID',all=T)
  
  outliers<-c('firstquartileAvg',"firstquartileR1R2",'secondquartileAvg',"secondquartileR1R2",'thirdquartileAvg',"thirdquartileR1R2",'fourthquartileAvg',"fourthquartileR1R2")
  
  for(w in outliers){
    tout<-get(w)
    toout<-tout[,2:ncol(tout)]
    toout<-scale(toout)
    while(length(which(toout > abs(3)))!=0){
      toout[toout > 3]<-NA
      toout[toout < -3]<-NA
      toout<-scale(toout)
    }
    toout<-cbind(tout[,1],toout)
    tout[is.na(toout)]<-NA
    t1t2outname<-paste0(w,"_",gsub('renamed','regressed',i))
    t1t2outname<-gsub('quartileAvg','quartile',t1t2outname)
    t1t2r1r2outname<-paste0(w,"_",gsub('renamed','regressed',abcdR1R2name))
    t1t2r1r2outname<-gsub('quartileR1R2','quartile',t1t2r1r2outname)
    if(grepl("Avg$",w)==TRUE){
      write.csv(tout,t1t2outname,row.names=F,na='')
    }else{
      write.csv(tout,t1t2r1r2outname,row.names=F,na='')
    }
  }
}
rm(list=ls())
```
