## Supplementary material for "Reliability and Stability Challenges in ABCD Task fMRI Data": Code for generating data and results: Script 2 - ReliabilityCalcCI.R

```
#ICC, activity, and change calculations.
#Set the working directory to the previously created ICC folder (line 10)
#Choose the sample you're interested in in the list.files pattern (line 11, currently searching for 'usable'), edit the output name (line 12),
#Select all and run and it will calculate values for all variables

if('irr' %in% rownames(installed.packages())==FALSE){
  install.packages('irr')}
library('irr')
library('psych')
setwd("C:/Users/James/Desktop/Release 3 Nov 2020/ICC")
measures<-list.files(pattern='^Rank_.*.R1R2.*.csv$')
outname<-'ICCs_Rank.csv'
combOut<-matrix(ncol=109,nrow=0)
for(i in measures){
  data<-read.csv(i, stringsAsFactors = F)
  dataavg<-gsub('R1R2','Avg',i)
  dataavg<-read.csv(dataavg)
  measureOut<-paste0('ICC(3,2)_Comparison',i)
  datanames<-names(data)
  datanamesstripped<-gsub("_run1_",'RunSame',datanames)
  datanamesstripped<-gsub("_run2_",'RunSame',datanamesstripped)
  datanamesstripped<-gsub("_0m",'',datanamesstripped)
  datanamesstripped<-gsub("_24m",'',datanamesstripped)
  usablevariables<-datanamesstripped[duplicated(datanamesstripped)]
  usablevariables<-unique(usablevariables)
  names(data)[1]<-'ID'
  names(dataavg)[1]<-'ID'
  data<-merge(data,dataavg,by='ID',all=T)
  output<-matrix(ncol=108,nrow=0)
  for(z in usablevariables){
    varnameR1T1<-paste0(gsub('RunSame',"_run1_",z),"_0m")
    varnameR1T2<-paste0(gsub('RunSame',"_run1_",z),"_24m")
    varnameR2T1<-paste0(gsub('RunSame',"_run2_",z),"_0m")
    varnameR2T2<-paste0(gsub('RunSame',"_run2_",z),"_24m")    
    varnameT1<-paste0(gsub('RunSame',"_all_",z),"_0m")
    varnameT2<-paste0(gsub('RunSame',"_all_",z),"_24m")        
    datause<-cbind(data$ID,data[[varnameR1T1]],data[[varnameR1T2]],data[[varnameR2T1]],data[[varnameR2T2]],data[[varnameT1]],data[[varnameT2]])
    datause<-as.data.frame(datause)
    names(datause)<-c('ID','R1T1','R1T2','R2T1','R2T2','T1','T2')
    datause$R1T1<-as.numeric(datause$R1T1)
    datause$R1T2<-as.numeric(datause$R1T2)
    datause$R2T1<-as.numeric(datause$R2T1)
    datause$R2T2<-as.numeric(datause$R2T2)
    datause$T1<-as.numeric(datause$T1)
    datause$T2<-as.numeric(datause$T2)
    iccR1output<-icc(cbind(datause$R1T1,datause$R1T2), model = 'twoway', type = 'consistency', unit='single')
    iccR2output<-icc(cbind(datause$R2T1,datause$R2T2), model = 'twoway', type = 'consistency', unit='single')
    iccT1output<-icc(cbind(datause$R1T1,datause$R2T1), model = 'twoway', type = 'consistency', unit='average')
    iccT2output<-icc(cbind(datause$R1T2,datause$R2T2), model = 'twoway', type = 'consistency', unit='average')
    iccoutput<-icc(cbind(datause$T1,datause$T2), model = 'twoway', type = 'consistency', 'single')
    oneSTT1<-t.test(datause$T1,mu=0)
    oneSTT2<-t.test(datause$T2,mu=0)
    oneSDT1<-(mean(datause$T1,na.rm=T)/(sd(datause$T1,na.rm=T)))
    oneSDT1CI<-cohen.d.ci(d=oneSDT1,n=length(complete.cases(datause$T1)))
    oneSDT2<-(mean(datause$T2,na.rm=T)/(sd(datause$T2,na.rm=T)))
    oneSDT2CI<-cohen.d.ci(d=oneSDT2,n=length(complete.cases(datause$T2)))
    oneSTR1T1<-t.test(datause$R1T1,mu=0)
    oneSTR1T2<-t.test(datause$R1T2,mu=0)
    oneSDR1T1<-(mean(datause$R1T1,na.rm=T)/(sd(datause$R1T1,na.rm=T)))
    oneSDR1T1CI<-cohen.d.ci(d=oneSDR1T1,n=length(complete.cases(datause$R1T1)))
    oneSDR1T2<-(mean(datause$R1T2,na.rm=T)/(sd(datause$R1T2,na.rm=T)))
    oneSDR1T2CI<-cohen.d.ci(d=oneSDR1T2,n=length(complete.cases(datause$R1T2)))
    oneSTR2T1<-t.test(datause$R2T1,mu=0)
    oneSTR2T2<-t.test(datause$R2T2,mu=0)
    oneSDR2T1<-(mean(datause$R2T1,na.rm=T)/(sd(datause$R2T1,na.rm=T)))
    oneSDR2T1CI<-cohen.d.ci(d=oneSDR2T1,n=length(complete.cases(datause$R2T1)))
    oneSDR2T2<-(mean(datause$R2T2,na.rm=T)/(sd(datause$R2T2,na.rm=T)))
    oneSDR2T2CI<-cohen.d.ci(d=oneSDR2T2,n=length(complete.cases(datause$R2T2)))
    pairedtBetween<-t.test(datause$T2,datause$T1,paired=T)
    pairedtDBetween<-((mean(datause$T2-datause$T1,na.rm = T))/sd(datause$T2-datause$T1,na.rm = T))
    pairedtDBetweenCI<-cohen.d.ci(d=pairedtDBetween,n1=length(complete.cases(datause$T1)),n2=length(complete.cases(datause$T2)))
    pairedtWithinT1<-t.test(datause$R2T1,datause$R1T1,paired=T)
    pairedtDWithinT1<-((mean(datause$R2T1-datause$R1T1,na.rm = T))/sd(datause$R2T1-datause$R1T1,na.rm = T))
    pairedtDWithinT1CI<-cohen.d.ci(d=pairedtDWithinT1,n1=length(complete.cases(datause$R1T1)),n2=length(complete.cases(datause$R2T1)))
    pairedtWithinT2<-t.test(datause$R2T2,datause$R1T2,paired=T)
    pairedtDWithinT2<-((mean(datause$R2T2-datause$R1T2,na.rm = T))/sd(datause$R2T2-datause$R1T2,na.rm = T))
    pairedtDWithinT2CI<-cohen.d.ci(d=pairedtDWithinT2,n1=length(complete.cases(datause$R1T2)),n2=length(complete.cases(datause$R2T2)))
    pairedtBetweenR1<-t.test(datause$R1T2,datause$R1T1,paired=T)
    pairedtDBetweenR1<-((mean(datause$R1T2-datause$R1T1,na.rm = T))/sd(datause$R1T2-datause$R1T1,na.rm = T))
    pairedtDBetweenR1CI<-cohen.d.ci(d=pairedtDBetweenR1,n1=length(complete.cases(datause$R1T1)),n2=length(complete.cases(datause$R1T2)))
    pairedtBetweenR2<-t.test(datause$R2T2,datause$R2T1,paired=T)
    pairedtDBetweenR2<-((mean(datause$R2T2-datause$R2T1,na.rm = T))/sd(datause$R2T2-datause$R2T1,na.rm = T))
    pairedtDBetweenR2CI<-cohen.d.ci(d=pairedtDBetweenR2,n1=length(complete.cases(datause$R2T1)),n2=length(complete.cases(datause$R2T2)))
    
    betweendata<-datause[,c(1,6,7)]
    betweendata<-betweendata[complete.cases(betweendata),]
    betweenSStotal<-(var(unlist(betweendata[,2:3]))*((nrow(betweendata)*(ncol(betweendata)-1))-1))
    betweenMSR<-var(rowMeans(betweendata[,2:3]))*(ncol(betweendata)-1)
    betweenMSC<-var(colMeans(betweendata[,2:3]))*nrow(betweendata)
    betweenMSE<-((betweenSStotal-betweenMSR*(nrow(betweendata)-1)-betweenMSC*(ncol(betweendata)-2)))/((nrow(betweendata)-1)*(ncol(betweendata)-2))
    betweenR1data<-datause[,c(1,2,3)]
    betweenR1data<-betweenR1data[complete.cases(betweenR1data),]
    betweenR1SStotal<-(var(unlist(betweenR1data[,2:3]))*((nrow(betweenR1data)*(ncol(betweenR1data)-1))-1))
    betweenR1MSR<-var(rowMeans(betweenR1data[,2:3]))*(ncol(betweenR1data)-1)
    betweenR1MSC<-var(colMeans(betweenR1data[,2:3]))*nrow(betweenR1data)
    betweenR1MSE<-((betweenR1SStotal-betweenR1MSR*(nrow(betweenR1data)-1)-betweenR1MSC*(ncol(betweenR1data)-2)))/((nrow(betweenR1data)-1)*(ncol(betweenR1data)-2))
    betweenR2data<-datause[,c(1,4,5)]
    betweenR2data<-betweenR2data[complete.cases(betweenR2data),]
    betweenR2SStotal<-(var(unlist(betweenR2data[,2:3]))*((nrow(betweenR2data)*(ncol(betweenR2data)-1))-1))
    betweenR2MSR<-var(rowMeans(betweenR2data[,2:3]))*(ncol(betweenR2data)-1)
    betweenR2MSC<-var(colMeans(betweenR2data[,2:3]))*nrow(betweenR2data)
    betweenR2MSE<-((betweenR2SStotal-betweenR2MSR*(nrow(betweenR2data)-1)-betweenR2MSC*(ncol(betweenR2data)-2)))/((nrow(betweenR2data)-1)*(ncol(betweenR2data)-2))
    withinT1data<-datause[,c(1,2,4)]
    withinT1data<-withinT1data[complete.cases(withinT1data),]
    withinT1SStotal<-(var(unlist(withinT1data[,2:3]))*((nrow(withinT1data)*(ncol(withinT1data)-1))-1))
    withinT1MSR<-var(rowMeans(withinT1data[,2:3]))*(ncol(withinT1data)-1)
    withinT1MSC<-var(colMeans(withinT1data[,2:3]))*nrow(withinT1data)
    withinT1MSE<-((withinT1SStotal-withinT1MSR*(nrow(withinT1data)-1)-withinT1MSC*(ncol(withinT1data)-2)))/((nrow(withinT1data)-1)*(ncol(withinT1data)-2))
    withinT2data<-datause[,c(1,3,5)]
    withinT2data<-withinT2data[complete.cases(withinT2data),]
    withinT2SStotal<-(var(unlist(withinT2data[,2:3]))*((nrow(withinT2data)*(ncol(withinT2data)-1))-1))
    withinT2MSR<-var(rowMeans(withinT2data[,2:3]))*(ncol(withinT2data)-1)
    withinT2MSC<-var(colMeans(withinT2data[,2:3]))*nrow(withinT2data)
    withinT2MSE<-((withinT2SStotal-withinT2MSR*(nrow(withinT2data)-1)-withinT2MSC*(ncol(withinT2data)-2)))/((nrow(withinT2data)-1)*(ncol(withinT2data)-2))
    
    results<-cbind(z,iccoutput$value,iccoutput$lbound,iccoutput$ubound,iccT1output$value,iccT1output$lbound,iccT1output$ubound,iccT2output$value,iccT2output$lbound,iccT2output$ubound,
                   iccR1output$value,iccR1output$lbound,iccR1output$ubound,iccR2output$value,iccR2output$lbound,iccR2output$ubound,
                   oneSTT1$statistic,oneSTT1$p.value,oneSTT1$conf.int[1],oneSTT1$conf.int[2],oneSTT2$statistic,oneSTT2$p.value,oneSTT2$conf.int[1],oneSTT2$conf.int[2],
                   oneSTR1T1$statistic,oneSTR1T1$p.value,oneSTR1T1$conf.int[1],oneSTR1T1$conf.int[2],
                   oneSTR2T1$statistic,oneSTR2T1$p.value,oneSTR2T1$conf.int[1],oneSTR2T1$conf.int[2],
                   oneSTR1T2$statistic,oneSTR1T2$p.value,oneSTR1T2$conf.int[1],oneSTR1T2$conf.int[2],
                   oneSTR2T2$statistic,oneSTR2T2$p.value,oneSTR2T2$conf.int[1],oneSTR2T2$conf.int[2],
                   oneSDT1,oneSDT1CI[1,1],oneSDT1CI[1,3],oneSDT2,oneSDT2CI[1,1],oneSDT2CI[1,3],
                   oneSDR1T1,oneSDR1T1CI[1,1],oneSDR1T1CI[1,3],oneSDR2T1,oneSDR2T1CI[1,1],oneSDR2T1CI[1,3],
                   oneSDR1T2,oneSDR1T2CI[1,1],oneSDR1T2CI[1,3],oneSDR2T2,oneSDR2T2CI[1,1],oneSDR2T2CI[1,3],
                   pairedtBetween$statistic,pairedtBetween$p.value,pairedtBetween$conf.int[1],pairedtBetween$conf.int[2],
                   pairedtWithinT1$statistic,pairedtWithinT1$p.value,pairedtWithinT1$conf.int[1],pairedtWithinT1$conf.int[2],
                   pairedtWithinT2$statistic,pairedtWithinT2$p.value,pairedtWithinT2$conf.int[1],pairedtWithinT2$conf.int[2],
                   pairedtBetweenR1$statistic,pairedtBetweenR1$p.value,pairedtBetweenR1$conf.int[1],pairedtBetweenR1$conf.int[2],
                   pairedtBetweenR2$statistic,pairedtBetweenR2$p.value,pairedtBetweenR2$conf.int[1],pairedtBetweenR2$conf.int[2],
                   pairedtDBetween,pairedtDBetweenCI[1,1],pairedtDBetweenCI[1,3],
                   pairedtDWithinT1,pairedtDWithinT1CI[1,1],pairedtDWithinT1CI[1,3],
                   pairedtDWithinT2,pairedtDWithinT2CI[1,1],pairedtDWithinT2CI[1,3],
                   pairedtDBetweenR1,pairedtDBetweenR1CI[1,1],pairedtDBetweenR1CI[1,3],
                   pairedtDBetweenR2,pairedtDBetweenR2CI[1,1],pairedtDBetweenR2CI[1,3],
                   betweenMSR,betweenMSC,betweenMSE,betweenR1MSR,betweenR1MSC,betweenR1MSE,betweenR2MSR,betweenR2MSC,betweenR2MSE,
                   withinT1MSR,withinT1MSC,withinT1MSE,withinT2MSR,withinT2MSC,withinT2MSE)
    output<-rbind(output,results)
  }
  combOut<-rbind(combOut,cbind(i,output))
  output<-as.data.frame(output)
  names(output)<-c('Variable','ICC Between','ICC Between 95% CI Lower','ICC Between 95% CI Upper',
                   'ICC Within T1','ICC Within T1 95% CI Lower','ICC Within T1 95% CI Upper',
                   'ICC Within T2','ICC Within T2 95% CI Lower','ICC Within T2 95% CI Upper',
                   'ICC Between R1','ICC Between R1 95% CI Lower','ICC Between R1 95% CI Upper',
                   'ICC Between R2','ICC Between R2 95% CI Lower','ICC Between R2 95% CI Upper',
                   'OneSTT1 t','OneSTT1 p','OneSTT1 95% CI Lower Mean','OneSTT1 95% CI Upper Mean',
                   'OneSTT2 t','OneSTT2 p','OneSTT2 95% CI Lower Mean','OneSTT2 95% CI Upper Mean',
                   'OneSTR1T1 t','OneSTR1T1 p','OneSTR1T1 95% CI Lower Mean','OneSTR1T1 95% CI Upper Mean',
                   'OneSTR2T1 t','OneSTR2T1 p','OneSTR2T1 95% CI Lower Mean','OneSTR2T1 95% CI Upper Mean',
                   'OneSTR1T2 t','OneSTR1T2 p','OneSTR1T2 95% CI Lower Mean','OneSTR1T2 95% CI Upper Mean',
                   'OneSTR2T2 t','OneSTR2T2 p','OneSTR2T2 95% CI Lower Mean','OneSTR2T2 95% CI Upper Mean',
                   'OneSTT1 D','OneSTT1 D 95% CI Lower','OneSTT1 D 95% CI Upper',
                   'OneSTT2 D','OneSTT2 D 95% CI Lower','OneSTT2 D 95% CI Upper',
                   'OneSTR1T1 D','OneSTR1T1 D 95% CI Lower','OneSTR1T1 D 95% CI Upper',
                   'OneSTR2T1 D','OneSTR2T1 D 95% CI Lower','OneSTR2T1 D 95% CI Upper',
                   'OneSTR1T2 D','OneSTR1T2 D 95% CI Lower','OneSTR1T2 D 95% CI Upper',
                   'OneSTR2T2 D','OneSTR2T2 D 95% CI Lower','OneSTR2T2 D 95% CI Upper',
                   'Paired T Between t','Paired T Between p','Paired T Between 95% CI Lower Difference','Paired T Between 95% CI Upper Difference',
                   'Paired T Within T1 t','Paired T Within T1 p','Paired T Within T1 95% CI Lower Difference','Paired T Within T1 95% CI Upper Difference',
                   'Paired T Within T2 t','Paired T Within T2 p','Paired T Within T2 95% CI Lower Difference','Paired T Within T2 95% CI Upper Difference',
                   'Paired T Between R1 t','Paired T Between R1 p','Paired T Between R1 95% CI Lower Difference','Paired T Between R1 95% CI Upper Difference',
                   'Paired T Between R2 t','Paired T Between R2 p','Paired T Between R2 95% CI Lower Difference','Paired T Between R2 95% CI Upper Difference',
                   'Paired T Between D','Paired T Between D 95% CI Lower','Paired T Between D 95% CI Upper',
                   'Paired T Within T1 D','Paired T Within T1 D 95% CI Lower','Paired T Within T1 D 95% CI Upper',
                   'Paired T Within T2 D','Paired T Within T2 D 95% CI Lower','Paired T Within T2 D 95% CI Upper',
                   'Paired T Between R1 D','Paired T Between R1 D 95% CI Lower','Paired T Between R1 D 95% CI Upper',
                   'Paired T Between R2 D','Paired T Between R2 D 95% CI Lower','Paired T Between R2 D 95% CI Upper',
                   'Between MSR','Between MSC','Between MSE','Between R1 MSR','Between R1 MSC','Between R1 MSE','Between R2 MSR','Between R2 MSC','Between R2 MSE',
                   'Within T1 MSR','Within T1 MSC','Within T1 MSE','Within T2 MSR','Within T2 MSC','Within T2 MSE')
  write.csv(output,measureOut,row.names = FALSE)
}
combOut<-as.data.frame(combOut)
names(combOut)<-c('File','Variable','ICC Between','ICC Between 95% CI Lower','ICC Between 95% CI Upper',
                  'ICC Within T1','ICC Within T1 95% CI Lower','ICC Within T1 95% CI Upper',
                  'ICC Within T2','ICC Within T2 95% CI Lower','ICC Within T2 95% CI Upper',
                  'ICC Between R1','ICC Between R1 95% CI Lower','ICC Between R1 95% CI Upper',
                  'ICC Between R2','ICC Between R2 95% CI Lower','ICC Between R2 95% CI Upper',
                  'OneSTT1 t','OneSTT1 p','OneSTT1 95% CI Lower Mean','OneSTT1 95% CI Upper Mean',
                  'OneSTT2 t','OneSTT2 p','OneSTT2 95% CI Lower Mean','OneSTT2 95% CI Upper Mean',
                  'OneSTR1T1 t','OneSTR1T1 p','OneSTR1T1 95% CI Lower Mean','OneSTR1T1 95% CI Upper Mean',
                  'OneSTR2T1 t','OneSTR2T1 p','OneSTR2T1 95% CI Lower Mean','OneSTR2T1 95% CI Upper Mean',
                  'OneSTR1T2 t','OneSTR1T2 p','OneSTR1T2 95% CI Lower Mean','OneSTR1T2 95% CI Upper Mean',
                  'OneSTR2T2 t','OneSTR2T2 p','OneSTR2T2 95% CI Lower Mean','OneSTR2T2 95% CI Upper Mean',
                  'OneSTT1 D','OneSTT1 D 95% CI Lower','OneSTT1 D 95% CI Upper',
                  'OneSTT2 D','OneSTT2 D 95% CI Lower','OneSTT2 D 95% CI Upper',
                  'OneSTR1T1 D','OneSTR1T1 D 95% CI Lower','OneSTR1T1 D 95% CI Upper',
                  'OneSTR2T1 D','OneSTR2T1 D 95% CI Lower','OneSTR2T1 D 95% CI Upper',
                  'OneSTR1T2 D','OneSTR1T2 D 95% CI Lower','OneSTR1T2 D 95% CI Upper',
                  'OneSTR2T2 D','OneSTR2T2 D 95% CI Lower','OneSTR2T2 D 95% CI Upper',
                  'Paired T Between t','Paired T Between p','Paired T Between 95% CI Lower Difference','Paired T Between 95% CI Upper Difference',
                  'Paired T Within T1 t','Paired T Within T1 p','Paired T Within T1 95% CI Lower Difference','Paired T Within T1 95% CI Upper Difference',
                  'Paired T Within T2 t','Paired T Within T2 p','Paired T Within T2 95% CI Lower Difference','Paired T Within T2 95% CI Upper Difference',
                  'Paired T Between R1 t','Paired T Between R1 p','Paired T Between R1 95% CI Lower Difference','Paired T Between R1 95% CI Upper Difference',
                  'Paired T Between R2 t','Paired T Between R2 p','Paired T Between R2 95% CI Lower Difference','Paired T Between R2 95% CI Upper Difference',
                  'Paired T Between D','Paired T Between D 95% CI Lower','Paired T Between D 95% CI Upper',
                  'Paired T Within T1 D','Paired T Within T1 D 95% CI Lower','Paired T Within T1 D 95% CI Upper',
                  'Paired T Within T2 D','Paired T Within T2 D 95% CI Lower','Paired T Within T2 D 95% CI Upper',
                  'Paired T Between R1 D','Paired T Between R1 D 95% CI Lower','Paired T Between R1 D 95% CI Upper',
                  'Paired T Between R2 D','Paired T Between R2 D 95% CI Lower','Paired T Between R2 D 95% CI Upper',
                  'Between MSR','Between MSC','Between MSE','Between R1 MSR','Between R1 MSC','Between R1 MSE','Between R2 MSR','Between R2 MSC','Between R2 MSE',
                  'Within T1 MSR','Within T1 MSC','Within T1 MSE','Within T2 MSR','Within T2 MSC','Within T2 MSE')
write.csv(combOut,outname,row.names = FALSE)

combined<-combOut

combined$Contrast<-combined$Variable
combined$Contrast<-gsub('_beta_.*.','',combined$Contrast)
combined$Contrast<-gsub('.*._all_','',combined$Contrast)
combined$Contrast<-gsub('.*.RunSame','',combined$Contrast)

combined$Parcel<-combined$Variable
combined$Parcel<-gsub('.*._aseg_','',combined$Parcel)
combined$Parcel<-gsub('.*._destrieux_','',combined$Parcel)
combined$Parcel<-gsub('.*._desikan_','',combined$Parcel)
combined$Parcel<-gsub('RunSame','',combined$Parcel)

combined<-combined[!(combined$Parcel=='cerebral_white_matter_lh'),]
combined<-combined[!(combined$Parcel=='cerebral_white_matter_rh'),]
combined<-combined[!(combined$Parcel=='cerebellum_white_matter_lh'),]
combined<-combined[!(combined$Parcel=='cerebellum_white_matter_rh'),]
combined<-combined[!(combined$Parcel=='lateral_ventricle_lh'),]
combined<-combined[!(combined$Parcel=='lateral_ventricle_rh'),]
combined<-combined[!(combined$Parcel=='inf_lat_vent_lh'),]
combined<-combined[!(combined$Parcel=='inf_lat_vent_rh'),]
combined<-combined[!(combined$Parcel=='csf'),]
combined<-combined[!(combined$Parcel=='3rd_ventricle'),]
combined<-combined[!(combined$Parcel=='4th_ventricle'),]

combined<-combined[!(combined$Contrast==''),]

combined$order<-combined$Parcel
combined$order<-gsub('g_and_s_frontomargin_lh','Dx001',combined$order)
combined$order<-gsub('g_and_s_occipital_inf_lh','Dx002',combined$order)
combined$order<-gsub('g_and_s_paracentral_lh','Dx003',combined$order)
combined$order<-gsub('g_and_s_subcentral_lh','Dx004',combined$order)
combined$order<-gsub('g_and_s_transv_frontopol_lh','Dx005',combined$order)
combined$order<-gsub('g_and_s_cingul_ant_lh','Dx006',combined$order)
combined$order<-gsub('g_and_s_cingul_mid_ant_lh','Dx007',combined$order)
combined$order<-gsub('g_and_s_cingul_mid_post_lh','Dx008',combined$order)
combined$order<-gsub('g_cingul_post_dorsal_lh','Dx009',combined$order)
combined$order<-gsub('g_cingul_post_ventral_lh','Dx010',combined$order)
combined$order<-gsub('g_cuneus_lh','Dx011',combined$order)
combined$order<-gsub('g_front_inf_opercular_lh','Dx012',combined$order)
combined$order<-gsub('g_front_inf_orbital_lh','Dx013',combined$order)
combined$order<-gsub('g_front_inf_triangul_lh','Dx014',combined$order)
combined$order<-gsub('g_front_middle_lh','Dx015',combined$order)
combined$order<-gsub('g_front_sup_lh','Dx016',combined$order)
combined$order<-gsub('g_ins_lg_and_s_cent_ins_lh','Dx017',combined$order)
combined$order<-gsub('g_insular_short_lh','Dx018',combined$order)
combined$order<-gsub('g_occipital_middle_lh','Dx019',combined$order)
combined$order<-gsub('g_occipital_sup_lh','Dx020',combined$order)
combined$order<-gsub('g_oc_temp_lat_fusifor_lh','Dx021',combined$order)
combined$order<-gsub('g_oc_temp_med_lingual_lh','Dx022',combined$order)
combined$order<-gsub('g_oc_temp_med_parahip_lh','Dx023',combined$order)
combined$order<-gsub('g_orbital_lh','Dx024',combined$order)
combined$order<-gsub('g_pariet_inf_angular_lh','Dx025',combined$order)
combined$order<-gsub('g_pariet_inf_supramar_lh','Dx026',combined$order)
combined$order<-gsub('g_parietal_sup_lh','Dx027',combined$order)
combined$order<-gsub('g_postcentral_lh','Dx028',combined$order)
combined$order<-gsub('g_precentral_lh','Dx029',combined$order)
combined$order<-gsub('g_precuneus_lh','Dx030',combined$order)
combined$order<-gsub('g_rectus_lh','Dx031',combined$order)
combined$order<-gsub('g_subcallosal_lh','Dx032',combined$order)
combined$order<-gsub('g_temp_sup_g_t_transv_lh','Dx033',combined$order)
combined$order<-gsub('g_temp_sup_lateral_lh','Dx034',combined$order)
combined$order<-gsub('g_temp_sup_plan_polar_lh','Dx035',combined$order)
combined$order<-gsub('g_temp_sup_plan_tempo_lh','Dx036',combined$order)
combined$order<-gsub('g_temporal_inf_lh','Dx037',combined$order)
combined$order<-gsub('g_temporal_middle_lh','Dx038',combined$order)
combined$order<-gsub('lat_fis_ant_horizont_lh','Dx039',combined$order)
combined$order<-gsub('lat_fis_ant_vertical_lh','Dx040',combined$order)
combined$order<-gsub('lat_fis_post_lh','Dx041',combined$order)
combined$order<-gsub('pole_occipital_lh','Dx042',combined$order)
combined$order<-gsub('pole_temporal_lh','Dx043',combined$order)
combined$order<-gsub('s_calcarine_lh','Dx044',combined$order)
combined$order<-gsub('s_central_lh','Dx045',combined$order)
combined$order<-gsub('s_cingul_marginalis_lh','Dx046',combined$order)
combined$order<-gsub('s_circular_insula_ant_lh','Dx047',combined$order)
combined$order<-gsub('s_circular_insula_inf_lh','Dx048',combined$order)
combined$order<-gsub('s_circular_insula_sup_lh','Dx049',combined$order)
combined$order<-gsub('s_collat_transv_ant_lh','Dx050',combined$order)
combined$order<-gsub('s_collat_transv_post_lh','Dx051',combined$order)
combined$order<-gsub('s_front_inf_lh','Dx052',combined$order)
combined$order<-gsub('s_front_middle_lh','Dx053',combined$order)
combined$order<-gsub('s_front_sup_lh','Dx054',combined$order)
combined$order<-gsub('s_interm_prim_jensen_lh','Dx055',combined$order)
combined$order<-gsub('s_intrapariet_and_p_trans_lh','Dx056',combined$order)
combined$order<-gsub('s_oc_middle_and_lunatus_lh','Dx057',combined$order)
combined$order<-gsub('s_oc_sup_and_transversal_lh','Dx058',combined$order)
combined$order<-gsub('s_occipital_ant_lh','Dx059',combined$order)
combined$order<-gsub('s_oc_temp_lat_lh','Dx060',combined$order)
combined$order<-gsub('s_oc_temp_med_and_lingual_lh','Dx061',combined$order)
combined$order<-gsub('s_orbital_lateral_lh','Dx062',combined$order)
combined$order<-gsub('s_orbital_med_olfact_lh','Dx063',combined$order)
combined$order<-gsub('s_orbital_h_shaped_lh','Dx064',combined$order)
combined$order<-gsub('s_parieto_occipital_lh','Dx065',combined$order)
combined$order<-gsub('s_pericallosal_lh','Dx066',combined$order)
combined$order<-gsub('s_postcentral_lh','Dx067',combined$order)
combined$order<-gsub('s_precentral_inf_part_lh','Dx068',combined$order)
combined$order<-gsub('s_precentral_sup_part_lh','Dx069',combined$order)
combined$order<-gsub('s_suborbital_lh','Dx070',combined$order)
combined$order<-gsub('s_subparietal_lh','Dx071',combined$order)
combined$order<-gsub('s_temporal_inf_lh','Dx072',combined$order)
combined$order<-gsub('s_temporal_sup_lh','Dx073',combined$order)
combined$order<-gsub('s_temporal_transverse_lh','Dx074',combined$order)
combined$order<-gsub('g_and_s_frontomargin_rh','Dx075',combined$order)
combined$order<-gsub('g_and_s_occipital_inf_rh','Dx076',combined$order)
combined$order<-gsub('g_and_s_paracentral_rh','Dx077',combined$order)
combined$order<-gsub('g_and_s_subcentral_rh','Dx078',combined$order)
combined$order<-gsub('g_and_s_transv_frontopol_rh','Dx079',combined$order)
combined$order<-gsub('g_and_s_cingul_ant_rh','Dx080',combined$order)
combined$order<-gsub('g_and_s_cingul_mid_ant_rh','Dx081',combined$order)
combined$order<-gsub('g_and_s_cingul_mid_post_rh','Dx082',combined$order)
combined$order<-gsub('g_cingul_post_dorsal_rh','Dx083',combined$order)
combined$order<-gsub('g_cingul_post_ventral_rh','Dx084',combined$order)
combined$order<-gsub('g_cuneus_rh','Dx085',combined$order)
combined$order<-gsub('g_front_inf_opercular_rh','Dx086',combined$order)
combined$order<-gsub('g_front_inf_orbital_rh','Dx087',combined$order)
combined$order<-gsub('g_front_inf_triangul_rh','Dx088',combined$order)
combined$order<-gsub('g_front_middle_rh','Dx089',combined$order)
combined$order<-gsub('g_front_sup_rh','Dx090',combined$order)
combined$order<-gsub('g_ins_lg_and_s_cent_ins_rh','Dx091',combined$order)
combined$order<-gsub('g_insular_short_rh','Dx092',combined$order)
combined$order<-gsub('g_occipital_middle_rh','Dx093',combined$order)
combined$order<-gsub('g_occipital_sup_rh','Dx094',combined$order)
combined$order<-gsub('g_oc_temp_lat_fusifor_rh','Dx095',combined$order)
combined$order<-gsub('g_oc_temp_med_lingual_rh','Dx096',combined$order)
combined$order<-gsub('g_oc_temp_med_parahip_rh','Dx097',combined$order)
combined$order<-gsub('g_orbital_rh','Dx098',combined$order)
combined$order<-gsub('g_pariet_inf_angular_rh','Dx099',combined$order)
combined$order<-gsub('g_pariet_inf_supramar_rh','Dx100',combined$order)
combined$order<-gsub('g_parietal_sup_rh','Dx101',combined$order)
combined$order<-gsub('g_postcentral_rh','Dx102',combined$order)
combined$order<-gsub('g_precentral_rh','Dx103',combined$order)
combined$order<-gsub('g_precuneus_rh','Dx104',combined$order)
combined$order<-gsub('g_rectus_rh','Dx105',combined$order)
combined$order<-gsub('g_subcallosal_rh','Dx106',combined$order)
combined$order<-gsub('g_temp_sup_g_t_transv_rh','Dx107',combined$order)
combined$order<-gsub('g_temp_sup_lateral_rh','Dx108',combined$order)
combined$order<-gsub('g_temp_sup_plan_polar_rh','Dx109',combined$order)
combined$order<-gsub('g_temp_sup_plan_tempo_rh','Dx110',combined$order)
combined$order<-gsub('g_temporal_inf_rh','Dx111',combined$order)
combined$order<-gsub('g_temporal_middle_rh','Dx112',combined$order)
combined$order<-gsub('lat_fis_ant_horizont_rh','Dx113',combined$order)
combined$order<-gsub('lat_fis_ant_vertical_rh','Dx114',combined$order)
combined$order<-gsub('lat_fis_post_rh','Dx115',combined$order)
combined$order<-gsub('pole_occipital_rh','Dx116',combined$order)
combined$order<-gsub('pole_temporal_rh','Dx117',combined$order)
combined$order<-gsub('s_calcarine_rh','Dx118',combined$order)
combined$order<-gsub('s_central_rh','Dx119',combined$order)
combined$order<-gsub('s_cingul_marginalis_rh','Dx120',combined$order)
combined$order<-gsub('s_circular_insula_ant_rh','Dx121',combined$order)
combined$order<-gsub('s_circular_insula_inf_rh','Dx122',combined$order)
combined$order<-gsub('s_circular_insula_sup_rh','Dx123',combined$order)
combined$order<-gsub('s_collat_transv_ant_rh','Dx124',combined$order)
combined$order<-gsub('s_collat_transv_post_rh','Dx125',combined$order)
combined$order<-gsub('s_front_inf_rh','Dx126',combined$order)
combined$order<-gsub('s_front_middle_rh','Dx127',combined$order)
combined$order<-gsub('s_front_sup_rh','Dx128',combined$order)
combined$order<-gsub('s_interm_prim_jensen_rh','Dx129',combined$order)
combined$order<-gsub('s_intrapariet_and_p_trans_rh','Dx130',combined$order)
combined$order<-gsub('s_oc_middle_and_lunatus_rh','Dx131',combined$order)
combined$order<-gsub('s_oc_sup_and_transversal_rh','Dx132',combined$order)
combined$order<-gsub('s_occipital_ant_rh','Dx133',combined$order)
combined$order<-gsub('s_oc_temp_lat_rh','Dx134',combined$order)
combined$order<-gsub('s_oc_temp_med_and_lingual_rh','Dx135',combined$order)
combined$order<-gsub('s_orbital_lateral_rh','Dx136',combined$order)
combined$order<-gsub('s_orbital_med_olfact_rh','Dx137',combined$order)
combined$order<-gsub('s_orbital_h_shaped_rh','Dx138',combined$order)
combined$order<-gsub('s_parieto_occipital_rh','Dx139',combined$order)
combined$order<-gsub('s_pericallosal_rh','Dx140',combined$order)
combined$order<-gsub('s_postcentral_rh','Dx141',combined$order)
combined$order<-gsub('s_precentral_inf_part_rh','Dx142',combined$order)
combined$order<-gsub('s_precentral_sup_part_rh','Dx143',combined$order)
combined$order<-gsub('s_suborbital_rh','Dx144',combined$order)
combined$order<-gsub('s_subparietal_rh','Dx145',combined$order)
combined$order<-gsub('s_temporal_inf_rh','Dx146',combined$order)
combined$order<-gsub('s_temporal_sup_rh','Dx147',combined$order)
combined$order<-gsub('s_temporal_transverse_rh','Dx148',combined$order)

combined$order<-(gsub('cerebellum_cortex_lh','A01',combined$order))
combined$order<-(gsub('thalamus_proper_lh','A02',combined$order))
combined$order<-(gsub('caudate_lh','A03',combined$order))
combined$order<-(gsub('putamen_lh','A04',combined$order))
combined$order<-(gsub('pallidum_lh','A05',combined$order))
combined$order<-(gsub('brain_stem','A06',combined$order))
combined$order<-(gsub('hippocampus_lh','A07',combined$order))
combined$order<-(gsub('amygdala_lh','A08',combined$order))
combined$order<-(gsub('accumbens_area_lh','A09',combined$order))
combined$order<-(gsub('ventraldc_lh','A10',combined$order))
combined$order<-(gsub('cerebellum_cortex_rh','A11',combined$order))
combined$order<-(gsub('thalamus_proper_rh','A12',combined$order))
combined$order<-(gsub('caudate_rh','A13',combined$order))
combined$order<-(gsub('putamen_rh','A14',combined$order))
combined$order<-(gsub('pallidum_rh','A15',combined$order))
combined$order<-(gsub('hippocampus_rh','A16',combined$order))
combined$order<-(gsub('amygdala_rh','A17',combined$order))
combined$order<-(gsub('accumbens_area_rh','A18',combined$order))
combined$order<-(gsub('ventraldc_rh','A19',combined$order))

combined$order<-(gsub('precuneus_lh','D24',combined$order))
combined$order<-(gsub('precuneus_rh','D58',combined$order))
combined$order<-(gsub('bankssts_lh','D01',combined$order))
combined$order<-(gsub('caudalanteriorcingulate_lh','D02',combined$order))
combined$order<-(gsub('caudalmiddlefrontal_lh','D03',combined$order))
combined$order<-(gsub('cuneus_lh','D04',combined$order))
combined$order<-(gsub('entorhinal_lh','D05',combined$order))
combined$order<-(gsub('fusiform_lh','D06',combined$order))
combined$order<-(gsub('inferiorparietal_lh','D07',combined$order))
combined$order<-(gsub('inferiortemporal_lh','D08',combined$order))
combined$order<-(gsub('isthmuscingulate_lh','D09',combined$order))
combined$order<-(gsub('lateraloccipital_lh','D10',combined$order))
combined$order<-(gsub('lateralorbitofrontal_lh','D11',combined$order))
combined$order<-(gsub('lingual_lh','D12',combined$order))
combined$order<-(gsub('medialorbitofrontal_lh','D13',combined$order))
combined$order<-(gsub('middletemporal_lh','D14',combined$order))
combined$order<-(gsub('parahippocampal_lh','D15',combined$order))
combined$order<-(gsub('paracentral_lh','D16',combined$order))
combined$order<-(gsub('parsopercularis_lh','D17',combined$order))
combined$order<-(gsub('parsorbitalis_lh','D18',combined$order))
combined$order<-(gsub('parstriangularis_lh','D19',combined$order))
combined$order<-(gsub('pericalcarine_lh','D20',combined$order))
combined$order<-(gsub('postcentral_lh','D21',combined$order))
combined$order<-(gsub('posteriorcingulate_lh','D22',combined$order))
combined$order<-(gsub('precentral_lh','D23',combined$order))
combined$order<-(gsub('rostralanteriorcingulate_lh','D25',combined$order))
combined$order<-(gsub('rostralmiddlefrontal_lh','D26',combined$order))
combined$order<-(gsub('superiorfrontal_lh','D27',combined$order))
combined$order<-(gsub('superiorparietal_lh','D28',combined$order))
combined$order<-(gsub('superiortemporal_lh','D29',combined$order))
combined$order<-(gsub('supramarginal_lh','D30',combined$order))
combined$order<-(gsub('frontalpole_lh','D31',combined$order))
combined$order<-(gsub('temporalpole_lh','D32',combined$order))
combined$order<-(gsub('transversetemporal_lh','D33',combined$order))
combined$order<-(gsub('insula_lh','D34',combined$order))
combined$order<-(gsub('bankssts_rh','D35',combined$order))
combined$order<-(gsub('caudalanteriorcingulate_rh','D36',combined$order))
combined$order<-(gsub('caudalmiddlefrontal_rh','D37',combined$order))
combined$order<-(gsub('cuneus_rh','D38',combined$order))
combined$order<-(gsub('entorhinal_rh','D39',combined$order))
combined$order<-(gsub('fusiform_rh','D40',combined$order))
combined$order<-(gsub('inferiorparietal_rh','D41',combined$order))
combined$order<-(gsub('inferiortemporal_rh','D42',combined$order))
combined$order<-(gsub('isthmuscingulate_rh','D43',combined$order))
combined$order<-(gsub('lateraloccipital_rh','D44',combined$order))
combined$order<-(gsub('lateralorbitofrontal_rh','D45',combined$order))
combined$order<-(gsub('lingual_rh','D46',combined$order))
combined$order<-(gsub('medialorbitofrontal_rh','D47',combined$order))
combined$order<-(gsub('middletemporal_rh','D48',combined$order))
combined$order<-(gsub('parahippocampal_rh','D49',combined$order))
combined$order<-(gsub('paracentral_rh','D50',combined$order))
combined$order<-(gsub('parsopercularis_rh','D51',combined$order))
combined$order<-(gsub('parsorbitalis_rh','D52',combined$order))
combined$order<-(gsub('parstriangularis_rh','D53',combined$order))
combined$order<-(gsub('pericalcarine_rh','D54',combined$order))
combined$order<-(gsub('postcentral_rh','D55',combined$order))
combined$order<-(gsub('posteriorcingulate_rh','D56',combined$order))
combined$order<-(gsub('precentral_rh','D57',combined$order))
combined$order<-(gsub('rostralanteriorcingulate_rh','D59',combined$order))
combined$order<-(gsub('rostralmiddlefrontal_rh','D60',combined$order))
combined$order<-(gsub('superiorfrontal_rh','D61',combined$order))
combined$order<-(gsub('superiorparietal_rh','D62',combined$order))
combined$order<-(gsub('superiortemporal_rh','D63',combined$order))
combined$order<-(gsub('supramarginal_rh','D64',combined$order))
combined$order<-(gsub('frontalpole_rh','D65',combined$order))
combined$order<-(gsub('temporalpole_rh','D66',combined$order))
combined$order<-(gsub('transversetemporal_rh','D67',combined$order))
combined$order<-(gsub('insula_rh','D68',combined$order))

combined$Parcellation<-combined$Variable
combined$Parcellation<-(gsub('.*._desikan_.*.','Desikan',combined$Parcellation))
combined$Parcellation<-(gsub('.*._destrieux_.*.','Destrieux',combined$Parcellation))
combined$Parcellation<-(gsub('.*._aseg_.*.','ASEG',combined$Parcellation))

write.csv(combined,outname,row.names = FALSE)

combined<-read.csv(outname)

fdrnames<-c('Contrast','Parcellation','Parcel','OneSTT1 p FDR','OneSTT2 p FDR','OneSTR1T1 p FDR','OneSTR2T1 p FDR','OneSTR1T2 p FDR','OneSTR2T2 p FDR','Paired T Between p FDR','Paired T Within T1 p FDR','Paired T Within T2 p FDR',
            'Paired T Between R1 p FDR','Paired T Between R2 p FDR')
Parcellations<-c('ASEG','Desikan','Destrieux')
contrasts<-unique(combined$Contrast)

combined<-combined[order(combined$order),]
combined<-combined[order(combined$Contrast),]
combined<-combined[order(combined$Parcellation),]

fdrcombined<-as.data.frame(matrix(nrow=0,ncol=14))
names(fdrcombined)<-fdrnames
for(x in Parcellations){
  parcelled<-subset(combined,Parcellation==paste0(x))
  for(y in contrasts){
    tofdr<-subset(parcelled,Contrast==paste0(y))
    fdrcorrected<-as.data.frame(cbind(p.adjust(tofdr$OneSTT1.p,method='fdr',n=nrow(tofdr)),p.adjust(tofdr$OneSTT2.p,method='fdr',n=nrow(tofdr)),p.adjust(tofdr$OneSTR1T1.p,method='fdr',n=nrow(tofdr)),
                                      p.adjust(tofdr$OneSTR2T1.p,method='fdr',n=nrow(tofdr)),p.adjust(tofdr$OneSTR1T2.p,method='fdr',n=nrow(tofdr)),p.adjust(tofdr$OneSTR2T2.p,method='fdr',n=nrow(tofdr)),
                                      p.adjust(tofdr$Paired.T.Between.p,method='fdr',n=nrow(tofdr)),p.adjust(tofdr$Paired.T.Within.T1.p,method='fdr',n=nrow(tofdr)),p.adjust(tofdr$Paired.T.Within.T2.p,method='fdr',n=nrow(tofdr)),
                                      p.adjust(tofdr$Paired.T.Between.R1.p,method='fdr',n=nrow(tofdr)),p.adjust(tofdr$Paired.T.Between.R2.p,method='fdr',n=nrow(tofdr))))
    fdrcorrected<-cbind(y,x,tofdr$Parcel,fdrcorrected)
    names(fdrcorrected)<-fdrnames
    fdrcombined<-rbind(fdrcombined,fdrcorrected)
  }
}
combined<-merge(combined,fdrcombined,by=c("Contrast","Parcellation","Parcel"),all=T)

combined<-combined[order(combined$order),]
combined<-combined[order(combined$Contrast),]
combined<-combined[order(combined$Parcellation),]

write.csv(combined,outname,row.names = FALSE)

combined$OneSTT1.D.Sig<-combined$OneSTT1.D
combined$OneSTT2.D.Sig<-combined$OneSTT2.D
combined$OneSTR1T1.D.Sig<-combined$OneSTR1T1.D
combined$OneSTR2T1.D.Sig<-combined$OneSTR2T1.D
combined$OneSTR1T2.D.Sig<-combined$OneSTR1T2.D
combined$OneSTR2T2.D.Sig<-combined$OneSTR2T2.D
combined$Paired.T.Between.D.Sig<-combined$Paired.T.Between.D
combined$Paired.T.Within.T1.D.Sig<-combined$Paired.T.Within.T1.D
combined$Paired.T.Within.T2.D.Sig<-combined$Paired.T.Within.T2.D
combined$Paired.T.Between.R1.D.Sig<-combined$Paired.T.Between.R1.D
combined$Paired.T.Between.R2.D.Sig<-combined$Paired.T.Between.R2.D

combined$OneSTT1.D.Sig[combined$`OneSTT1 p FDR`>.05]<-0
combined$OneSTT2.D.Sig[combined$`OneSTT2 p FDR`>.05]<-0
combined$OneSTR1T1.D.Sig[combined$`OneSTR1T1 p FDR`>.05]<-0
combined$OneSTR2T1.D.Sig[combined$`OneSTR2T1 p FDR`>.05]<-0
combined$OneSTR1T2.D.Sig[combined$`OneSTR1T2 p FDR`>.05]<-0
combined$OneSTR2T2.D.Sig[combined$`OneSTR2T2 p FDR`>.05]<-0
combined$Paired.T.Between.D.Sig[combined$`Paired T Between p FDR`>.05]<-0
combined$Paired.T.Within.T1.D.Sig[combined$`Paired T Within T1 p FDR`>.05]<-0
combined$Paired.T.Within.T2.D.Sig[combined$`Paired T Within T2 p FDR`>.05]<-0
combined$Paired.T.Between.R1.D.Sig[combined$`Paired T Between R1 p FDR`>.05]<-0
combined$Paired.T.Between.R2.D.Sig[combined$`Paired T Between R2 p FDR`>.05]<-0

combined<-combined[order(combined$order),]
combined<-combined[order(combined$Contrast),]
combined<-combined[order(combined$Parcellation),]

write.csv(combined,outname,row.names = FALSE)

fdrnames<-c('Contrast','Parcel','ASDS OneSTT1 p FDR','ASDS OneSTT2 p FDR','ASDS OneSTR1T1 p FDR','ASDS OneSTR2T1 p FDR','ASDS OneSTR1T2 p FDR','ASDS OneSTR2T2 p FDR','ASDS Paired T Between p FDR','ASDS Paired T Within T1 p FDR','ASDS Paired T Within T2 p FDR',
            'ASDS Paired T Between R1 p FDR','ASDS Paired T Between R2 p FDR')

rm(fdrcorrected)
rm(tofdr)
fdrcombined<-as.data.frame(matrix(nrow=0,ncol=13))
names(fdrcombined)<-fdrnames
asds<-combined[!combined$Parcellation=='Destrieux',]
for(y in contrasts){
  tofdr<-subset(asds,Contrast==paste0(y))
  fdrcorrected<-as.data.frame(cbind(p.adjust(tofdr$OneSTT1.p,method='fdr',n=nrow(tofdr)),p.adjust(tofdr$OneSTT2.p,method='fdr',n=nrow(tofdr)),p.adjust(tofdr$OneSTR1T1.p,method='fdr',n=nrow(tofdr)),
                                      p.adjust(tofdr$OneSTR2T1.p,method='fdr',n=nrow(tofdr)),p.adjust(tofdr$OneSTR1T2.p,method='fdr',n=nrow(tofdr)),p.adjust(tofdr$OneSTR2T2.p,method='fdr',n=nrow(tofdr)),
                                      p.adjust(tofdr$Paired.T.Between.p,method='fdr',n=nrow(tofdr)),p.adjust(tofdr$Paired.T.Within.T1.p,method='fdr',n=nrow(tofdr)),p.adjust(tofdr$Paired.T.Within.T2.p,method='fdr',n=nrow(tofdr)),
                                      p.adjust(tofdr$Paired.T.Between.R1.p,method='fdr',n=nrow(tofdr)),p.adjust(tofdr$Paired.T.Between.R2.p,method='fdr',n=nrow(tofdr))))
  fdrcorrected<-cbind(y,tofdr$Parcel,fdrcorrected)
  names(fdrcorrected)<-fdrnames
  fdrcombined<-rbind(fdrcombined,fdrcorrected)
}

combined<-merge(combined,fdrcombined,by=c("Contrast","Parcel"),all=T)

combined<-combined[order(combined$order),]
combined<-combined[order(combined$Contrast),]
combined<-combined[order(combined$Parcellation),]

write.csv(combined,outname,row.names = FALSE)

combined$ASDSOneSTT1.D.Sig<-combined$OneSTT1.D
combined$ASDSOneSTT2.D.Sig<-combined$OneSTT2.D
combined$ASDSOneSTR1T1.D.Sig<-combined$OneSTR1T1.D
combined$ASDSOneSTR2T1.D.Sig<-combined$OneSTR2T1.D
combined$ASDSOneSTR1T2.D.Sig<-combined$OneSTR1T2.D
combined$ASDSOneSTR2T2.D.Sig<-combined$OneSTR2T2.D
combined$ASDSPaired.T.Between.D.Sig<-combined$Paired.T.Between.D
combined$ASDSPaired.T.Within.T1.D.Sig<-combined$Paired.T.Within.T1.D
combined$ASDSPaired.T.Within.T2.D.Sig<-combined$Paired.T.Within.T2.D
combined$ASDSPaired.T.Between.R1.D.Sig<-combined$Paired.T.Between.R1.D
combined$ASDSPaired.T.Between.R2.D.Sig<-combined$Paired.T.Between.R2.D

combined$ASDSOneSTT1.D.Sig[combined$`ASDS OneSTT1 p FDR`>.05]<-0
combined$ASDSOneSTT2.D.Sig[combined$`ASDS OneSTT2 p FDR`>.05]<-0
combined$ASDSOneSTR1T1.D.Sig[combined$`ASDS OneSTR1T1 p FDR`>.05]<-0
combined$ASDSOneSTR2T1.D.Sig[combined$`ASDS OneSTR2T1 p FDR`>.05]<-0
combined$ASDSOneSTR1T2.D.Sig[combined$`ASDS OneSTR1T2 p FDR`>.05]<-0
combined$ASDSOneSTR2T2.D.Sig[combined$`ASDS OneSTR2T2 p FDR`>.05]<-0
combined$ASDSPaired.T.Between.D.Sig[combined$`ASDS Paired T Between p FDR`>.05]<-0
combined$ASDSPaired.T.Within.T1.D.Sig[combined$`ASDS Paired T Within T1 p FDR`>.05]<-0
combined$ASDSPaired.T.Within.T2.D.Sig[combined$`ASDS Paired T Within T2 p FDR`>.05]<-0
combined$ASDSPaired.T.Between.R1.D.Sig[combined$`ASDS Paired T Between R1 p FDR`>.05]<-0
combined$ASDSPaired.T.Between.R2.D.Sig[combined$`ASDS Paired T Between R2 p FDR`>.05]<-0

combined<-combined[order(combined$order),]
combined<-combined[order(combined$Contrast),]
combined<-combined[order(combined$Parcellation),]

write.csv(combined,outname,row.names = FALSE)

fdrnames<-c('Contrast','Parcel','ASDX OneSTT1 p FDR','ASDX OneSTT2 p FDR','ASDX OneSTR1T1 p FDR','ASDX OneSTR2T1 p FDR','ASDX OneSTR1T2 p FDR','ASDX OneSTR2T2 p FDR','ASDX Paired T Between p FDR','ASDX Paired T Within T1 p FDR','ASDX Paired T Within T2 p FDR',
            'ASDX Paired T Between R1 p FDR','ASDX Paired T Between R2 p FDR')

fdrcombined<-as.data.frame(matrix(nrow=0,ncol=13))
names(fdrcombined)<-fdrnames
rm(fdrcorrected)
rm(tofdr)
asdx<-combined[!combined$Parcellation=='Desikan',]
for(y in contrasts){
  tofdr<-subset(asdx,Contrast==paste0(y))
  fdrcorrected<-as.data.frame(cbind(p.adjust(tofdr$OneSTT1.p,method='fdr',n=nrow(tofdr)),p.adjust(tofdr$OneSTT2.p,method='fdr',n=nrow(tofdr)),p.adjust(tofdr$OneSTR1T1.p,method='fdr',n=nrow(tofdr)),
                                    p.adjust(tofdr$OneSTR2T1.p,method='fdr',n=nrow(tofdr)),p.adjust(tofdr$OneSTR1T2.p,method='fdr',n=nrow(tofdr)),p.adjust(tofdr$OneSTR2T2.p,method='fdr',n=nrow(tofdr)),
                                    p.adjust(tofdr$Paired.T.Between.p,method='fdr',n=nrow(tofdr)),p.adjust(tofdr$Paired.T.Within.T1.p,method='fdr',n=nrow(tofdr)),p.adjust(tofdr$Paired.T.Within.T2.p,method='fdr',n=nrow(tofdr)),
                                    p.adjust(tofdr$Paired.T.Between.R1.p,method='fdr',n=nrow(tofdr)),p.adjust(tofdr$Paired.T.Between.R2.p,method='fdr',n=nrow(tofdr))))
  fdrcorrected<-cbind(y,tofdr$Parcel,fdrcorrected)
  names(fdrcorrected)<-fdrnames
  fdrcombined<-rbind(fdrcombined,fdrcorrected)
}

combined<-merge(combined,fdrcombined,by=c("Contrast","Parcel"),all=T)

combined<-combined[order(combined$order),]
combined<-combined[order(combined$Contrast),]
combined<-combined[order(combined$Parcellation),]

write.csv(combined,outname,row.names = FALSE)

combined$ASDXOneSTT1.D.Sig<-combined$OneSTT1.D
combined$ASDXOneSTT2.D.Sig<-combined$OneSTT2.D
combined$ASDXOneSTR1T1.D.Sig<-combined$OneSTR1T1.D
combined$ASDXOneSTR2T1.D.Sig<-combined$OneSTR2T1.D
combined$ASDXOneSTR1T2.D.Sig<-combined$OneSTR1T2.D
combined$ASDXOneSTR2T2.D.Sig<-combined$OneSTR2T2.D
combined$ASDXPaired.T.Between.D.Sig<-combined$Paired.T.Between.D
combined$ASDXPaired.T.Within.T1.D.Sig<-combined$Paired.T.Within.T1.D
combined$ASDXPaired.T.Within.T2.D.Sig<-combined$Paired.T.Within.T2.D
combined$ASDXPaired.T.Between.R1.D.Sig<-combined$Paired.T.Between.R1.D
combined$ASDXPaired.T.Between.R2.D.Sig<-combined$Paired.T.Between.R2.D

combined$ASDXOneSTT1.D.Sig[combined$`ASDX OneSTT1 p FDR`>.05]<-0
combined$ASDXOneSTT2.D.Sig[combined$`ASDX OneSTT2 p FDR`>.05]<-0
combined$ASDXOneSTR1T1.D.Sig[combined$`ASDX OneSTR1T1 p FDR`>.05]<-0
combined$ASDXOneSTR2T1.D.Sig[combined$`ASDX OneSTR2T1 p FDR`>.05]<-0
combined$ASDXOneSTR1T2.D.Sig[combined$`ASDX OneSTR1T2 p FDR`>.05]<-0
combined$ASDXOneSTR2T2.D.Sig[combined$`ASDX OneSTR2T2 p FDR`>.05]<-0
combined$ASDXPaired.T.Between.D.Sig[combined$`ASDX Paired T Between p FDR`>.05]<-0
combined$ASDXPaired.T.Within.T1.D.Sig[combined$`ASDX Paired T Within T1 p FDR`>.05]<-0
combined$ASDXPaired.T.Within.T2.D.Sig[combined$`ASDX Paired T Within T2 p FDR`>.05]<-0
combined$ASDXPaired.T.Between.R1.D.Sig[combined$`ASDX Paired T Between R1 p FDR`>.05]<-0
combined$ASDXPaired.T.Between.R2.D.Sig[combined$`ASDX Paired T Between R2 p FDR`>.05]<-0

combined<-combined[order(combined$order),]
combined<-combined[order(combined$Contrast),]
combined<-combined[order(combined$Parcellation),]

combined$ASDSOneSTT1.D.Sig[is.na(combined$`ASDS OneSTT1 p FDR`)]<-0
combined$ASDSOneSTT2.D.Sig[is.na(combined$`ASDS OneSTT2 p FDR`)]<-0
combined$ASDSOneSTR1T1.D.Sig[is.na(combined$`ASDS OneSTR1T1 p FDR`)]<-0
combined$ASDSOneSTR2T1.D.Sig[is.na(combined$`ASDS OneSTR2T1 p FDR`)]<-0
combined$ASDSOneSTR1T2.D.Sig[is.na(combined$`ASDS OneSTR1T2 p FDR`)]<-0
combined$ASDSOneSTR2T2.D.Sig[is.na(combined$`ASDS OneSTR2T2 p FDR`)]<-0
combined$ASDSPaired.T.Between.D.Sig[is.na(combined$`ASDS Paired T Between p FDR`)]<-0
combined$ASDSPaired.T.Within.T1.D.Sig[is.na(combined$`ASDS Paired T Within T1 p FDR`)]<-0
combined$ASDSPaired.T.Within.T2.D.Sig[is.na(combined$`ASDS Paired T Within T2 p FDR`)]<-0
combined$ASDSPaired.T.Between.R1.D.Sig[is.na(combined$`ASDS Paired T Between R1 p FDR`)]<-0
combined$ASDSPaired.T.Between.R2.D.Sig[is.na(combined$`ASDS Paired T Between R2 p FDR`)]<-0

combined$ASDXOneSTT1.D.Sig[is.na(combined$`ASDX OneSTT1 p FDR`)]<-0
combined$ASDXOneSTT2.D.Sig[is.na(combined$`ASDX OneSTT2 p FDR`)]<-0
combined$ASDXOneSTR1T1.D.Sig[is.na(combined$`ASDX OneSTR1T1 p FDR`)]<-0
combined$ASDXOneSTR2T1.D.Sig[is.na(combined$`ASDX OneSTR2T1 p FDR`)]<-0
combined$ASDXOneSTR1T2.D.Sig[is.na(combined$`ASDX OneSTR1T2 p FDR`)]<-0
combined$ASDXOneSTR2T2.D.Sig[is.na(combined$`ASDX OneSTR2T2 p FDR`)]<-0
combined$ASDXPaired.T.Between.D.Sig[is.na(combined$`ASDX Paired T Between p FDR`)]<-0
combined$ASDXPaired.T.Within.T1.D.Sig[is.na(combined$`ASDX Paired T Within T1 p FDR`)]<-0
combined$ASDXPaired.T.Within.T2.D.Sig[is.na(combined$`ASDX Paired T Within T2 p FDR`)]<-0
combined$ASDXPaired.T.Between.R1.D.Sig[is.na(combined$`ASDX Paired T Between R1 p FDR`)]<-0
combined$ASDXPaired.T.Between.R2.D.Sig[is.na(combined$`ASDX Paired T Between R2 p FDR`)]<-0

write.csv(combined,outname,row.names = FALSE)
```
