## Supplementary material for "Reliability and Stability Challenges in ABCD Task fMRI Data": Code for generating data and results: Script 3 - DifferenceReliabilityCI.R

```
#ICC, activity, and change calculations.
#Set the working directory to the previously created ICC folder (line 10)
#Choose the sample you're interested in in the list.files pattern (line 11, currently searching for 'usable'), edit the output name (line 12),
#Select all and run and it will calculate values for all variables

if('irr' %in% rownames(installed.packages())==FALSE){
  install.packages('irr')}
library('irr')
library('psych')
setwd("C:/Users/James/Desktop/Release 3 Nov 2020/ICC")
measures<-list.files(pattern='^Outlier_.*.R1R2.*.csv$')
outname<-'ICCs_Difference.csv'
combOut<-matrix(ncol=109,nrow=0)
for(i in measures){
  data<-read.csv(i, stringsAsFactors = F)
  measureOut<-paste0('ICC(3,2)_Difference',i)
  datanames<-names(data)
  datanamesstripped<-gsub("_run1_",'RunSame',datanames)
  datanamesstripped<-gsub("_run2_",'RunSame',datanamesstripped)
  datanamesstripped<-gsub("_0m",'',datanamesstripped)
  datanamesstripped<-gsub("_24m",'',datanamesstripped)
  usablevariables<-datanamesstripped[duplicated(datanamesstripped)]
  usablevariables<-unique(usablevariables)
  names(data)[1]<-'ID'
  output<-matrix(ncol=108,nrow=0)
  for(z in usablevariables){
    varnameR1T1<-paste0(gsub('RunSame',"_run1_",z),"_0m")
    varnameR1T2<-paste0(gsub('RunSame',"_run1_",z),"_24m")
    varnameR2T1<-paste0(gsub('RunSame',"_run2_",z),"_0m")
    varnameR2T2<-paste0(gsub('RunSame',"_run2_",z),"_24m")    
    datause<-cbind(data$ID,data[[varnameR1T1]],data[[varnameR1T2]],data[[varnameR2T1]],data[[varnameR2T2]])
    datause<-as.data.frame(datause)
    names(datause)<-c('ID','R1T1','R1T2','R2T1','R2T2')
    datause$R1T1<-as.numeric(datause$R1T1)
    datause$R1T2<-as.numeric(datause$R1T2)
    datause$R2T1<-as.numeric(datause$R2T1)
    datause$R2T2<-as.numeric(datause$R2T2)
    datause$WT1Diff<-datause$R2T1-datause$R1T1
    datause$WT2Diff<-datause$R2T2-datause$R1T2
    datause$BTR1Diff<-datause$R1T2-datause$R1T1
    datause$BTR2Diff<-datause$R2T2-datause$R2T1
    
    iccWT<-icc(cbind(datause$WT1Diff,datause$WT2Diff), model = 'twoway', type = 'consistency', unit='single')
    iccBTR<-icc(cbind(datause$BTR1Diff,datause$BTR2Diff), model = 'twoway', type = 'consistency', unit='single')
    
    oneSTWT1Diff<-t.test(datause$WT1Diff,mu=0)
    oneSTWT2Diff<-t.test(datause$WT2Diff,mu=0)
    
    oneSTBTR1Diff<-t.test(datause$BTR1Diff,mu=0)
    oneSTBTR2Diff<-t.test(datause$BTR2Diff,mu=0)
    
    oneSDWT1Diff<-(mean(datause$WT1Diff,na.rm=T)/(sd(datause$WT1Diff,na.rm=T)))
    oneSDWT1DiffCI<-cohen.d.ci(d=oneSDWT1Diff,n=length(complete.cases(datause$WT1Diff)))
    oneSDWT2Diff<-(mean(datause$WT2Diff,na.rm=T)/(sd(datause$WT2Diff,na.rm=T)))
    oneSDWT2DiffCI<-cohen.d.ci(d=oneSDWT2Diff,n=length(complete.cases(datause$WT2Diff)))
    
    oneSDBTR1Diff<-(mean(datause$BTR1Diff,na.rm=T)/(sd(datause$BTR1Diff,na.rm=T)))
    oneSDBTRDiffCI<-cohen.d.ci(d=oneSDBTR1Diff,n=length(complete.cases(datause$BTR1Diff)))
    oneSDBTR2Diff<-(mean(datause$BTR2Diff,na.rm=T)/(sd(datause$BTR2Diff,na.rm=T)))
    oneSDBTR2DiffCI<-cohen.d.ci(d=oneSDBTR2Diff,n=length(complete.cases(datause$BTR2Diff)))
    
    pairedtWTDiff<-t.test(datause$WT2Diff,datause$WT1Diff,paired=T)
    pairedtDWTDiff<-((mean(datause$WT2Diff-datause$WT1Diff,na.rm = T))/sd(datause$WT2Diff-datause$WT1Diff,na.rm = T))
    pairedtDWTDiffCI<-cohen.d.ci(d=pairedtDWTDiff,n1=length(complete.cases(datause$WT1Diff)),n2=length(complete.cases(datause$WT2Diff)))
    
    pairedtBTRDiff<-t.test(datause$BTR2Diff,datause$BTR1Diff,paired=T)
    pairedtDBTRDiff<-((mean(datause$BTR2Diff-datause$BTR1Diff,na.rm = T))/sd(datause$BTR2Diff-datause$BTR1Diff,na.rm = T))
    pairedtDBTRDiffCI<-cohen.d.ci(d=pairedtDBTRDiff,n1=length(complete.cases(datause$BTR1Diff)),n2=length(complete.cases(datause$BTR2Diff)))

    betweendata<-datause[,c(1,8,9)]
    betweendata<-betweendata[complete.cases(betweendata),]
    betweenSStotal<-(var(unlist(betweendata[,2:3]))*((nrow(betweendata)*(ncol(betweendata)-1))-1))
    betweenMSR<-var(rowMeans(betweendata[,2:3]))*(ncol(betweendata)-1)
    betweenMSC<-var(colMeans(betweendata[,2:3]))*nrow(betweendata)
    betweenMSE<-((betweenSStotal-betweenMSR*(nrow(betweendata)-1)-betweenMSC*(ncol(betweendata)-2)))/((nrow(betweendata)-1)*(ncol(betweendata)-2))
    withindata<-datause[,c(1,6,7)]
    withindata<-withindata[complete.cases(withindata),]
    withinSStotal<-(var(unlist(withindata[,2:3]))*((nrow(withindata)*(ncol(withindata)-1))-1))
    withinMSR<-var(rowMeans(withindata[,2:3]))*(ncol(withindata)-1)
    withinMSC<-var(colMeans(withindata[,2:3]))*nrow(withindata)
    withinMSE<-((withinSStotal-withinMSR*(nrow(withindata)-1)-withinMSC*(ncol(withindata)-2)))/((nrow(withindata)-1)*(ncol(withindata)-2))
    
    results<-cbind(z,iccWT$value,iccWT$lbound,iccWT$ubound,iccBTR$value,iccBTR$lbound,iccBTR$ubound,
                   oneSTWT1Diff$statistic,oneSTWT1Diff$p.value,oneSTWT1Diff$conf.int[1],oneSTWT1Diff$conf.int[2],
                   oneSTWT2Diff$statistic,oneSTWT2Diff$p.value,oneSTWT2Diff$conf.int[1],oneSTWT2Diff$conf.int[2],
                   oneSTBTR1Diff$statistic,oneSTBTR1Diff$p.value,oneSTBTR1Diff$conf.int[1],oneSTBTR1Diff$conf.int[2],
                   oneSTBTR2Diff$statistic,oneSTBTR2Diff$p.value,oneSTBTR2Diff$conf.int[1],oneSTBTR2Diff$conf.int[2],
                   oneSDWT1Diff,oneSDWT1DiffCI[1,1],oneSDWT1DiffCI[1,3],oneSDWT2Diff,oneSDWT2DiffCI[1,1],oneSDWT2DiffCI[1,3],
                   oneSDBTR1Diff,oneSDBTR1DiffCI[1,1],oneSDBTR1DiffCI[1,3],oneSDBTR2Diff,oneSDBTR2DiffCI[1,1],oneSDBTR2DiffCI[1,3],
                   pairedtWTDiff$statistic,pairedtWTDiff$p.value,pairedtWTDiff$conf.int[1],pairedtWTDiff$conf.int[2],
                   pairedtBTRDiff$statistic,pairedtBTRDiff$p.value,pairedtBTRDiff$conf.int[1],pairedtBTRDiff$conf.int[2],
                   pairedtDWTDiff,pairedtDWTDiffCI[1,1],pairedtDWTDiffCI[1,3],
                   pairedtDBTRDiff,pairedtDBTRDiffCI[1,1],pairedtDBTRDiffCI[1,3],
                   betweenMSR,betweenMSC,betweenMSE,withinMSR,withinMSC,withinMSE)
    output<-rbind(output,results)
  }
  combOut<-rbind(combOut,cbind(i,output))
  output<-as.data.frame(output)
  names(output)<-c('Variable','ICC Within Diff','ICC Within Diff 95% CI Lower','ICC Within Diff 95% CI Upper',
                   'ICC Between Run Diff','ICC Between Run Diff 95% CI Lower','ICC Between Run Diff 95% CI Upper',
                   'OneST Within T1 Diff t','OneST Within T1 Diff p','OneST Within T1 Diff 95% CI Lower Mean','OneST Within T1 Diff 95% CI Upper Mean',
                   'OneST Within T2 Diff t','OneST Within T2 Diff p','OneST Within T2 Diff 95% CI Lower Mean','OneST Within T2 Diff 95% CI Upper Mean',
                   'OneST Betwen Run R1 Diff t','OneST Between Run R1 Diff p','OneST Between Run R1 Diff 95% CI Lower Mean','OneST Between Run R1 Diff 95% CI Upper Mean',
                   'OneST Betwen Run R2 Diff t','OneST Between Run R2 Diff p','OneST Between Run R2 Diff 95% CI Lower Mean','OneST Between Run R2 Diff 95% CI Upper Mean',
                   'OneST Within T1 Diff D','OneST Within T1 Diff D 95% CI Lower','OneST Within T1 Diff D 95% CI Upper',
                   'OneST Within T2 Diff D','OneST Within T2 Diff D 95% CI Lower','OneST Within T2 Diff D 95% CI Upper',
                   'OneST Between Run R1 Diff D','OneST Between Run R1 Diff D 95% CI Lower','OneST Between Run R1 Diff D 95% CI Upper',
                   'OneST Between Run R2 Diff D','OneST Between Run R2 Diff D 95% CI Lower','OneST Between Run R2 Diff D 95% CI Upper',
                   'Paired T Within Diff t','Paired T Within Diff p','Paired T Within Diff 95% CI Lower Difference','Paired T Within Diff 95% CI Upper Difference',
                   'Paired T Between Diff t','Paired T Between Diff p','Paired T Between Diff 95% CI Lower Difference','Paired T Between Diff 95% CI Upper Difference',
                   'Paired T Within Diff D','Paired T Within Diff D 95% CI Lower','Paired T Within Diff D 95% CI Upper',
                   'Paired T Between Diff D','Paired T Between Diff D 95% CI Lower','Paired T Between Diff D 95% CI Upper',
                   'Between Diff MSR','Between Diff MSC','Between Diff MSE','Within Diff MSR','Within Diff MSC','Within Diff MSE')
  write.csv(output,measureOut,row.names = FALSE)
}
combOut<-as.data.frame(combOut)
names(combOut)<-c('File','Variable','ICC Within Diff','ICC Within Diff 95% CI Lower','ICC Within Diff 95% CI Upper',
                  'ICC Between Run Diff','ICC Between Run Diff 95% CI Lower','ICC Between Run Diff 95% CI Upper',
                  'OneST Within T1 Diff t','OneST Within T1 Diff p','OneST Within T1 Diff 95% CI Lower Mean','OneST Within T1 Diff 95% CI Upper Mean',
                  'OneST Within T2 Diff t','OneST Within T2 Diff p','OneST Within T2 Diff 95% CI Lower Mean','OneST Within T2 Diff 95% CI Upper Mean',
                  'OneST Betwen Run R1 Diff t','OneST Between Run R1 Diff p','OneST Between Run R1 Diff 95% CI Lower Mean','OneST Between Run R1 Diff 95% CI Upper Mean',
                  'OneST Betwen Run R2 Diff t','OneST Between Run R2 Diff p','OneST Between Run R2 Diff 95% CI Lower Mean','OneST Between Run R2 Diff 95% CI Upper Mean',
                  'OneST Within T1 Diff D','OneST Within T1 Diff D 95% CI Lower','OneST Within T1 Diff D 95% CI Upper',
                  'OneST Within T2 Diff D','OneST Within T2 Diff D 95% CI Lower','OneST Within T2 Diff D 95% CI Upper',
                  'OneST Between Run R1 Diff D','OneST Between Run R1 Diff D 95% CI Lower','OneST Between Run R1 Diff D 95% CI Upper',
                  'OneST Between Run R2 Diff D','OneST Between Run R2 Diff D 95% CI Lower','OneST Between Run R2 Diff D 95% CI Upper',
                  'Paired T Within Diff t','Paired T Within Diff p','Paired T Within Diff 95% CI Lower Difference','Paired T Within Diff 95% CI Upper Difference',
                  'Paired T Between Diff t','Paired T Between Diff p','Paired T Between Diff 95% CI Lower Difference','Paired T Between Diff 95% CI Upper Difference',
                  'Paired T Within Diff D','Paired T Within Diff D 95% CI Lower','Paired T Within Diff D 95% CI Upper',
                  'Paired T Between Diff D','Paired T Between Diff D 95% CI Lower','Paired T Between Diff D 95% CI Upper',
                  'Between Diff MSR','Between Diff MSC','Between Diff MSE','Within Diff MSR','Within Diff MSC','Within Diff MSE')
write.csv(combOut,outname,row.names = FALSE)

combined<-combOut

combined$Contrast<-combined$Variable
combined$Contrast<-gsub('_beta_.*.','',combined$Contrast)
combined$Contrast<-gsub('.*._all_','',combined$Contrast)
combined$Contrast<-gsub('.*.RunSame','',combined$Contrast)

combined$Parcel<-combined$Variable
combined$Parcel<-gsub('.*._aseg_','',combined$Parcel)
combined$Parcel<-gsub('.*._destrieux_','',combined$Parcel)
combined$Parcel<-gsub('.*._desikan_','',combined$Parcel)
combined$Parcel<-gsub('RunSame','',combined$Parcel)

combined<-combined[!(combined$Parcel=='cerebral_white_matter_lh'),]
combined<-combined[!(combined$Parcel=='cerebral_white_matter_rh'),]
combined<-combined[!(combined$Parcel=='cerebellum_white_matter_lh'),]
combined<-combined[!(combined$Parcel=='cerebellum_white_matter_rh'),]
combined<-combined[!(combined$Parcel=='lateral_ventricle_lh'),]
combined<-combined[!(combined$Parcel=='lateral_ventricle_rh'),]
combined<-combined[!(combined$Parcel=='inf_lat_vent_lh'),]
combined<-combined[!(combined$Parcel=='inf_lat_vent_rh'),]
combined<-combined[!(combined$Parcel=='csf'),]
combined<-combined[!(combined$Parcel=='3rd_ventricle'),]
combined<-combined[!(combined$Parcel=='4th_ventricle'),]

combined<-combined[!(combined$Contrast==''),]

combined$order<-combined$Parcel
combined$order<-gsub('g_and_s_frontomargin_lh','Dx001',combined$order)
combined$order<-gsub('g_and_s_occipital_inf_lh','Dx002',combined$order)
combined$order<-gsub('g_and_s_paracentral_lh','Dx003',combined$order)
combined$order<-gsub('g_and_s_subcentral_lh','Dx004',combined$order)
combined$order<-gsub('g_and_s_transv_frontopol_lh','Dx005',combined$order)
combined$order<-gsub('g_and_s_cingul_ant_lh','Dx006',combined$order)
combined$order<-gsub('g_and_s_cingul_mid_ant_lh','Dx007',combined$order)
combined$order<-gsub('g_and_s_cingul_mid_post_lh','Dx008',combined$order)
combined$order<-gsub('g_cingul_post_dorsal_lh','Dx009',combined$order)
combined$order<-gsub('g_cingul_post_ventral_lh','Dx010',combined$order)
combined$order<-gsub('g_cuneus_lh','Dx011',combined$order)
combined$order<-gsub('g_front_inf_opercular_lh','Dx012',combined$order)
combined$order<-gsub('g_front_inf_orbital_lh','Dx013',combined$order)
combined$order<-gsub('g_front_inf_triangul_lh','Dx014',combined$order)
combined$order<-gsub('g_front_middle_lh','Dx015',combined$order)
combined$order<-gsub('g_front_sup_lh','Dx016',combined$order)
combined$order<-gsub('g_ins_lg_and_s_cent_ins_lh','Dx017',combined$order)
combined$order<-gsub('g_insular_short_lh','Dx018',combined$order)
combined$order<-gsub('g_occipital_middle_lh','Dx019',combined$order)
combined$order<-gsub('g_occipital_sup_lh','Dx020',combined$order)
combined$order<-gsub('g_oc_temp_lat_fusifor_lh','Dx021',combined$order)
combined$order<-gsub('g_oc_temp_med_lingual_lh','Dx022',combined$order)
combined$order<-gsub('g_oc_temp_med_parahip_lh','Dx023',combined$order)
combined$order<-gsub('g_orbital_lh','Dx024',combined$order)
combined$order<-gsub('g_pariet_inf_angular_lh','Dx025',combined$order)
combined$order<-gsub('g_pariet_inf_supramar_lh','Dx026',combined$order)
combined$order<-gsub('g_parietal_sup_lh','Dx027',combined$order)
combined$order<-gsub('g_postcentral_lh','Dx028',combined$order)
combined$order<-gsub('g_precentral_lh','Dx029',combined$order)
combined$order<-gsub('g_precuneus_lh','Dx030',combined$order)
combined$order<-gsub('g_rectus_lh','Dx031',combined$order)
combined$order<-gsub('g_subcallosal_lh','Dx032',combined$order)
combined$order<-gsub('g_temp_sup_g_t_transv_lh','Dx033',combined$order)
combined$order<-gsub('g_temp_sup_lateral_lh','Dx034',combined$order)
combined$order<-gsub('g_temp_sup_plan_polar_lh','Dx035',combined$order)
combined$order<-gsub('g_temp_sup_plan_tempo_lh','Dx036',combined$order)
combined$order<-gsub('g_temporal_inf_lh','Dx037',combined$order)
combined$order<-gsub('g_temporal_middle_lh','Dx038',combined$order)
combined$order<-gsub('lat_fis_ant_horizont_lh','Dx039',combined$order)
combined$order<-gsub('lat_fis_ant_vertical_lh','Dx040',combined$order)
combined$order<-gsub('lat_fis_post_lh','Dx041',combined$order)
combined$order<-gsub('pole_occipital_lh','Dx042',combined$order)
combined$order<-gsub('pole_temporal_lh','Dx043',combined$order)
combined$order<-gsub('s_calcarine_lh','Dx044',combined$order)
combined$order<-gsub('s_central_lh','Dx045',combined$order)
combined$order<-gsub('s_cingul_marginalis_lh','Dx046',combined$order)
combined$order<-gsub('s_circular_insula_ant_lh','Dx047',combined$order)
combined$order<-gsub('s_circular_insula_inf_lh','Dx048',combined$order)
combined$order<-gsub('s_circular_insula_sup_lh','Dx049',combined$order)
combined$order<-gsub('s_collat_transv_ant_lh','Dx050',combined$order)
combined$order<-gsub('s_collat_transv_post_lh','Dx051',combined$order)
combined$order<-gsub('s_front_inf_lh','Dx052',combined$order)
combined$order<-gsub('s_front_middle_lh','Dx053',combined$order)
combined$order<-gsub('s_front_sup_lh','Dx054',combined$order)
combined$order<-gsub('s_interm_prim_jensen_lh','Dx055',combined$order)
combined$order<-gsub('s_intrapariet_and_p_trans_lh','Dx056',combined$order)
combined$order<-gsub('s_oc_middle_and_lunatus_lh','Dx057',combined$order)
combined$order<-gsub('s_oc_sup_and_transversal_lh','Dx058',combined$order)
combined$order<-gsub('s_occipital_ant_lh','Dx059',combined$order)
combined$order<-gsub('s_oc_temp_lat_lh','Dx060',combined$order)
combined$order<-gsub('s_oc_temp_med_and_lingual_lh','Dx061',combined$order)
combined$order<-gsub('s_orbital_lateral_lh','Dx062',combined$order)
combined$order<-gsub('s_orbital_med_olfact_lh','Dx063',combined$order)
combined$order<-gsub('s_orbital_h_shaped_lh','Dx064',combined$order)
combined$order<-gsub('s_parieto_occipital_lh','Dx065',combined$order)
combined$order<-gsub('s_pericallosal_lh','Dx066',combined$order)
combined$order<-gsub('s_postcentral_lh','Dx067',combined$order)
combined$order<-gsub('s_precentral_inf_part_lh','Dx068',combined$order)
combined$order<-gsub('s_precentral_sup_part_lh','Dx069',combined$order)
combined$order<-gsub('s_suborbital_lh','Dx070',combined$order)
combined$order<-gsub('s_subparietal_lh','Dx071',combined$order)
combined$order<-gsub('s_temporal_inf_lh','Dx072',combined$order)
combined$order<-gsub('s_temporal_sup_lh','Dx073',combined$order)
combined$order<-gsub('s_temporal_transverse_lh','Dx074',combined$order)
combined$order<-gsub('g_and_s_frontomargin_rh','Dx075',combined$order)
combined$order<-gsub('g_and_s_occipital_inf_rh','Dx076',combined$order)
combined$order<-gsub('g_and_s_paracentral_rh','Dx077',combined$order)
combined$order<-gsub('g_and_s_subcentral_rh','Dx078',combined$order)
combined$order<-gsub('g_and_s_transv_frontopol_rh','Dx079',combined$order)
combined$order<-gsub('g_and_s_cingul_ant_rh','Dx080',combined$order)
combined$order<-gsub('g_and_s_cingul_mid_ant_rh','Dx081',combined$order)
combined$order<-gsub('g_and_s_cingul_mid_post_rh','Dx082',combined$order)
combined$order<-gsub('g_cingul_post_dorsal_rh','Dx083',combined$order)
combined$order<-gsub('g_cingul_post_ventral_rh','Dx084',combined$order)
combined$order<-gsub('g_cuneus_rh','Dx085',combined$order)
combined$order<-gsub('g_front_inf_opercular_rh','Dx086',combined$order)
combined$order<-gsub('g_front_inf_orbital_rh','Dx087',combined$order)
combined$order<-gsub('g_front_inf_triangul_rh','Dx088',combined$order)
combined$order<-gsub('g_front_middle_rh','Dx089',combined$order)
combined$order<-gsub('g_front_sup_rh','Dx090',combined$order)
combined$order<-gsub('g_ins_lg_and_s_cent_ins_rh','Dx091',combined$order)
combined$order<-gsub('g_insular_short_rh','Dx092',combined$order)
combined$order<-gsub('g_occipital_middle_rh','Dx093',combined$order)
combined$order<-gsub('g_occipital_sup_rh','Dx094',combined$order)
combined$order<-gsub('g_oc_temp_lat_fusifor_rh','Dx095',combined$order)
combined$order<-gsub('g_oc_temp_med_lingual_rh','Dx096',combined$order)
combined$order<-gsub('g_oc_temp_med_parahip_rh','Dx097',combined$order)
combined$order<-gsub('g_orbital_rh','Dx098',combined$order)
combined$order<-gsub('g_pariet_inf_angular_rh','Dx099',combined$order)
combined$order<-gsub('g_pariet_inf_supramar_rh','Dx100',combined$order)
combined$order<-gsub('g_parietal_sup_rh','Dx101',combined$order)
combined$order<-gsub('g_postcentral_rh','Dx102',combined$order)
combined$order<-gsub('g_precentral_rh','Dx103',combined$order)
combined$order<-gsub('g_precuneus_rh','Dx104',combined$order)
combined$order<-gsub('g_rectus_rh','Dx105',combined$order)
combined$order<-gsub('g_subcallosal_rh','Dx106',combined$order)
combined$order<-gsub('g_temp_sup_g_t_transv_rh','Dx107',combined$order)
combined$order<-gsub('g_temp_sup_lateral_rh','Dx108',combined$order)
combined$order<-gsub('g_temp_sup_plan_polar_rh','Dx109',combined$order)
combined$order<-gsub('g_temp_sup_plan_tempo_rh','Dx110',combined$order)
combined$order<-gsub('g_temporal_inf_rh','Dx111',combined$order)
combined$order<-gsub('g_temporal_middle_rh','Dx112',combined$order)
combined$order<-gsub('lat_fis_ant_horizont_rh','Dx113',combined$order)
combined$order<-gsub('lat_fis_ant_vertical_rh','Dx114',combined$order)
combined$order<-gsub('lat_fis_post_rh','Dx115',combined$order)
combined$order<-gsub('pole_occipital_rh','Dx116',combined$order)
combined$order<-gsub('pole_temporal_rh','Dx117',combined$order)
combined$order<-gsub('s_calcarine_rh','Dx118',combined$order)
combined$order<-gsub('s_central_rh','Dx119',combined$order)
combined$order<-gsub('s_cingul_marginalis_rh','Dx120',combined$order)
combined$order<-gsub('s_circular_insula_ant_rh','Dx121',combined$order)
combined$order<-gsub('s_circular_insula_inf_rh','Dx122',combined$order)
combined$order<-gsub('s_circular_insula_sup_rh','Dx123',combined$order)
combined$order<-gsub('s_collat_transv_ant_rh','Dx124',combined$order)
combined$order<-gsub('s_collat_transv_post_rh','Dx125',combined$order)
combined$order<-gsub('s_front_inf_rh','Dx126',combined$order)
combined$order<-gsub('s_front_middle_rh','Dx127',combined$order)
combined$order<-gsub('s_front_sup_rh','Dx128',combined$order)
combined$order<-gsub('s_interm_prim_jensen_rh','Dx129',combined$order)
combined$order<-gsub('s_intrapariet_and_p_trans_rh','Dx130',combined$order)
combined$order<-gsub('s_oc_middle_and_lunatus_rh','Dx131',combined$order)
combined$order<-gsub('s_oc_sup_and_transversal_rh','Dx132',combined$order)
combined$order<-gsub('s_occipital_ant_rh','Dx133',combined$order)
combined$order<-gsub('s_oc_temp_lat_rh','Dx134',combined$order)
combined$order<-gsub('s_oc_temp_med_and_lingual_rh','Dx135',combined$order)
combined$order<-gsub('s_orbital_lateral_rh','Dx136',combined$order)
combined$order<-gsub('s_orbital_med_olfact_rh','Dx137',combined$order)
combined$order<-gsub('s_orbital_h_shaped_rh','Dx138',combined$order)
combined$order<-gsub('s_parieto_occipital_rh','Dx139',combined$order)
combined$order<-gsub('s_pericallosal_rh','Dx140',combined$order)
combined$order<-gsub('s_postcentral_rh','Dx141',combined$order)
combined$order<-gsub('s_precentral_inf_part_rh','Dx142',combined$order)
combined$order<-gsub('s_precentral_sup_part_rh','Dx143',combined$order)
combined$order<-gsub('s_suborbital_rh','Dx144',combined$order)
combined$order<-gsub('s_subparietal_rh','Dx145',combined$order)
combined$order<-gsub('s_temporal_inf_rh','Dx146',combined$order)
combined$order<-gsub('s_temporal_sup_rh','Dx147',combined$order)
combined$order<-gsub('s_temporal_transverse_rh','Dx148',combined$order)

combined$order<-(gsub('cerebellum_cortex_lh','A01',combined$order))
combined$order<-(gsub('thalamus_proper_lh','A02',combined$order))
combined$order<-(gsub('caudate_lh','A03',combined$order))
combined$order<-(gsub('putamen_lh','A04',combined$order))
combined$order<-(gsub('pallidum_lh','A05',combined$order))
combined$order<-(gsub('brain_stem','A06',combined$order))
combined$order<-(gsub('hippocampus_lh','A07',combined$order))
combined$order<-(gsub('amygdala_lh','A08',combined$order))
combined$order<-(gsub('accumbens_area_lh','A09',combined$order))
combined$order<-(gsub('ventraldc_lh','A10',combined$order))
combined$order<-(gsub('cerebellum_cortex_rh','A11',combined$order))
combined$order<-(gsub('thalamus_proper_rh','A12',combined$order))
combined$order<-(gsub('caudate_rh','A13',combined$order))
combined$order<-(gsub('putamen_rh','A14',combined$order))
combined$order<-(gsub('pallidum_rh','A15',combined$order))
combined$order<-(gsub('hippocampus_rh','A16',combined$order))
combined$order<-(gsub('amygdala_rh','A17',combined$order))
combined$order<-(gsub('accumbens_area_rh','A18',combined$order))
combined$order<-(gsub('ventraldc_rh','A19',combined$order))

combined$order<-(gsub('precuneus_lh','D24',combined$order))
combined$order<-(gsub('precuneus_rh','D58',combined$order))
combined$order<-(gsub('bankssts_lh','D01',combined$order))
combined$order<-(gsub('caudalanteriorcingulate_lh','D02',combined$order))
combined$order<-(gsub('caudalmiddlefrontal_lh','D03',combined$order))
combined$order<-(gsub('cuneus_lh','D04',combined$order))
combined$order<-(gsub('entorhinal_lh','D05',combined$order))
combined$order<-(gsub('fusiform_lh','D06',combined$order))
combined$order<-(gsub('inferiorparietal_lh','D07',combined$order))
combined$order<-(gsub('inferiortemporal_lh','D08',combined$order))
combined$order<-(gsub('isthmuscingulate_lh','D09',combined$order))
combined$order<-(gsub('lateraloccipital_lh','D10',combined$order))
combined$order<-(gsub('lateralorbitofrontal_lh','D11',combined$order))
combined$order<-(gsub('lingual_lh','D12',combined$order))
combined$order<-(gsub('medialorbitofrontal_lh','D13',combined$order))
combined$order<-(gsub('middletemporal_lh','D14',combined$order))
combined$order<-(gsub('parahippocampal_lh','D15',combined$order))
combined$order<-(gsub('paracentral_lh','D16',combined$order))
combined$order<-(gsub('parsopercularis_lh','D17',combined$order))
combined$order<-(gsub('parsorbitalis_lh','D18',combined$order))
combined$order<-(gsub('parstriangularis_lh','D19',combined$order))
combined$order<-(gsub('pericalcarine_lh','D20',combined$order))
combined$order<-(gsub('postcentral_lh','D21',combined$order))
combined$order<-(gsub('posteriorcingulate_lh','D22',combined$order))
combined$order<-(gsub('precentral_lh','D23',combined$order))
combined$order<-(gsub('rostralanteriorcingulate_lh','D25',combined$order))
combined$order<-(gsub('rostralmiddlefrontal_lh','D26',combined$order))
combined$order<-(gsub('superiorfrontal_lh','D27',combined$order))
combined$order<-(gsub('superiorparietal_lh','D28',combined$order))
combined$order<-(gsub('superiortemporal_lh','D29',combined$order))
combined$order<-(gsub('supramarginal_lh','D30',combined$order))
combined$order<-(gsub('frontalpole_lh','D31',combined$order))
combined$order<-(gsub('temporalpole_lh','D32',combined$order))
combined$order<-(gsub('transversetemporal_lh','D33',combined$order))
combined$order<-(gsub('insula_lh','D34',combined$order))
combined$order<-(gsub('bankssts_rh','D35',combined$order))
combined$order<-(gsub('caudalanteriorcingulate_rh','D36',combined$order))
combined$order<-(gsub('caudalmiddlefrontal_rh','D37',combined$order))
combined$order<-(gsub('cuneus_rh','D38',combined$order))
combined$order<-(gsub('entorhinal_rh','D39',combined$order))
combined$order<-(gsub('fusiform_rh','D40',combined$order))
combined$order<-(gsub('inferiorparietal_rh','D41',combined$order))
combined$order<-(gsub('inferiortemporal_rh','D42',combined$order))
combined$order<-(gsub('isthmuscingulate_rh','D43',combined$order))
combined$order<-(gsub('lateraloccipital_rh','D44',combined$order))
combined$order<-(gsub('lateralorbitofrontal_rh','D45',combined$order))
combined$order<-(gsub('lingual_rh','D46',combined$order))
combined$order<-(gsub('medialorbitofrontal_rh','D47',combined$order))
combined$order<-(gsub('middletemporal_rh','D48',combined$order))
combined$order<-(gsub('parahippocampal_rh','D49',combined$order))
combined$order<-(gsub('paracentral_rh','D50',combined$order))
combined$order<-(gsub('parsopercularis_rh','D51',combined$order))
combined$order<-(gsub('parsorbitalis_rh','D52',combined$order))
combined$order<-(gsub('parstriangularis_rh','D53',combined$order))
combined$order<-(gsub('pericalcarine_rh','D54',combined$order))
combined$order<-(gsub('postcentral_rh','D55',combined$order))
combined$order<-(gsub('posteriorcingulate_rh','D56',combined$order))
combined$order<-(gsub('precentral_rh','D57',combined$order))
combined$order<-(gsub('rostralanteriorcingulate_rh','D59',combined$order))
combined$order<-(gsub('rostralmiddlefrontal_rh','D60',combined$order))
combined$order<-(gsub('superiorfrontal_rh','D61',combined$order))
combined$order<-(gsub('superiorparietal_rh','D62',combined$order))
combined$order<-(gsub('superiortemporal_rh','D63',combined$order))
combined$order<-(gsub('supramarginal_rh','D64',combined$order))
combined$order<-(gsub('frontalpole_rh','D65',combined$order))
combined$order<-(gsub('temporalpole_rh','D66',combined$order))
combined$order<-(gsub('transversetemporal_rh','D67',combined$order))
combined$order<-(gsub('insula_rh','D68',combined$order))

combined$Parcellation<-combined$Variable
combined$Parcellation<-(gsub('.*._desikan_.*.','Desikan',combined$Parcellation))
combined$Parcellation<-(gsub('.*._destrieux_.*.','Destrieux',combined$Parcellation))
combined$Parcellation<-(gsub('.*._aseg_.*.','ASEG',combined$Parcellation))

write.csv(combined,outname,row.names = FALSE)
```
